# On the impact of competing intra- and intermolecular triplet-state quenching on photobleaching and photoswitching kinetics of organic fluorophores

**DOI:** 10.1101/371443

**Authors:** Jochem H. Smit, Jasper H. M. van der Velde, Jingyi Huang, Vanessa Trauschke, Sarah S. Henrikus, Si Chen, Nikolaos Eleftheriadis, Eliza M. Warszawik, Andreas Herrmann, Thorben Cordes

**Author notes:** present address: Physical Chemistry, Department of Chemistry, Ludwig Maximilians-Universität München, Butenandtstr. 11, 81377 Munich, Germany (Vanessa Trauschke). present address: School of Chemistry, University of Wollongong, Northfields Avenue Wollongong NSW 2522, Australia.

## Abstract

While buffer cocktails remain the gold-standard for photostabilization and photoswitching of fluorescent markers, intramolecular triplet-state quenchers emerge as an alternative strategy to impart fluorophores with ‘self-healing’ or even functional properties such as photoswitching. In this contribution, we evaluated various combinations of both approaches and show that inter- and intramolecular triplet-state quenching processes compete with each other rather than being additive or even synergistic. Often intramolecular processes dominate the photophysical situation for combinations of covalently-linked and solution-based photostabilizers and photoswitching agents. In this context we identified a new function of intramolecular photostabilizers, i.e., protection of fluorophores from reversible off-switching events caused by solution-additives, which were previously misinterpreted as photobleaching. Our studies also provide practical guidance for usage of photostabilizer-dye conjugates for STORM-type super-resolution microscopy permitting the exploitation of their improved photophysics for increased spatio-temporal resolution. Finally, we provide evidence that the biochemical environment, e.g., proximity of aromatic amino-acids such as tryptophan, reduces the photostabilization efficiency of commonly used buffer cocktails. Not only have our results important implications for a deeper mechanistic understanding of self-healing dyes, but they will provide a general framework to select label positions for optimal and reproducible photostability or photoswitching kinetics.

## 1. Introduction

Non-fluorescent dark-states such as the triplet or radical states have been identified as major sources of photobleaching in (organic) fluorophores due to their relatively long lifetime and high chemical reactivity.^1–3^ Besides their tendency to directly undergo irreversible chemical reactions to non-fluorescent products (photobleaching), they can be involved in the generation of other transient and chemically reactive species. Notably, these dark-states can be used for localization-based super-resolution imaging (e.g., in techniques abbreviated STORM etc.) in case that the photoswitching kinetics can be controlled. To optimize signal strength and quality, quencher molecules, herein denoted as photostabilizers, are used to deplete triplet- or radical-states. Common quenching mechanisms for high photostability are triplet-triplet annihilation via molecular oxygen^4–6^, dexter-type triplet energy transfer (cyclooctatetraene (COT),^7,8^ diphenylhexatriene (DPHT)^9^) or photo-induced electron transfer (Trolox (TX),^10^ methylviologen (MV), ascorbic acid (AA)^2,11^). Whereas other damage pathways such as multi-photon absorption processes and bleaching from singlet states are known to occur^12^, the primary strategy for photostabilization relies on suppressing triplet- and radical-state formation as well as removal of molecular oxygen^11^ to avoid formation of reactive oxygen species (ROS).

All approaches depend upon collisions between photostabilizer and reactive fluorophore intermediates. These are generated by either addition of the photostabilizer to the imaging buffer at millimolar concentrations^2,3,10,13^ to allow diffusional intermolecular quenching (Figure 1a), or via covalent linkage, i.e., to create high local concentrations of the photostabilizer (Figure 1b).^14–20^

**Figure 1.**
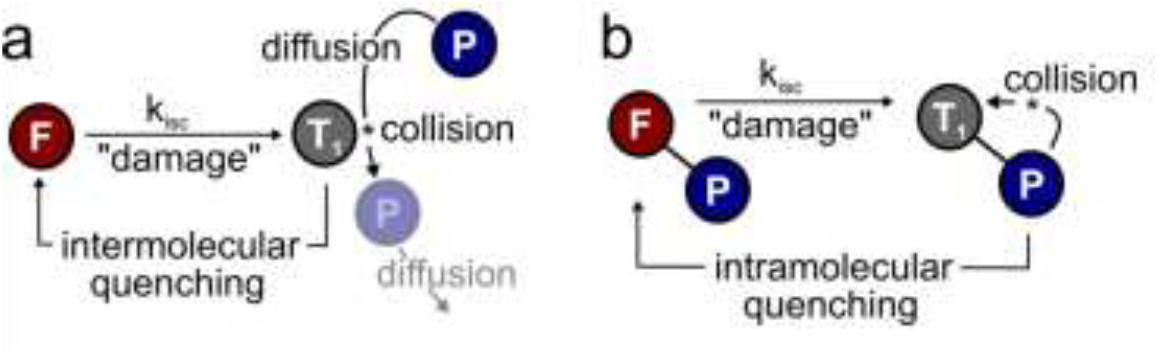
Methods to create collisions between photostabilizer and reactive fluorophore state (here triplet-state T_1_) using intermolecular diffusional quenching (a) or intramolecular processes (b). In both cases the long-lived reactive triplet-state collides with the photostabilizer to restore the fluorophore’s singlet ground state.

Currently, intermolecular photostabilization (Figure 1a) with buffer cocktails is the most effective way to stabilize various classes of organic fluorophores in different biochemical environments.^21,22,23^ This approach is widely accepted as the gold standard for photostabilization and the antioxidant TX^10^, COT^7,13^ or combinations of redox-active compounds (“ROXS”^2^) have become the additives of choice.^24^

Intramolecular photostabilization, a technique already established in the 1980s^25,26^ (Figure 1b), does not require buffer additives.^14,15^ The method can improve photophysical parameters whenever buffer additives would not be tolerated due to toxicity or interference with biological structure and function. Furthermore, intramolecular photostabilization is an effective option for live-cell imaging or in cases where buffer additives remain ineffective due to the lack of collisions between photostabilizer and an “enclosed” fluorophore. In spite of their promise, self-healing dyes have been less frequently used in part because they must be synthesized, but also because limited options for bioconjugation exist, and they have lower photostabilization efficiencies.^17,19^ Recently their applicability for super-resolution microscopy has been suggested. Previous studies by our group demonstrated that self-healing dyes allow for STED-type imaging in fixed mammalian cells with a higher number of accessible successive images due to an increased photostability.^19^ It is, however, unclear whether self-healing dyes can be used for STORM imaging via fluorophore blinking (ON/OFF switching).

Another related, yet unanswered question is whether inter- and intramolecular photostabilization can be used in combination to increase the overall photostabilization efficiency or the range of fluorophores that can be stabilized simultaneously. To address these questions, we studied the effect of commonly used solution-based healers (TX, COT) on a number of commonly-used fluorophores and their photostabilizer-conjugates, using an aromatic nitro moiety and COT. Our results suggest that either inter- or intramolecular triplet-state quenching dominate the photophysical processes of self-healing dyes, a fact that prohibits a useful combination of both approaches. These findings also prompted us to characterize the interaction of a photoswitching buffer additive used for STORM-type super-resolution imaging, tris(2-carboxyethyl)phosphine (TCEP) and cysteamine (MEA), with Cy5-COT conjugates. Once again we found that the intramolecular photostabilizer outcompetes intermolecular pathways. TCEP and MEA induce reversible off-switching, a process that can be used for superior STORM imaging, but the observed “apparent photobleaching” can also lead to misinterpretation of photostability increases whenever these compounds are used in buffer systems. These results suggest a novel role for intramolecular triplet-state quenchers; rather than inferring with photobleaching, they most likely act as protective agents for reversible off-switching reactions. Ultimately, we link our findings, i.e., a non-additive behaviour of inter- and intramolecular photostabilization effects to interactions of fluorophores with natural aromatic amino-acids and show that tryptophan residues significantly reduce diffusion-based photostabilization effects.

## 2. Results

The focus of previous work on self-healing fluorophores has been on the optimization of photostabilization efficiency,^18,20^ the development of versatile bioconjugation strategies,^19^ and to benchmark the method for different fluorophore types^19^ or environmental conditions.^18^ Here we conducted experiments in which photostabilizers were present in solution and linked to the fluorophore simultaneously (Figure 2), in an attempt to determine whether inter- and intramolecular photostablization effects can be additive or synergistic.

**Figure 2.**
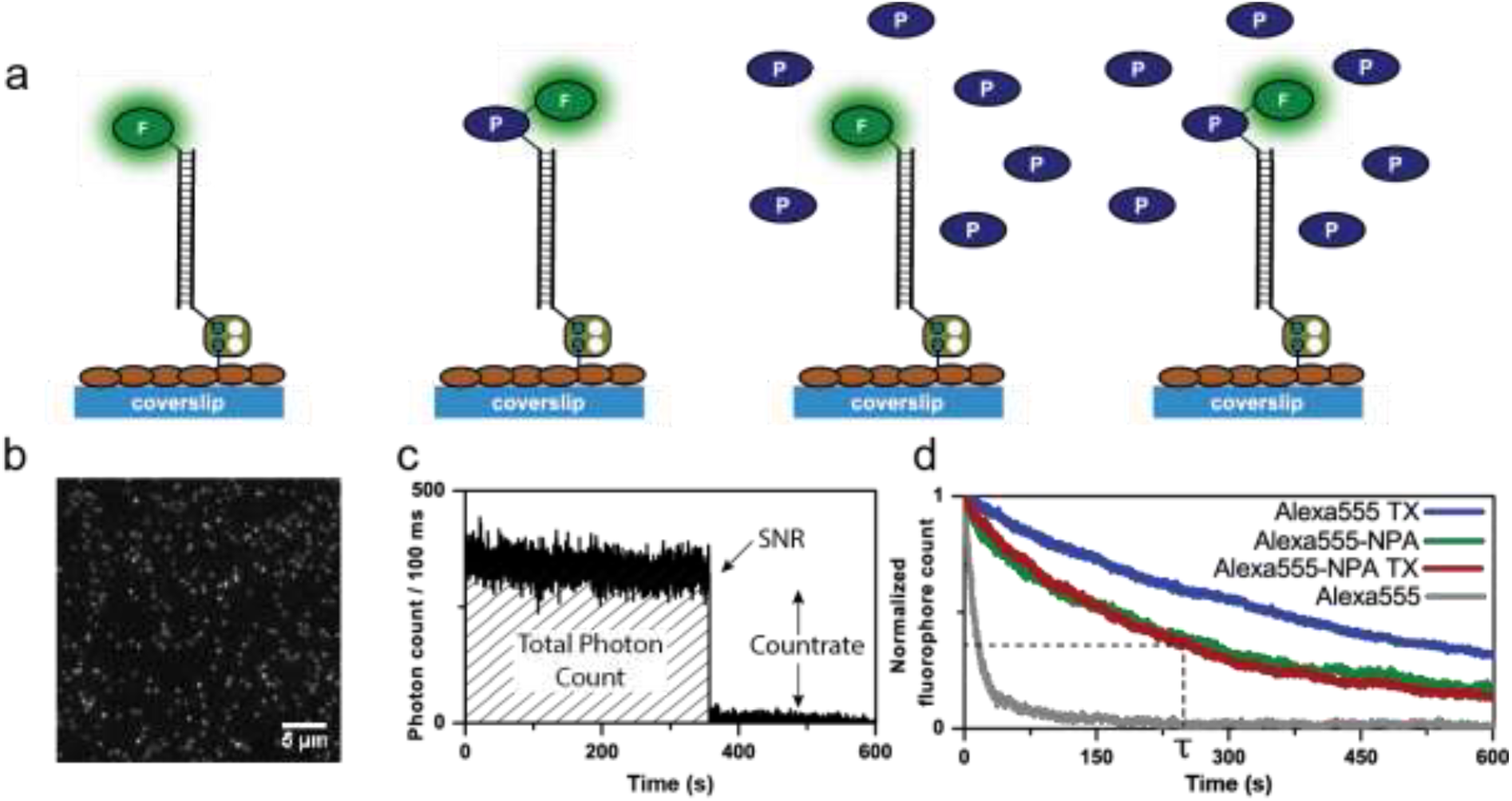
Experimental strategy and design for the characterization of the photophysical parameters of DNA-linked dyes and their photostabilizer conjugates. Different buffer systems were tested in the absence of oxygen using an enzymatic oxygen removal systems based on glucose-oxidase and catalase (GOX). a) Schematics of immobilization and experimental conditions. Representative single-molecule data sets of b) TIRF images of individual fluorophores, c) fluorescent transients of individual fluorophores which were used to extract photophysical parameters. d) Normalized plots of fluorophore counts in each time frame providing quantitative information on photobleaching kinetics.

To this end, fluorescent dyes and their photostabilizer-dye conjugates were immobilized on a streptavidin-functionalized coverslip via double-stranded DNA-scaffolds as described previously^19^ (Figure 2a). Total-internal-reflection fluorescence (TIRF) microscopy was employed to quantify the photophysical properties of different permutations of inter- and intramolecular photostabilization (Figure 2a) in a deoxygenated environment. Quantitative photophysical parameters were obtained from single emitters (Figure 2b-d) identified in TIRF images. Detailed analysis of their fluorescent transients allowed us to extract the count-rate and signal-to-noise ratio (SNR). The number of fluorescent emitters over time (Figure 2d) was fitted to an exponential decay which gave the mean photobleaching lifetime. From this, the total number of photons emitted before photobleaching can be calculated by multiplication of the photobleaching lifetime and the average count-rate.

We tested three frequently used fluorophores, namely ATTO647N, Alexa555 and Cy5, and compared them to their respective photostabilizer-dye conjugates (nitrophenyl alanine (NPA)-ATTO647N, NPA-Alexa555, NPA-Cy5, Cy5-COT Figure 3, Figure S1-S20). The performance of the dyes was evaluated under either deoxygenated conditions (glucose oxidase based oxygen scavenging, GOX), or deoxygenated conditions with additional photostabilizers: 2 mM aged TX, (antioxidant) or 2mM COT (triplet-state quencher). The GOX-buffer system in Figure 2 and 3 contains 200 μM TCEP, which was added for enzyme stability.

**Figure 3.**
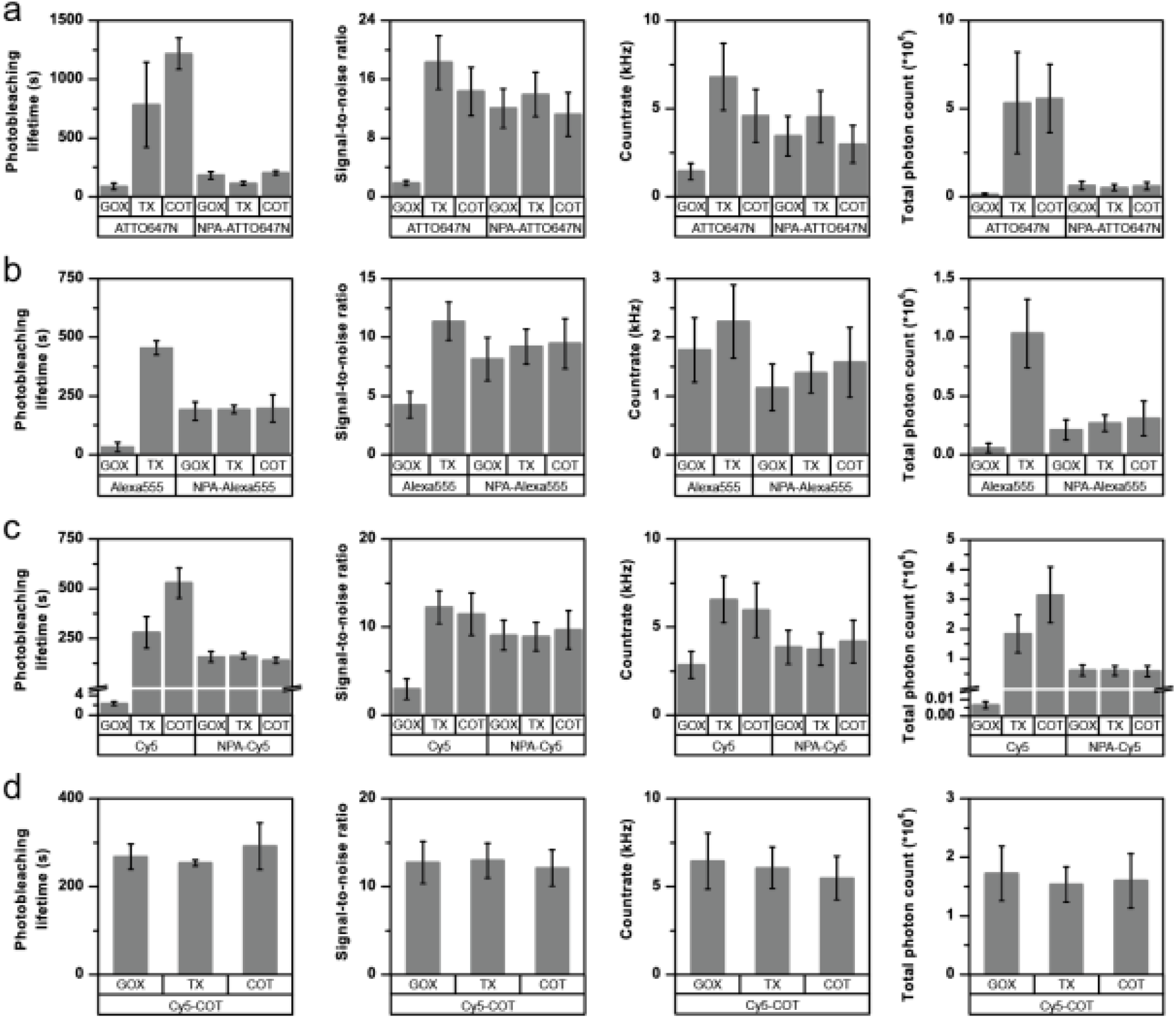
Photophysical parameters of a) ATTO647N and NPA-ATTO647N b) Alexa555 and NPA-Alexa555 c) Cy5 and NPA-Cy5 d) Cy5-COT under deoxygenated conditions without the addition of solution-based photostabilizers (GOX) and in the presence of 2 mM TX or 2 mM COT. The glucose-oxidase catalase enzyme cocktail contains 200 μM of TCEP. Photobleaching lifetime numbers were obtained by fitting several photobleaching decay curves and averaging the obtained time constants. Error bars are standard deviation of 3 independent measurements. Signal-to-noise ratio and count-rate were determined from individual fluorescent transients for ≥ 200 molecules. The total photon count is the product of the photobleaching lifetime and count-rate. The error bar shows the standard deviation of repeats on two different days. A detailed description is given in the “Material and Methods” section.

In agreement with previously published results^15,16,19^, we found an improvement of photophysical parameters in the presence of solution additives or when comparing unmodified fluorophores with their NPA- or COT-conjugates as shown in Figure 3. This included increased photobleaching lifetimes, signal-to-noise ratio, count-rates and total number of collected photons, and can be attributed to the depopulation of the dark triplet-state (Figure 3a-c and Figure S1-S17).

### Inter- and intramolecular quenching processes compete with each other

Strikingly, for all fluorophores and stabilizer combinations, we observed no improvement of photophysical properties upon combining inter- and intramolecular photostabilization. Instead, the photobleaching lifetimes did not increase beyond values observed for NPA- or COT-conjugates. For example, when comparing the photobleaching lifetimes of the carbopyronine fluorophore ATTO647N under different conditions (Figure 3a), there was a clear increase between the parent dye in GOX buffer (oxygen removal only) and TX-based or COT-based buffers. A smaller increase was also observed for covalent linkage of NPA to the dye (NPA-ATTO647N, GOX) with respect to the parent compound (ATTO647N, GOX). However, the combination of both inter- and intramolecular stabilization, e.g., NPA-ATTO647N with either TX or COT as buffer additives, did not result in an increased photostability with respect to NPA-ATTO647N.

Similar results were obtained for Alexa555 (Figure 3b) and Cy5 (Figure 3c) in combination with the covalent stabilizer NPA (Figure 3a-c) or COT (Figure 3d). It can thus be concluded that the photostabilizing effect observed for the two approaches individually is not additive. Instead, the photobleaching lifetimes were similar to that of the photostabilizer conjugates and were not affected further by stabilizers in solution. Besides these general trends, there were subtle differences in the photophysics of NPA-fluorophores in GOX buffer and TX/COT-containing buffers. For NPA-ATTO647N and NPA-Cy5, addition of TX removed faster signal fluctuations (Figure S4 vs. S5 & Figure S15 vs. S16) whereas addition of COT induced short up-blinks of unknown origin (Figure S4 vs. S6). Also, the addition of TX/COT increased the overall count-rate for NPA-based dyes (Figure S4 vs. S5 & Figure S9 vs. S11) but not for intramolecular-linked COT that fully shields the fluorophore from other photostabilizers in solution (Figures S18-S20). This could be rationalized as being due to the presence of a transient dark state in the NPA-based quenching pathway, which does not contribute to photobleaching, and which was removed by TX and COT in solution, thus improving the other photophysical parameters.

These findings suggest that in the competition for the triplet-state, the covalently bound stabilizer effectively outcompetes the solution based stabilizers, i.e., it quenches the triplet-state faster. This can be understood in view of the fact that in the intramolecular case, the photostabilizer is present in much higher local concentrations as compared to the solution additives. However, faster quenching of the triplet-state (which is difficult to observe directly in single-molecule experiments) did not result in higher levels of photostability.

These conclusions are supported by our earlier findings that two covalently tethered stabilizers (e.g., a fluorophore with both TX and NPA attached) did not show improved performance beyond that of the individual stabilizers.^16^ In this case too, photostabilization was dominated by one of the two stabilizers – presumably the one which quenches the triplet faster.

In previous work, we also investigated the importance of stabilizer-fluorophore arrangement and linker length for self-healing processes.^18^ Different linking chemistries or orientations of the photostabilizer resulted in different photostabilities. Our interpretation of the data in Figure 3 is further corroborated by experiments in which covalently linked NPA was arranged differently in space with respect to Cy5 (Figure S29). The nitrophenyl group used as a photostabilizer was either linked directly to the fluorophore through an amino-acid scaffold on the 3’-end of a DNA strand (NPA-Cy5), or was attached on its own to the 5’-end of the complementary DNA strand (NPAA) such that upon annealing it was brought into proximity with the fluorophore directly attached to the other strand (Cy5-NPAA); Figure S29. The photobleaching lifetimes of NPA-Cy5 and Cy5-NPAA were found to be different (207 ± 25 s and 647 ± 241 s, respectively). Significantly, annealing the DNA strand NPA-Cy5 with the NPAA strand to place Cy5 in proximity of two stabilizers moieties (Figure S29) did not lead a strong additive of synergistic effect on the photobleaching lifetime of Cy5. While there is a slight difference between the spatial arrangement of NPAA and Cy5 in the absence and presence of the amino-acid NPA-moiety, this data suggests again that the photophysics is dominated by the closest photostabilizer unit.

A simple combination of inter- and intramolecular approaches (or multiple intramolecular stabilizers) was found not to be beneficial for photostability. This finding allowed us to draw further mechanistic conclusions which we discuss in detail below.

### Photoswitching competes with intramolecular triplet-state quenching and photobleaching

Our results have direct implications for the use of self-healing dyes in localization-based super-resolution microscopy, because these techniques require controlled on/off-switching of single fluorophores via solution additives.^27^ In STORM, compounds such as mercaptoehtylamine (2-aminoethane-1-thiol, MEA), ß–mercaptoethanol (2-sulfanylethan-1-ol, ß-ME) or TCEP are used to from a meta-stable dark state. Typically, this dark state can be recovered to the singlet ground state by illuminating with UV light, thus achieving reversible photoswitching. The effect of reducing compounds and particularly that of thiols has been extensively studied.^27–33^ A suitable concentration of photoswitching additive allows for the control of the on/off-duty cycle needed for optimal temporal separation of fluorophore emission in STORM.^21,30,34,35^

To study the interplay of intramolecular photostabilizers with photoswitching compounds in solution and explore whether self-healing dyes could be useful for STORM imaging, we investigated the influence of the photoswitching agent TCEP and MEA on the photophysical properties of Cy5 and Cy5-COT. In STORM microscopy, the off-switching events are either (i) photo-induced or (ii) proceed via an equilibrium step. As an example, the reversible addition reaction between TCEP and Cy5 occurs with an equilibrium constant of 0.91 mM^-1^.^27^ With this information at hand, we first examined the impact of 200 μM of TCEP as a buffer additive on the photophysics of Cy5 and Cy5-COT. TCEP was typically present in our buffers at this concentration to maintain GOX enzyme stability (Figure 3). According to the equilibrium constant, 16% of the Cy5 molecules should be quenched. Strikingly, however, the presence of 200 μM TCEP reduced the signal duration by 11-fold, which was a much larger effect than expected (Figure 4a, Cy5 ± 200 μM TCEP).

**Figure 4.**
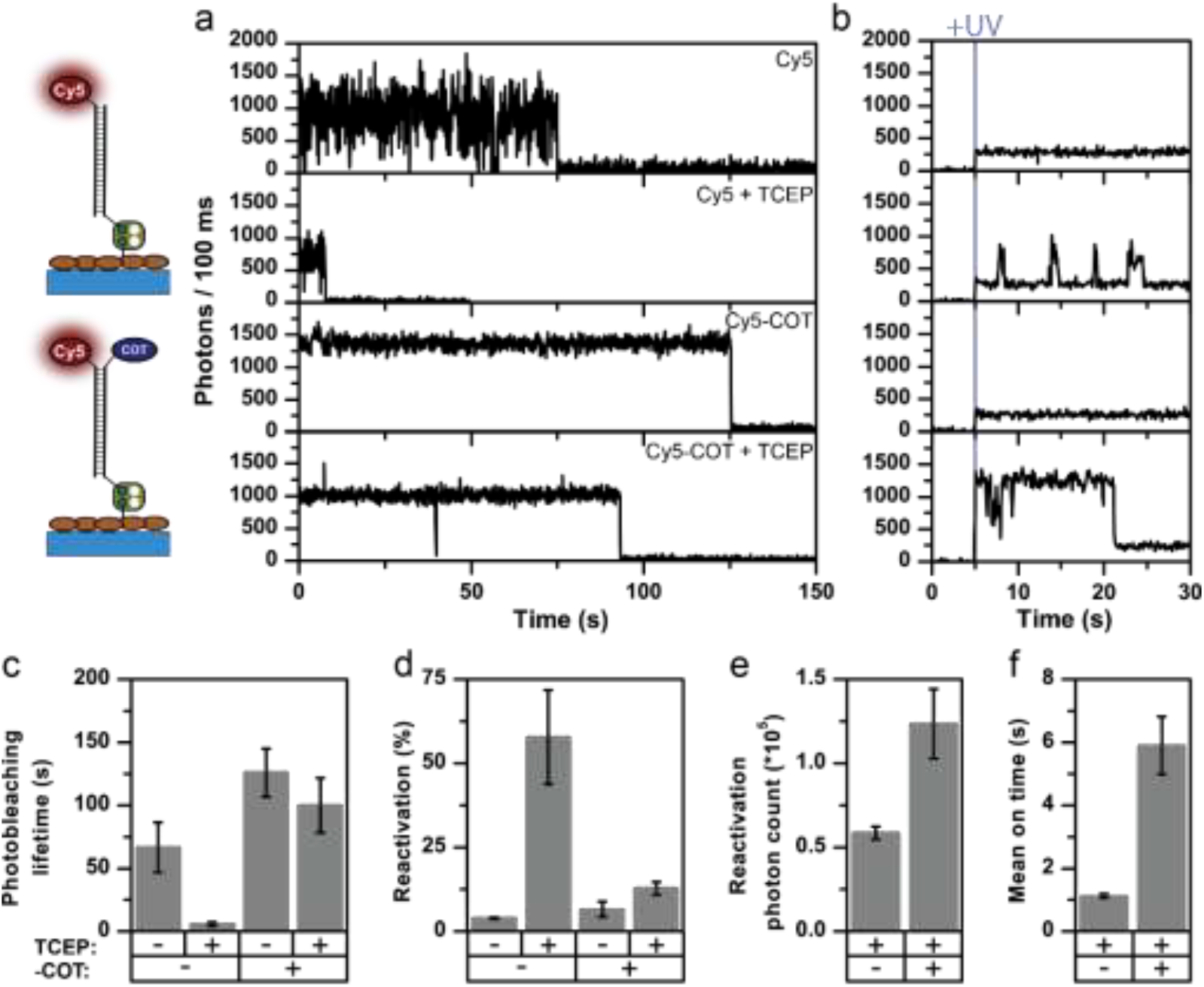
Comparison of Cy5 parent fluorophore and stabilized Cy5-COT in the absence and presence of 200 μM TCEP. Imaging performed under deoxygenated conditions (GOX) with excitation at 637 nm, 0.4 kW cm^-2^. a) Representative fluorescence transients. b) Representative fluorescence transient under UV illumination (375 nm, 0.1 kW cm^-2^). c) Photobleaching lifetimes. d) Reactivation yield during UV illumination. e) Mean photon count per activated fluorophore and e) mean on-time during activation. Error bars are either standard deviations (c, d) of repeats on 3 different days or SEM (e, f). A detailed description is given in the “Material and Methods” section.

This result cannot be explained solely by equilibrium quenching. We postulate that the faster signal loss observed in the presence of TCEP involves a photo-induced off-switching. Apparently, TCEP uses the Cy5 triplet-state for off-switching even at very low concentrations similar to the mechanism of thiols.^28,33,36^ Experiments with Cy5-COT support this notion of photo-induced off-switching via triplet-state quenching since the impact of TCEP was reduced for Cy5-COT (Figure 4a). In Cy5-COT, both TCEP and COT apparently compete for the triplet-state and the intramolecular quencher successfully outcompetes TCEP due to its high local concentration.

This conclusion was further corroborated by photoactivation experiments using 375 nm UV light (Figure 4a, second column). In our experiments, both real photobleaching and off-switching result in signal loss, which we relate to “apparent photobleaching”. After apparent photobleaching, the UV laser was turned on to monitor potential fluorescence recovery and thus to disentangle the reversible off-switching contribution caused by TCEP from actual irreversible photobleaching. As expected, when TCEP is omitted, the fluorophore reactivation upon applying UV light is minimal and considered to be background signal. This indicates that the observed signal loss in the absence of TCEP is indeed irreversible photobleaching. In contrast, a significant fraction, >50% of the fluorophores, can be reactivated in the presence of TCEP (Figure 4d). When reactivating Cy5-COT, the activation yield in the presence of TCEP is much lower than with Cy5 (Figure 4d), but always significantly higher than in TCEP absence (Figure 4d). Consequently, the mean relative on-times and photon counts after photoactivation were higher for Cy5-COT. Longer on-times of Cy5-COT compared to Cy5 are attributed to prevention of TCEP-induced darkening but not improved photophysics of Cy5 in the on-state. The blinking kinetics of the parent fluorophore Cy5 were also characterized by confocal microscopy (Figure S30) to determine the origin of fast blinking. By calculating the autocorrelation function of fluorescent transients, the *cis*-*trans* and triplet-related off times of Cy5 blinking were measured. We found that these values were not influenced by TCEP, suggesting that TCEP caused only the appearance of photoactivatable long off states, but did not induce short-timescale blinking.

Similar effects were observed for PET-based photostabilizer-dye conjugates NPA-Cy5 and TX-Cy5 conjugates in the presence of TCEP (Figure S33). Also the presence of these distinct photostabilizers protects the fluorophore against reversible off-switching, resulting in longer apparent photobleaching times and on-duration after UV-activation. Overall, however, lower reactivation yields are observed for both conjugates. In the presence of 5 mM MEA, (Figure S34) a reducer frequently used for STORM imaging, the reactivation yields are generally higher, suggesting under these conditions a dark state is able to efficiently form.

Since we and other groups in the field^1,2,13,14,20^ use reducing agents such as TCEP, MEA and ß-ME as additives in imaging buffers when characterizing the photophysics of (self-healing) dyes, we determined the apparent photostability values of our dyes in the presence and absence of 200 μM TCEP for representative buffer conditions, i.e., comparable to those in Figure 3. While there was only a small impact on Alexa555 and no impact on ATTO647N (Figure S31), the apparent bleaching lifetime of Cy5 was drastically shortened by TCEP (Figure 4a/c). This suggests that the previously reported photostability increases of Cy5 fluorophores upon linkage to stabilizers represent an overestimation.^2,13–16,19,20^ Here, the dye-standard without the linked stabilizer showed artificially low photon numbers that likely represent the kinetics of reversible Cy5-off-switching by TCEP/MEA/ß-ME, but not those of photobleaching. Fast signal loss via reversible off-switching has been described earlier by Aitken et al.^11^ for TCEP, ß-ME, DTT and other reducing agents when added at 10 mM concentrations. Lower concentrations of e.g., 200 μM TCEP had, however, not been characterized until now, but seem to demand similar attention. Our findings suggest that reducing compounds should not be used when the photophysics of fluorophores are to be characterized, and reveal a new role for intramolecular photostabilizers, namely the protection of fluorophores from reversible photoswitching reactions caused by reducing agents. The implications of our findings for design criteria and mechanistic understanding of self-healing dyes and their use in super-resolution microscopy will be discussed in detail below.

### Self-healing dyes for STORM-type super-resolution microscopy

We previously demonstrated that self-healing dyes are useful for STED-microscopy because their superior photostability increases the number of available successive images.^19^ In STORM, imaging is a more complex interplay of photoswitching kinetics (to temporally separate the labels) and photostability. The available photon number N in the on-state relates to a lower bound of the localization error which can be approximated by

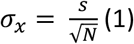

with s being the standard deviation of the fitted Gaussian function.^37,38^ Intramolecular photostabilizers vastly change the photoswitching kinetics, efficiency of fluorophore reactivation, available on-times and number of photons N during one on period as shown in Figure 4. Consequently, they also impact the localization accuracy, which is linked to the achievable (theoretical) resolution and the number of emitters that can be separated in time. The latter indirectly links to spatial resolution via the Nyquist-Shannon sampling theorem^39,40^ but also to temporal resolution of the technique.^41^ From this perspective, the results shown in Figure 4 indicate that Cy5-COT is not well suited for use in STORM at low TCEP concentrations and low laser powers due to a poor on-off duty cycle, i.e., long on-times with low count-rate.

To improve the situation, we systematically tested variations of laser power and TCEP concentration in experiments shown in Figure 5. We also varied the UV activation conditions, and as expected from previous work,^27^ increasing UV activation power was found to shorten the off-times (Figure S32a). In our experiments in Figure 5, we used 375 nm activation light at 90 W/cm^2^. In practical applications these tuneable off-times can be used to optimize the on-off duty ratio according to the needs of the specific experiment.

**Figure 5.**
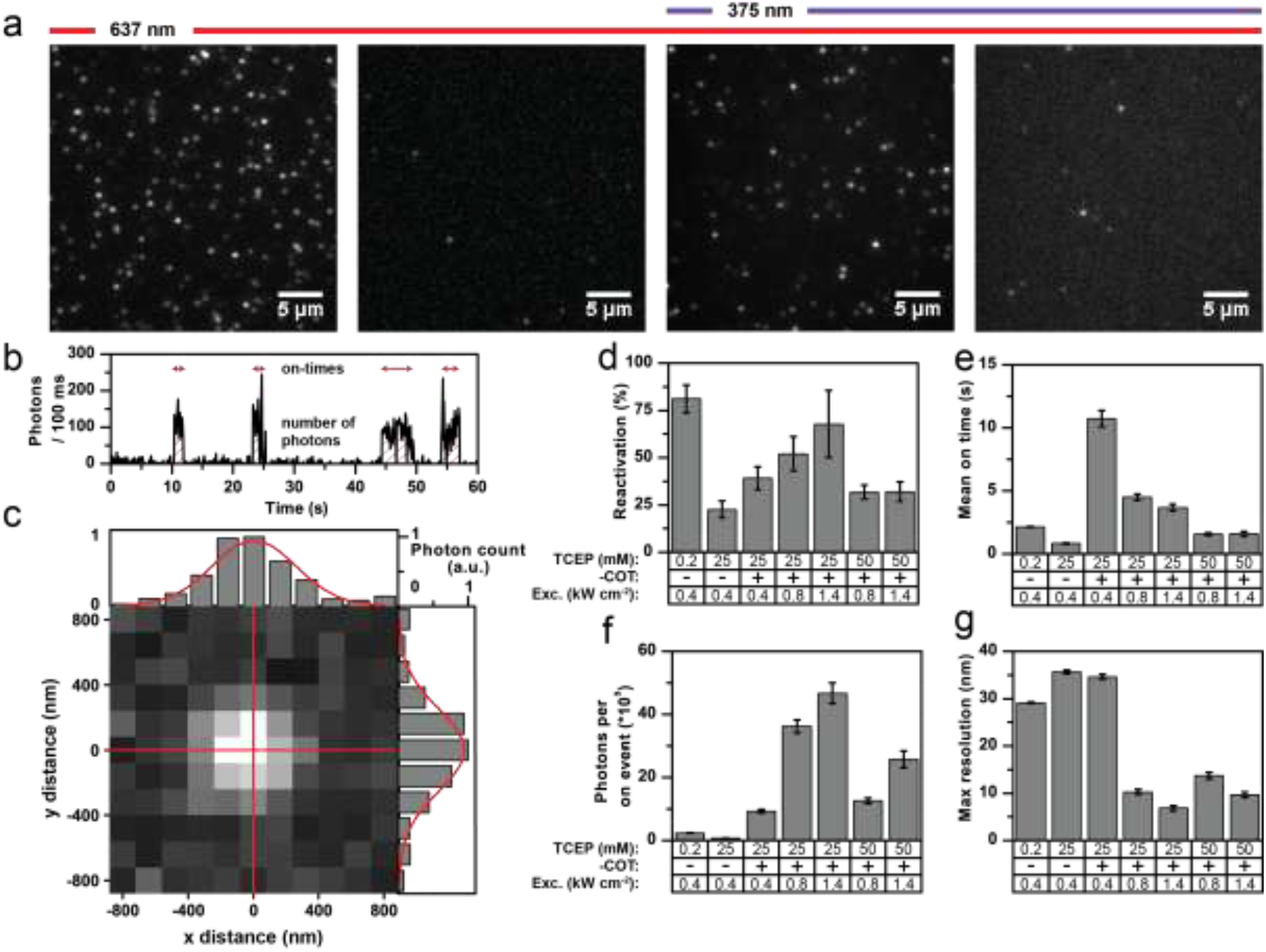
Characterization of Cy5 and Cy5-COT for application in STORM-type super-resolution microscopy. Imaging was done under deoxygenated conditions (GOX) with different TCEP concentrations. a) TIRF images of (1) frames 1-10, (2) last frame of image acquisition, (3) frames 1-500 averaged, and (4) last frame of subsequent photoactivation acquisition. b) A representative fluorescent transient showing the photoactivation of Cy5 in the presence of 200 uM TCEP. c) A single photoactivated emitter along with x and y cross-section showing the result of 2D Gaussian fitting (red). The standard deviation of the Gaussian fit was found to be 201 nm. (d-g) Parameters describing the performance of the dyes for super-resolution. Error bars are standard deviation of 2-3 different days (d) or SEM for each different condition. See main text and materials and methods for further details.

Figure 5a/b show the details of how we extracted quantitative parameters to evaluate the suitability of Cy5-COT for STORM. Figure 5a shows an initial field of view of individual Cy5 or Cy5-COT molecules that are continually irradiated with 637 nm excitation light to induce photobleaching or photoswitching. An algorithm counts the number of molecules in the initial field of view until only a few to none remain. Then the 375 nm excitation laser was switched on and all subsequently reactivated molecules were counted. The reactivation yield provided in Figure 5d is the ratio between the number of reactivated molecules and the initial molecule count. Single activation events were found by thresholding the fluorescent transients, where short off blinks (< 400 ms) were ignored. For every activation event, the on-time and the total number of photons was calculated; the mean values for the different conditions are shown in Figure 5e and 5f, respectively. The localization error was calculated according to equation (1) and the resulting values were multiplied by 2.35 to give the theoretically obtainable resolution.^32^ The mean values per condition are a shown in Figure 5g.

Our analysis shows that Cy5-COT can become a superior dye for STORM microscopy compared to Cy5; the optimal settings were found to be 1.4 kW/cm^2^ of laser excitation and 25 mM of TCEP. In the ideal case, the reactivation yield (Figure 5d) of both Cy5 and Cy5-COT can be tuned to be >60%, where the parent Cy5 performs slightly better than the photostabilizer-dye conjugate (but only at low TCEP concentrations). Consistent with expectations, the on-duration decreased with higher laser power (Figure 5e). Interestingly, the photons per on-period decreased for parent Cy5 but increased for Cy5-COT with higher laser power (Figure 5f). This unique feature of Cy5-COT enables optimum photon-yields to be reached and with it maximum obtainable resolution. As mentioned before, the on-off duty ratio, which needs to be small to allow temporal fluorophore separation in STORM-imaging, can be adjusted further by decreasing UV excitation or increasing the TCEP concentration (Figure 5e). In these experiments, Cy5-COT was clearly revealed to be a superior dye for STORM imaging. The number of reactivation events per fluorophore (Cy5 vs. Cy5-COT) suggests that the gain likely comes from a 6-fold lower number of activation events for Cy5-COT compared to Cy5 (Figure S32b), and from an overall higher total photon yield (Figure S32c).

When comparing the Cy5-COT conjugates (Figure 4,5,S32) with redox-based NPA-Cy5 and TX-Cy5 (Figure S33), TCEP reveals poor reactivation yields and on-times. In case of using MEA as photoswitching agent of Cy5, a better reactivation yields is achieved for NPA-Cy5 and TX-Cy5 compared to Cy5-COT. NPA-Cy5 in combination with MEA has also extremely short apparent photobleaching lifetimes and thus represents a good candidate for low on/off-duty ratios.

### Proximal tryptophan residues can reduce the photostabilization effects of solution-based healers

In the previous sections, we have shown that covalently coupled triplet-state quenchers effectively outcompete solution-based additives for photostabilization and –switching, whenever the underlying quenching mechanism involves the triplet-state. These findings prompted us to investigate whether natural aromatic amino-acids might have a similar effect, since they are known to quench both singlet- and triplet-states of xanthene and oxazine dyes, via a PET quenching mechanism.^42–44^ Our hypothesis was that aromatic amino-acids in close proximity to the fluorophore might reduce photostabilization by intermolecular stabilizers in a similar manner to covalently-linked NPA or COT moieties. Such a mechanism might explain the large variations in the size of the photostabilization effects observed with identical buffer cocktails and dye pairs on different targets such as proteins and oligonucleotides.^2,45,46^

To test our hypothesis, we prepared DNA-conjugates of Cy5 and ATTO647N with a single tryptophan residue (Figure 6a). The photostability of the compounds, i.e., fluorophore and its corresponding tryptophan conjugate, were investigated in the presence and absence of Trolox. As a result of conclusion drawn in the previous section, all experiments were now performed without TCEP in the GOX buffer, and photobleaching lifetimes for Cy5 and ATTO647N of ^~^100 s and ^~^50 s, respectively, were observed (Figure 6a,b).

**Figure 6.**
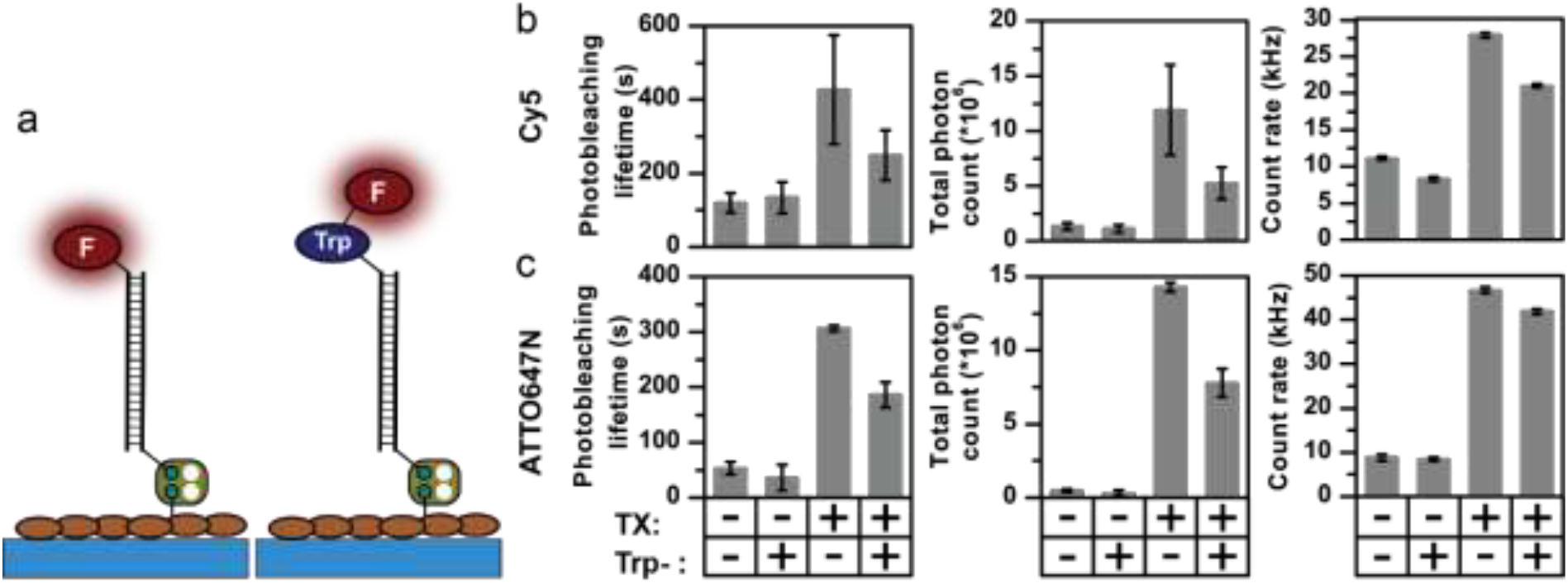
Photophycial properties of (b) Cy5 and (c) ATTO647N, showing the influence of a covalently coupled tryptophan residue in the absence and presence of 2 mM TX. All measurements were done in the absence of oxygen with 637 nm laser power of 800 W cm^-2^ Error bars show standard deviation of repeated experiments as described in the “Material and Methods” section.

Linking a tryptophan residue covalently to the fluorophore did not drastically influence the photophysics of the two dyes (Figures S21-S28). A small decrease in Cy5 brightness was observed, in line with reported relative quantum yields in literature,^47^ likely due to singlet-state quenching. Since the total photon output is not increased upon tryptophan conjugation, it appears that it is not able to quench both the Cy5 or ATTO647N triplet states. However, the photostabilizing effect of TX in solution was significantly affected. Whereas the addition of 2 mM TX increased the photobleaching lifetime of Cy5 to ^~^450 s and that of ATTO647N to ^~^300 s, both tryptophan conjugates showed an ^~^2-fold lower stability and thus also lower total photon counts (Figure 6b). Identical experiments to those shown in Figure 6 were also performed with phenylalanine and tyrosine; both had no notable influence on the observed photostability (Figure S31). These experiments are suggestive of a key role for tryptophan in the photostabilization of organic dyes (as also suggest previously by others), and support the notion that the biochemical environment of a fluorophore has a strong influence on its photophysical properties.

Interestingly, tryptophan-dependent effects on photostability were not observed when examining Cy5 and Trp-Cy5 in combination with 2 mM COT in the buffer (data not shown), suggesting that COT might compete more efficiently with Trp for triplet-state quenching. An alternative explanation is that Trp interacts with fluorophore radical states which are only formed during the solution-based quenching pathway with TX but not in the case of COT. This might be the mechanistic reason for the observation that COT is often a more potent photostabilizer for proteins single-molecule fluorescence studies.^13^ The notion that tryptophan interacts with fluorophore radical states might have implications for super-resolution microscopy (in particular STORM), which relies on the formation of a stable dark state to obtain the desired blinking kinetics using TCEP, MEA or ß-ME.^48^ The presence of tryptophan in close proximity to the fluorophore label could explain observed differences in blinking kinetics when coupled to different targets and antibodies.

## 4. Discussion & Outlook

The work presented in this study revealed the underlying mechanisms of competing inter- and intramolecular pathways for quenching of triplet-states in organic fluorophores (Figure 7). We found that (i) inter- and intramolecular processes compete with each other rather than being additive or even synergistic (Figure 7a-c). This is caused by a high local concentration and faster triplet-state quenching by intramolecular processes either via NPA (Figure 7a, PET) or direct triplet-state quenching using COT. (ii) Inter- and intramolecular pathways have different photostabilization efficiencies; although the first step of the intramolecular pathway is fast, long-lasting and reactive subsequent intermediates prevent effective overall photostabilization. (iii) A new verified function of intramolecular photostabilizers is the protection^67^ of fluorophores from reversible off-switching reactions caused by solution-additives (Figure 7a vs. 7c), which can be misinterpreted as irreversible photobleaching. (iv) Self-healing dyes can be used for STORM-type super-resolution microscopy and provide improved photon-yields and potentially higher spatial resolution. (v) Proximity of tryptophan residues significantly reduces the photostabilization efficiency of commonly used buffer cocktails. These findings have significant implications for the design and characterization of self-healing dyes, as well as for their applications.

**Figure 7.**
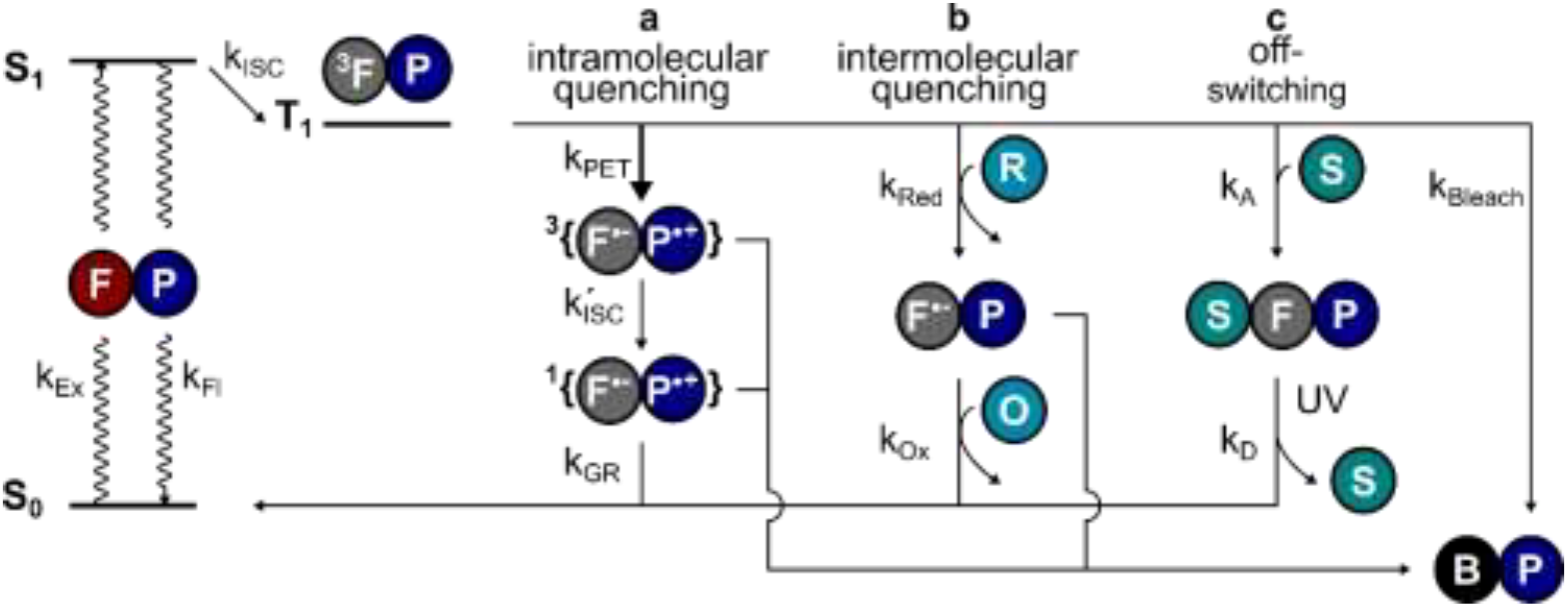
Schematic view of the main pathways through which photostabilization occurs by depletion of the triplet-state. a) In self-healing dyes, the first step of Intramolecular quenching k_PET_ outcompetes the other pathways due to a higher localized concentration of the photostabilizer P. However, bleaching from subsequent radical intermediates can still limit the photostability. b) Intermolecular quenching (ROXS) takes places in two steps with reduction (k_Red_) followed by oxidation (k_Ox_) to recover the fluorophore ground state. c) Off-switching for STORM-type super resolution is facilitated by thiols or other reducing agents that form a dark state via the triplet-state route. This pathway can be misinterpreted as apparent photobleaching since the fluorophore can be recovered through illumination with UV light.

A remaining unanswered question is whether buffer additives could successfully support the photostabilizer and with that positively influence fluorophore photophysics.^49^ Assuming that intramolecular healing is faster at triplet-state quenching than solution based methods, it would have be expected that intramolecular healing outperforms intermolecular healing. This prediction, however, is not supported in the present work by the observed photobleaching lifetimes and total photon counts. One possible explanation might be that the limiting factor is the stability of the photostabilizing group. We have previously reported photophysical events^15,50^ in which the fluorescent signal abruptly transitions from a stable signal into parent fluorophore photophysics. In reality, these events are rare and can thus not account for the observed discrepancy, unless photochemical destruction of the photostabilizer also results in immediate bleaching of the parent fluorophore. Another possible explanation is that additional photobleaching pathways exist and these are facilitated by the intramolecular quenching mechanism (Figure 7a). The mechanism of self-healing might possibly include reactive states such as charge-separated states (NPA) or triplet-states (COT), which can contribute to photobleaching and thereby limit the maximal photobleaching lifetime. This is supported by recent work from the Cosa lab^51^, where it was shown that efficient charge recombination in self-healing dyes requires intersystem crossing (ISC) of the biradical (k’ISC, Figure 7a). This slow ISC, and therefore long-lasting biradical, might be the cause for differences in photostability between different fluorophore-stabilizer geometries (Figure S29), because a longer linker may allow faster ISC and therefore faster recovery of the fluorophore ground state.

Our findings also have various implications for the rational design of self-healing dyes with photoswitching kinetics tailored for super-resolution imaging and might explain why the same dye has completely different photoswitching in different biochemical sourroundings, e.g., antibody vs. DNA or protein. They should also be taken into account when quantitative determinations of photostabilization effects are being made, since TCEP, MEA or ß-ME are often found in buffers to ensure sample stability. The thiol compounds MEA and ß-ME are known to reversible switch dyes off through a similar mechanism as proposed here for TCEP.^28,33^ We and others have shown that some dyes, particularly cyanine-based ones, have a strong tendency for fast but reversible signal loss, which can be totally unrelated to photobleaching. In the presence of TCEP (and likely also for MEA or ß-ME), it was impossible with our methods to distinguish between irreversible photobleaching and reversible photoswitching. Consequently, these compounds have to be omitted for quantitative determination of photobleaching lifetimes. We found these effects to be most pronounced for Cy5 dyes but almost negligible for Alexa555 and ATTO647N. These observations undermine the conclusions drawn in studies reporting enhancement factors for self-healing cyanine dyes.^2,13–16,19,20^ In these earlier investigations, competition between reversible off-switching and triplet-state quenching of the photostabilizer most likely would have varied, hindering a quantitative determination of the degree of photostabilization, and impeding a true understanding of the mechanism of self-healing. And finally, self-healing dyes can be used for STORM-microscopy as demonstrated for Cy5-COT, and probably also in a straightforward fashion for techniques that do not rely on photoswitching, e.g., PAINT or DNA PAINT.^52,53^

Recently, the connection between the reduction of triplet-state lifetime and increases in photostability has been reported.^20^ However, the authors of the study did not take into account reversible off-switching by triplet-reactive reagents such as MEA or ß-ME, suggesting that the rapid triplet depletion that they observed was not directly translated into photostability, but may have originated from a protection of the fluorophore from photoswitching processes. The conjugates developed in that work^20^, however, still represent significantly improved dyes for both live-cell and *in vitro* studies, because the triplet-state was effectively quenched, the probes are bright and photostable, and no longer act as photosensitizers, thus preventing cell damage through phototoxicity.^54–56^ Moreover, it has been shown that fluorophores in close proximity to specific amino-acid residues can generate reactive and long-lived species,^57–59^ which can lead to artefacts in, for example, single-molecule FRET measurements on proteins.^60^ The overall role of oxygen and its competition with intramolecular healers still has to be clarified and was not taken into account in our work here.

In conclusion, we suggest an improved protocol for determining photostabilization of self-healing dyes, where triplet-reactive solution additives are omitted. When using self-healing dyes, photostabilizing buffer additives have little or no effect, and the presence of photoswitching reagents can lead to the misinterpretation of reversible signal loss (off-switching) as apparent photobleaching. We also show that the biochemical surroundings of “normal” fluorophores have to be carefully chosen to ensure no harmful residues such as tryptophan might impact stabilization or switching efficiencies. Taken together our results serve as a guide on how to rationally optimize the parameters needed to use the higher photon output of dyes in general and that of self-healing ones for STORM-type super-resolution microscopy. Finally, our study suggests a novel function of intramolecular triplet-state quenchers as a protector of fluorophores from reversible photoswitching reactions.

## Materials and Methods

### Synthesis

All chemical compounds were purchased from commercial suppliers and used without further purification. A Varian 400MHz was used to record ^1^H-NMR and ^13^C-NMR spectra.

The dsDNA-fluorophore samples with the labels ATTO647N, NPA-ATTO647, Alexa555, NPA-Alexa555, Cy5, and NPA-Cy5, were synthesized and prepared as previously described^15,19^. Cy5-COT was prepared by labelling a biotinylated ssDNA-NH2 strand named P2 (Biotin-5’-CGT CCA GAG GAA TCG AAT ATT A-3’-NH_2_) with NHS-COT; see SI for synthetic details using a modification of a procedure from the literature^14^. The resulting oligonucleotide was characterized by MALDI-TOF mass spectrometry (Figure S46) using a ABI Voyager DE-PRO MALDI-TOF (delayed extraction reflector) Biospectrometry Workstation mass spectrometer. Hybridizing this strand to ssDNA-Cy5 (Cy5-5’-TAA TAT TCG ATT CCT CTG GAC G-3’) gave the Cy5-COT sample. The ssDNA-Trp-Cy5/ATTO647N constructs were synthesized by subsequently coupling tryptophan (Fmoc-Trp-NHS) and the different dyes to a ssDNA-NH_2_ named P1 (NH_2_-5’-TAA TAT TCG ATT CCT CTG GAC G-3’), according to previously established procedures^19^. The resulting oligonucleotide was characterized by ESI-MS (Figure S47-48). The final ssDNA product was hybridized to a biotin-modified complementary DNA strand (unmodified P2) to allow immobilization.

### Microscopy and sample preparation

Immobilization of single fluorophores was achieved using a dsDNA scaffold as previously described.^19^ In summary, Lab-Tek 8-well 750 uL chambered cover slides (Nunc/VWR, The Netherlands) were cleaned by incubating for 2 minutes with 0.1 M HF and washed with PBS buffer. The chambers were incubated with a mixture of 5 mg/mL BSA and 1 mg/mL BSA-Biotin in PBS overnight at 4 °C. After rinsing with PBS buffer, the chambers were incubated with 0.2 mg/mL streptavidin in PBS for 10 min and subsequently rinsed with PBS buffer. The fluorophores were immobilized via biotin-streptavidin interaction by incubating with a 50-100 pM solution leading to a surface coverage allowing the imaging of single emitters. All experiments were carried out at room temperature. Deoxygenated conditions were achieved by using an oxygen-scavenging system (PBS, 10%w/v glucose and 10% v/v glycerol, 50 μg/mL glucose oxidase, 100-200 μg/mL catalase and 0.2 mM Tris(2-carboxyethyl)phosphine hydrochloride (TCEP) unless stated otherwise).

Widefield TIRF imaging was performed on an inverted microscope (Olympus IX-71, UPlanSApo x100 NA 1.49 Objective, Olympus, Germany) in a similar manner as described before.^15^ Images were collected with a back-illuminated electron multiplying charge-coupled device camera (512×512 pixel, C9100-13, Hammamatsu, Japan) with matching filters and optics. They were stored using either MetaMorph or MicroManager.^61^ Whereas all the data shown in Figures S1-S21 were obtained on an Olympus IX-71 setup, that for Figures S22-S28 and the related main text Figure 5 were obtained on an Olympus IX-83 setup. The photon numbers and count-rates on the latter setup were slightly different which we attribute to differences in effective excitation power, collection efficiency, detection efficiency and count-to-photons calibration.

Confocal microscopy was performed on a home-built setup as described previously.^15^

### TIRF – data analysis

Individual fluorophores were detected in TIRF movies using a fixed threshold and discoidal averaging filter. The number of emitters as a function of time was fitted to a mono-exponential decay to obtain the mean photobleaching lifetime. For a typical experiment, 5 movies were recorded of a given condition, which was repeated on 3 different days. Fluorescent transients were extracted from the data by selecting a 3×3 pixel area (pixel size 160 nm) around the emitter and plotting the resulting mean fluorescence intensity in time. These fluorescent transients were then processed in home-written software to extract other photophysical parameters such as signal-to-noise ratio and count-rate. First, the photobleaching step and long off-blinks were identified using an implementation of a change-point algorithm.^62,63^ The signal after photobleaching was identified as the background and was subtracted from the fluorescent transient. For fluorophores which do not bleach within the duration of the acquisition, the average background value of neighbouring emitters was used. Long off-blinks (> several frames) were excluded from the analysis. The AD counts from the camera were converted to photon counts by calibrating the camera using the relation between variance and mean of a poissonian process. Finally, signal-to-noise ratio and count-rate were calculated from single transients, where signal-to-noise ratio was given as the ratio between the mean and standard deviation of the signal.

### STORM – data analysis

Reactivation yields were given by the ratio between the number of activated molecules and the number of initially present molecules in a given field of view. First, in a new area on the microscope coverslip, the fluorophores were switched off or photobleached by excitation with 637 nm. The first 20 frames were averaged in time to remove fast off-blinking and a peak finding algorithm was used to count the number of molecules. Any remaining molecules (typically 0-10 molecules) were removed from the initial molecule count. For conditions with high TCEP concentration (≥ 25 mM for Cy5, ≥ 50 mM), the initial off-switching was fast or most emitters were already darkened before imaging. For these conditions, the sample was imaged in conditions without TCEP present and the number of fluorophores was determined for multiple field of views. The molecule distribution was assumed to be homogenous and therefore the average initial molecule count can be approximated. After this first step, the 375 nm (90 W cm^-2^) activation laser was turned on and another movie was acquired. To ensure all activated molecules were found in time, the maximum pixel intensity was projected along the time axis and a peak finding algorithm was applied on the resulting image giving the number of activated molecules.

Fluorescent transients were extracted (3×3 pixel area as above) and background corrected for every activated molecule. The transients were thresholded by Otsu’s method^64,65^ to find activation events. Short off-blinks (< 400 ms) were excluded from analysis. For every activation event, the on-time and number of photons was determined. A single emitter was fitted to a 2D Gaussian to find its standard deviation, s. Then, using this value of s and equation (1), the localization error can be calculated for every on-event. Finally, these values were multiplied by 2.35 to obtain the theoretical maximum resolution and then averaged to give a single maximum resolution for the different imaging conditions.

Autocorrelation analysis was performed on fluorescent transients obtained from confocal microscopy (Figure S30). The transients were fitted to bi-exponential decays to obtain on and off times.^66^

## ACKNOWLEGDEMENTS

This work was financed by an ERC Starting Grant (ERC-STG 638536 – SM-IMPORT to T.C.) and an ERC Advanced Grant (ERC-ADG 694610 – SUPRABIOTICS to A.H.). J.H.M.v.d.V acknowledges a PhD stipend from the Ubbo-Emmius funds (University of Groningen). T.C. was supported by the Center of Nanoscience Munich (CeNS), Deutsche Forschungsgemeinschaft within GRK2062, LMUexcellent and the Center for integrated protein science Munich (CiPSM). We thank D. Griffith, V. Glembockyte and P. Tinnefeld for thoroughly reading the manuscript and for their critical and useful comments.

## Supplementary Information

### 1. Additional data

**Figure S1:**
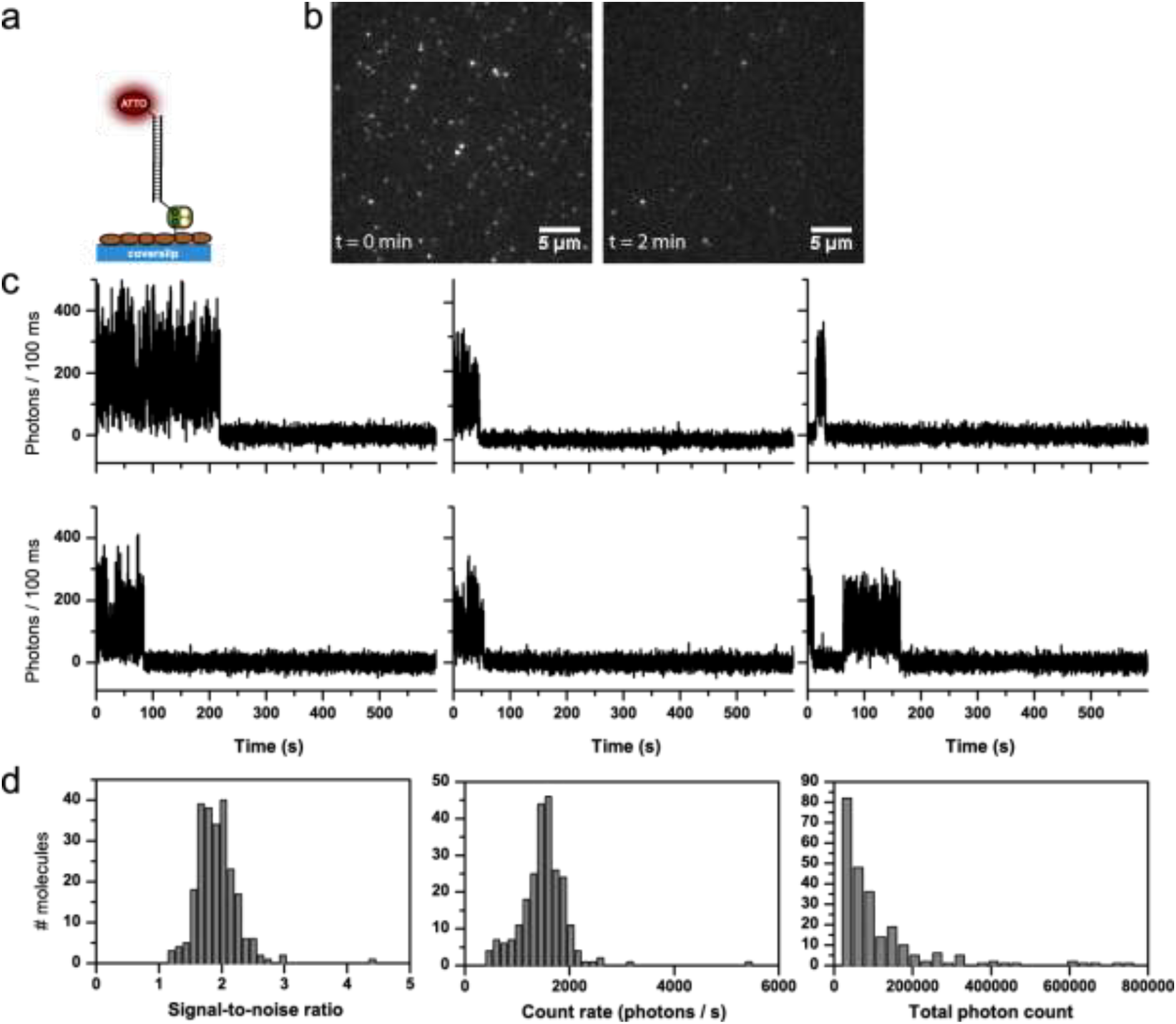
Detailed photophysical characterization of ATTO647N (GOX buffer, no photostabilizer in solution). a) Schematic representation of dye conjugation on DNA immobilized on a BSA/BSA-Biotin surface. b) TIRF images at different points in time, brightness and contrast 2848 to 35868 (AD counts). c) Representative time traces. d) Histograms of signal-to-noise ratio, count-rate and total photon count from individual time traces.

**Figure S2:**
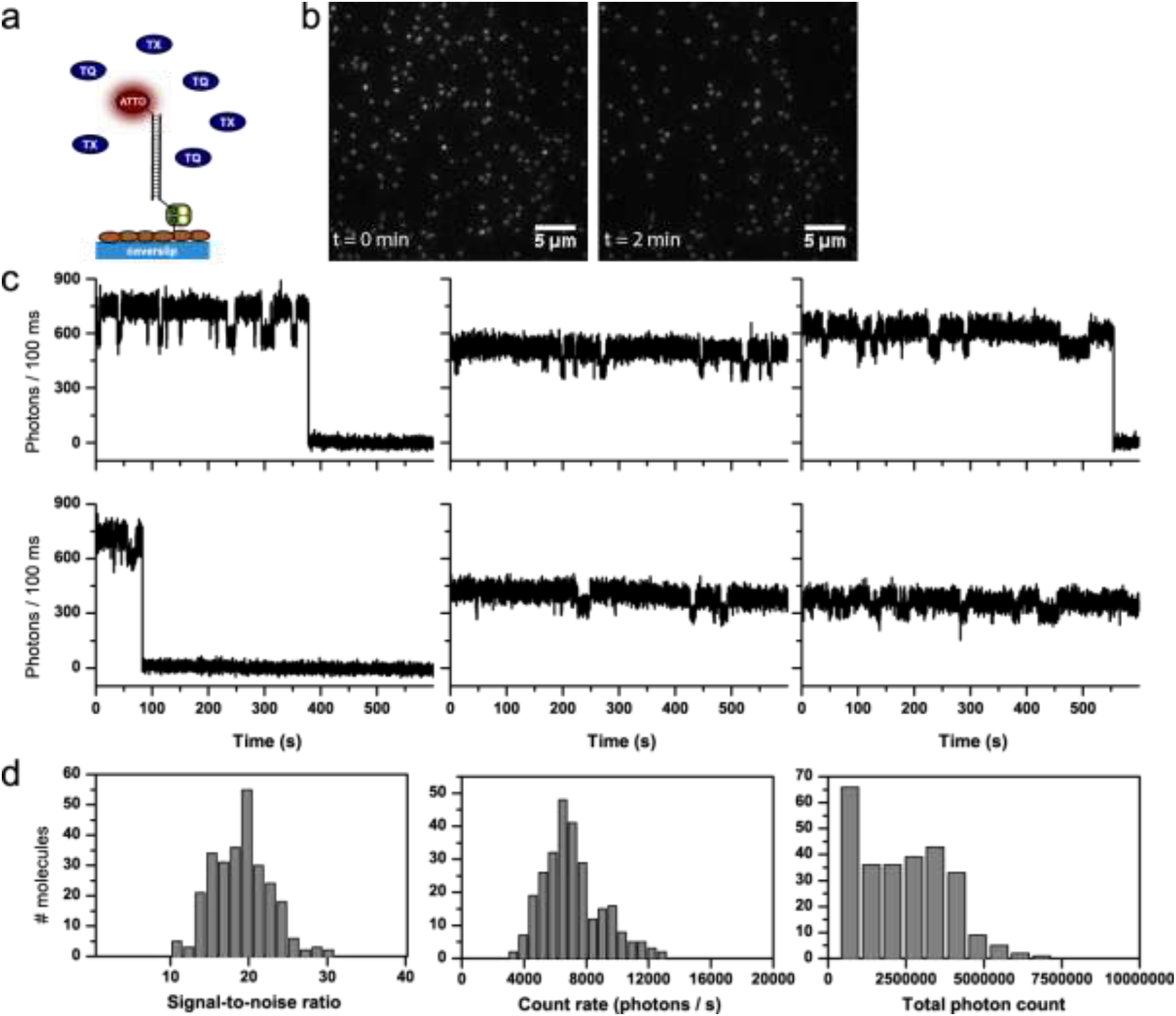
Detailed photophysical characterization of ATTO647N (buffer with 2 mM TX). a) Schematic representation of dye conjugation on DNA immobilized on a BSA/BSA-Biotin surface. b) TIRF images at different points in time, brightness and contrast 3494 to 54418 (AD counts). c) Representative time traces. d) Histograms of signal-to-noise ratio, count-rate and total photon count from individual time traces.

**Figure S3:**
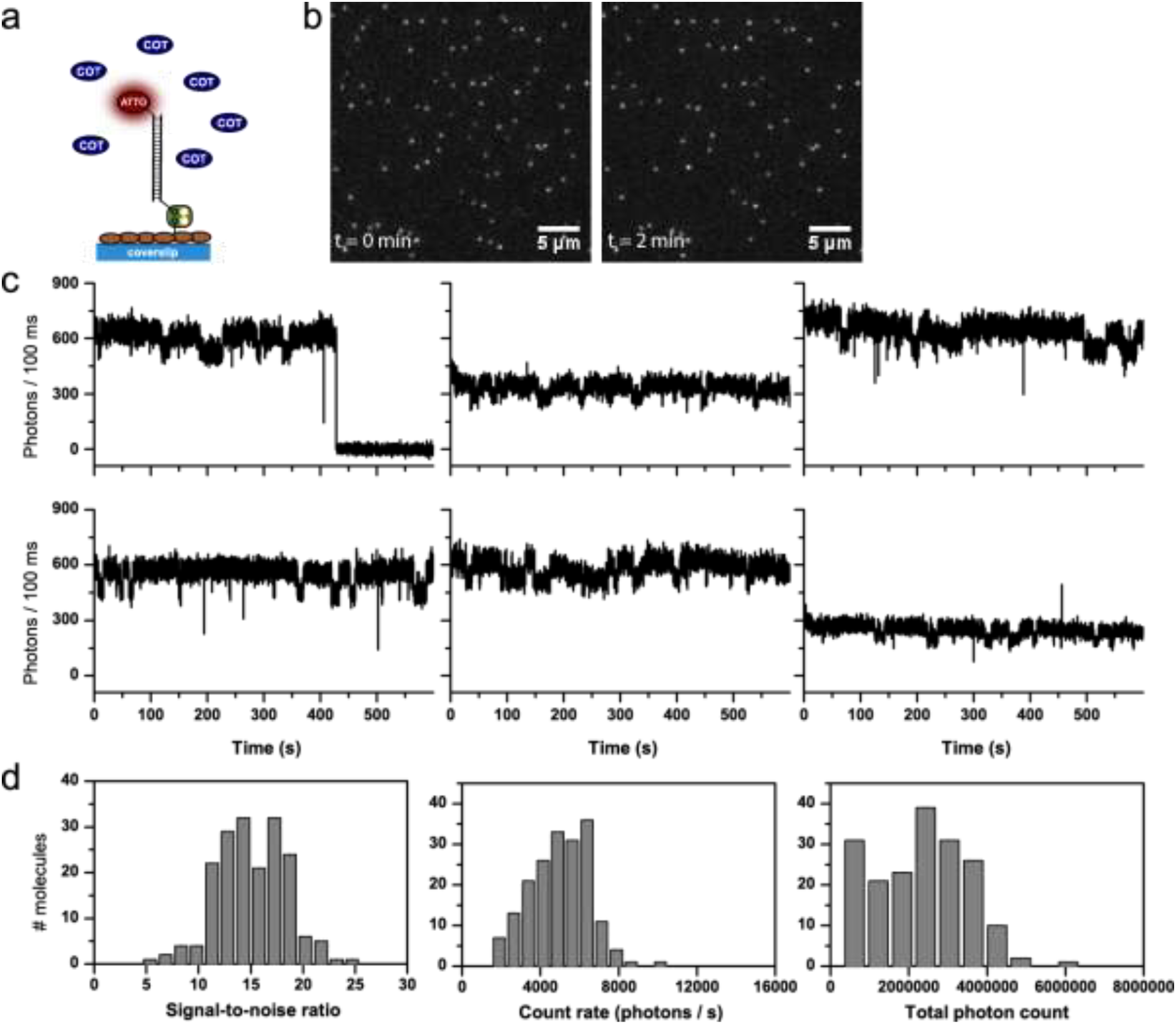
Detailed photophysical characterization of ATTO647N (buffer with 2 mM COT). a) Schematic representation of dye conjugation on DNA immobilized on a BSA/BSA-Biotin surface. b) TIRF images at different points in time, brightness and contrast 2394 to 16983 (AD counts). c) Representative time traces. d) Histograms of signal-to-noise ratio, count-rate and total photon count from individual time traces.

**Figure S4:**
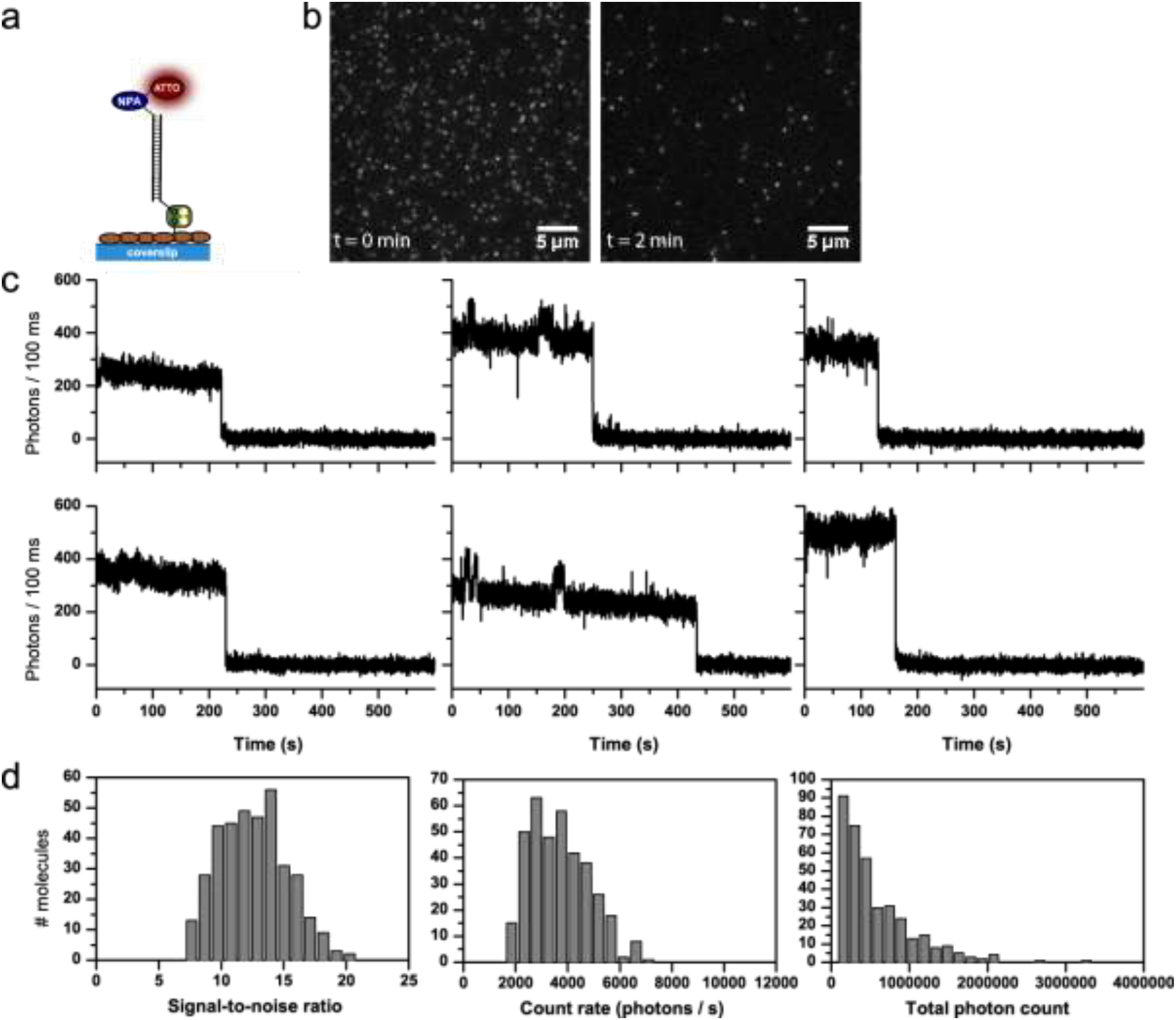
Detailed photophysical characterization of NPA-ATTO647N (GOX buffer, no photostabilizer in solution). a) Schematic representation of dye conjugation on DNA immobilized on a BSA/BSA-Biotin surface. b) TIRF images at different points in time, brightness and contrast 3037 to 24420 (AD counts). c) Representative time traces. d) Histograms of signal-to-noise ratio, count-rate and total photon count from individual time traces.

**Figure S5:**
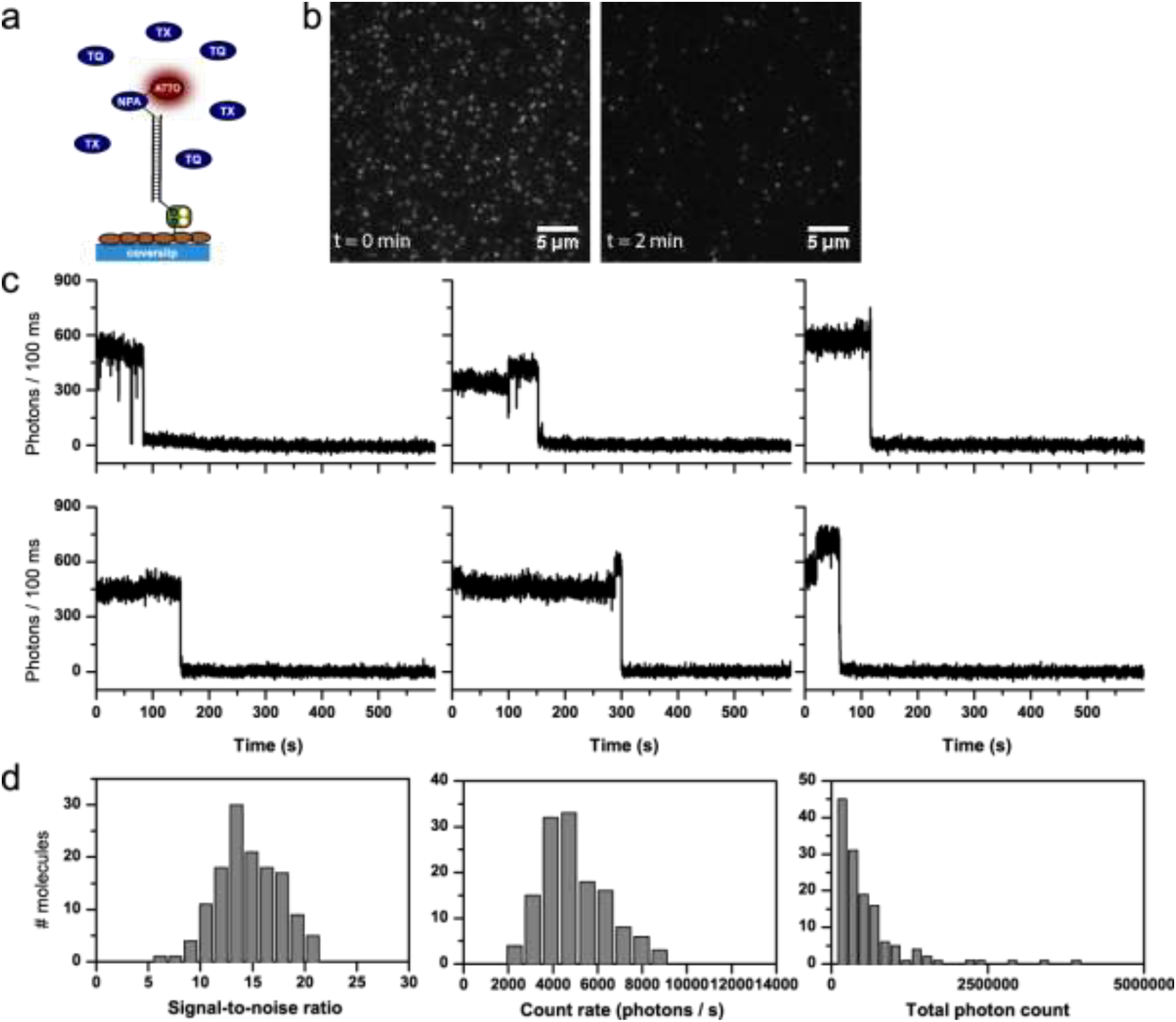
Detailed photophysical characterization of NPA-ATTO647N (buffer with 2 mM TX). a) Schematic representation of photostabilizer-dye conjugates on DNA immobilized on a BSA/BSA-Biotin surface. b) TIRF images at different points in time, brightness and contrast 3177 to 25515 (AD counts). c) Representative time traces. d) Histograms of signal-to-noise ratio, count-rate and total photon count from individual time traces.

**Figure S6:**
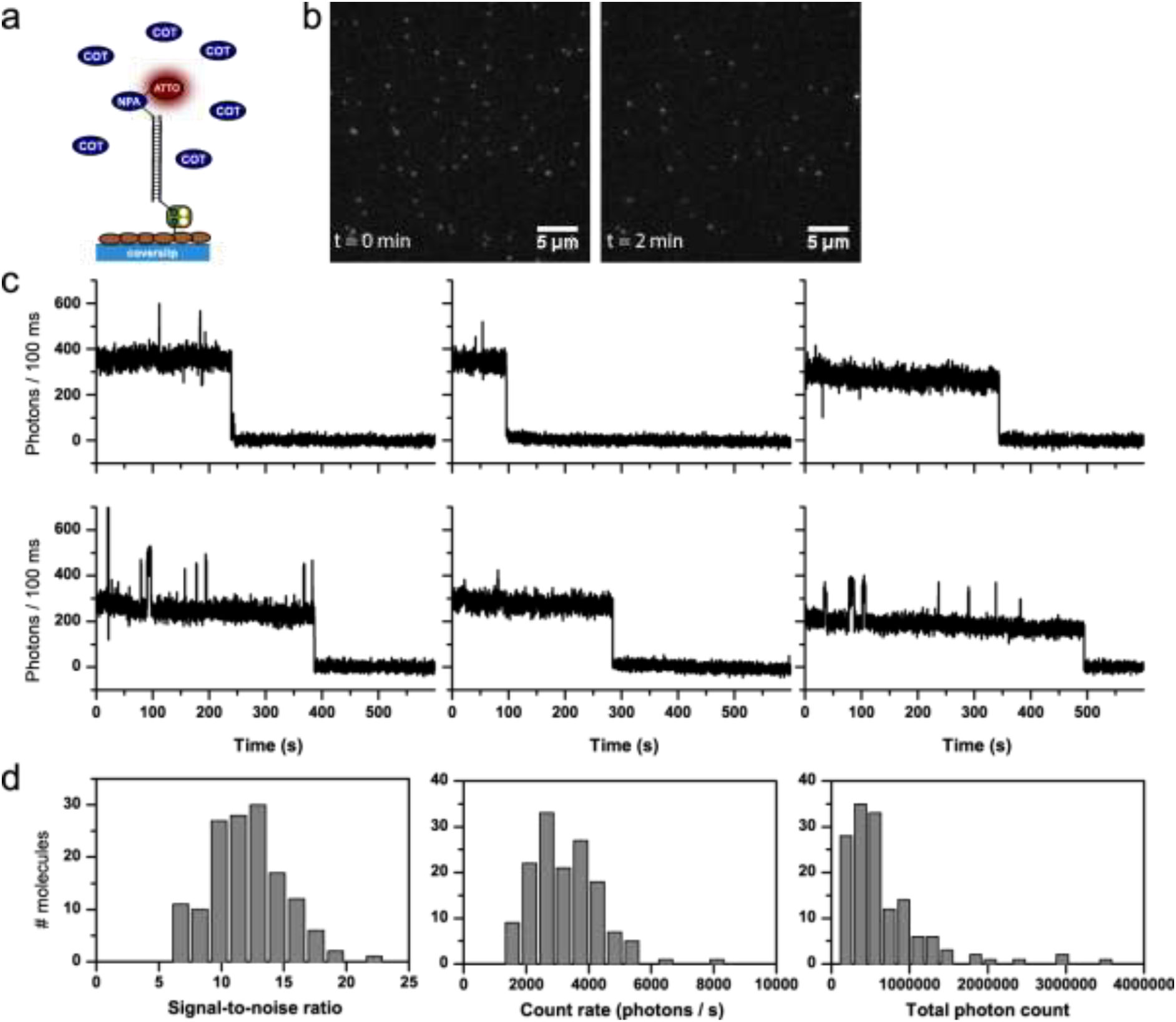
Detailed photophysical characterization of NPA-ATTO647N (buffer with 2 mM COT). a) Schematic representation of photostabilizer-dye conjugates on DNA immobilized on a BSA/BSA-Biotin surface. b) TIRF images at different points in time, brightness and contrast 2699 to 25823 (AD counts). c) Representative time traces. d) Histograms of signal-to-noise ratio, count-rate and total photon count from individual time traces.

**Figure S7:**
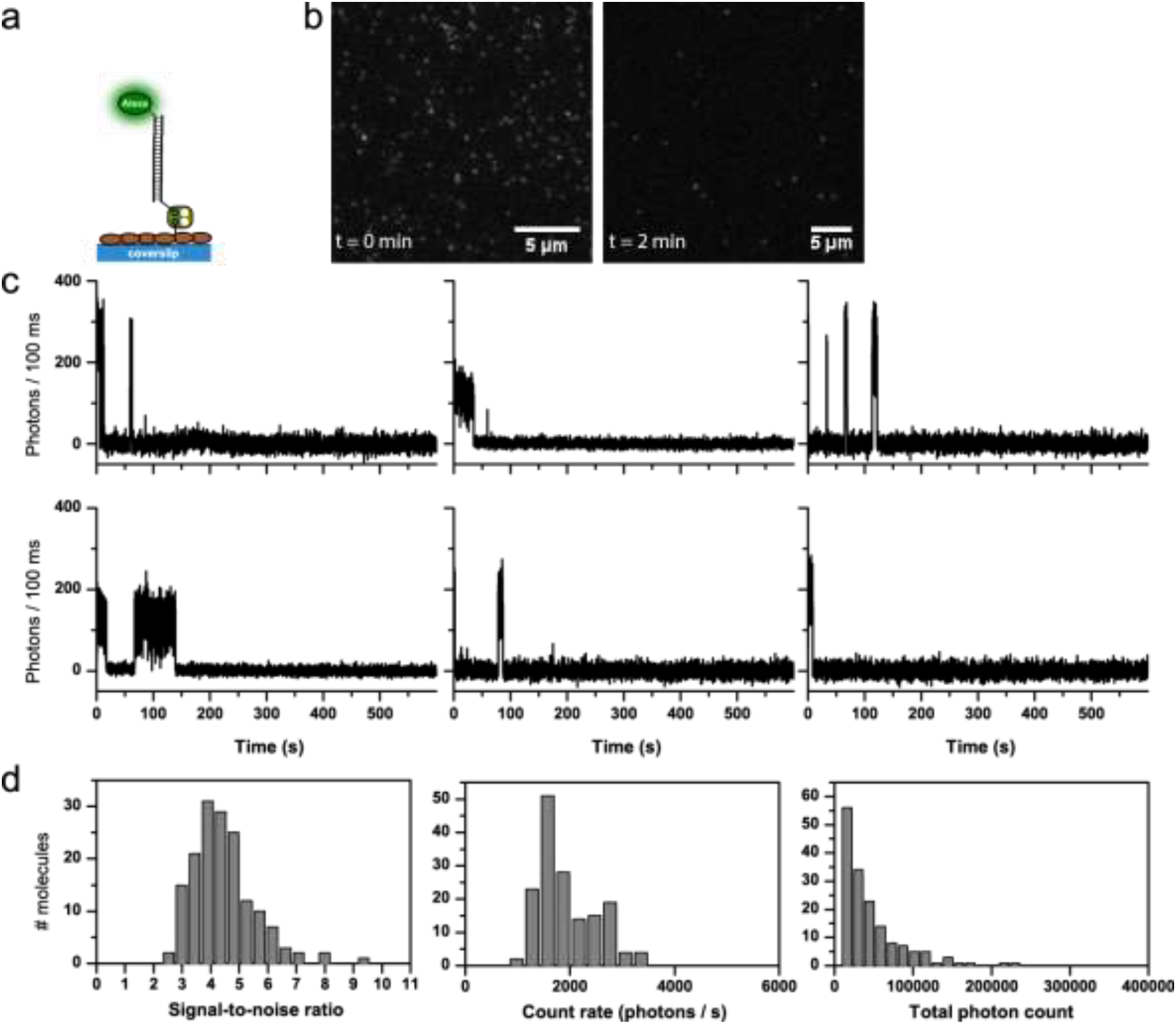
Detailed photophysical characterization of Alexa555 (GOX buffer, no photostabilizer in solution). a) Schematic representation of dye conjugation on DNA immobilized on a BSA/BSA-Biotin surface. b) TIRF images at different points in time, brightness and contrast 1876 to 14482 (AD counts). c) Representative time traces. d) Histograms of signal-to-noise ratio, count-rate and total photon count from individual time traces.

**Figure S8:**
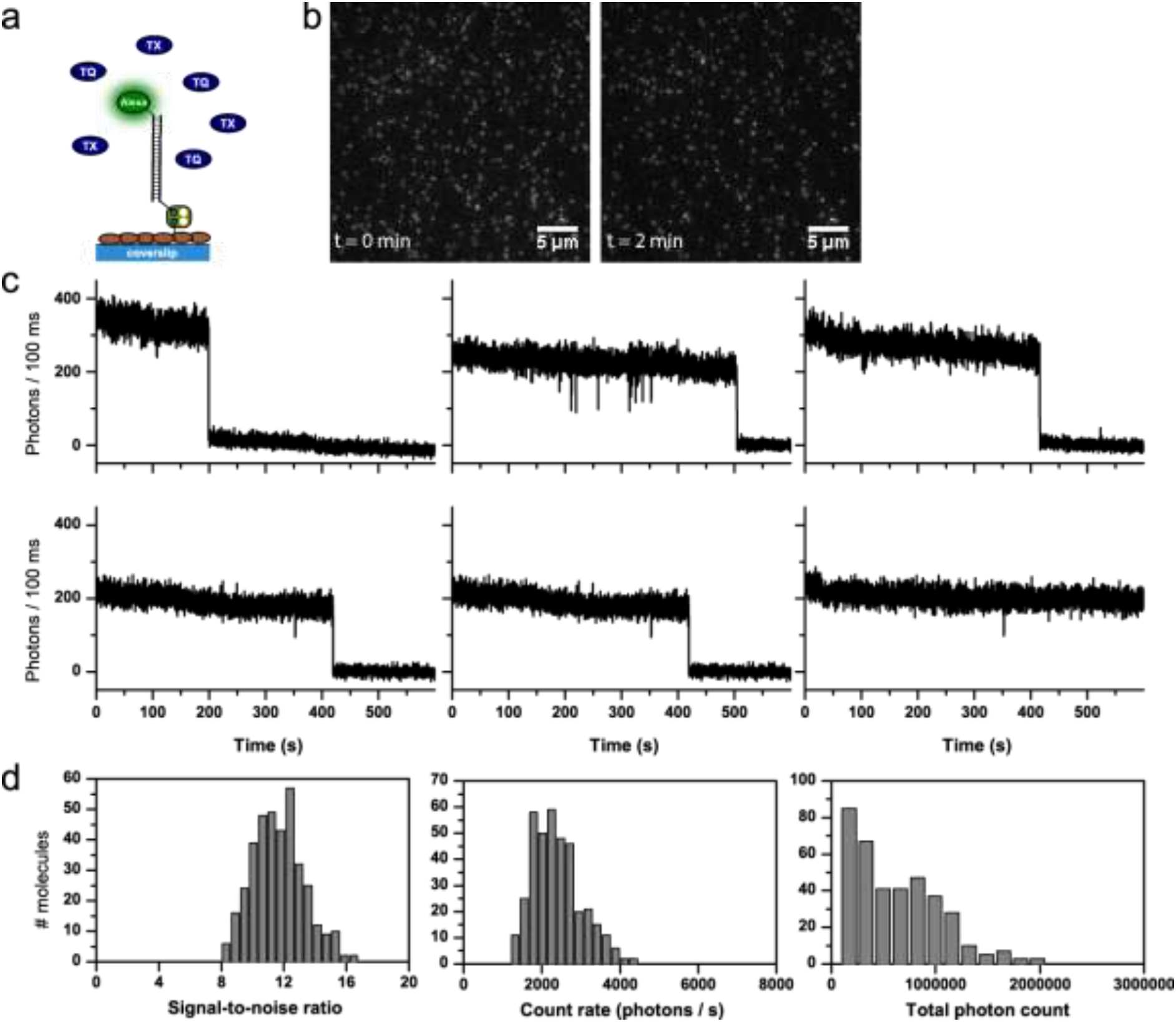
Detailed photophysical characterization of Alexa555 (buffer with 2 mM TX). a) Schematic representation of dye conjugation on DNA immobilized on a BSA/BSA-Biotin surface. b) TIRF images at different points in time, brightness and contrast 2083 to 16300 (AD counts). c) Representative time traces. d) Histograms of signal-to-noise ratio, count-rate and total photon count from individual time traces.

**Figure S9:**
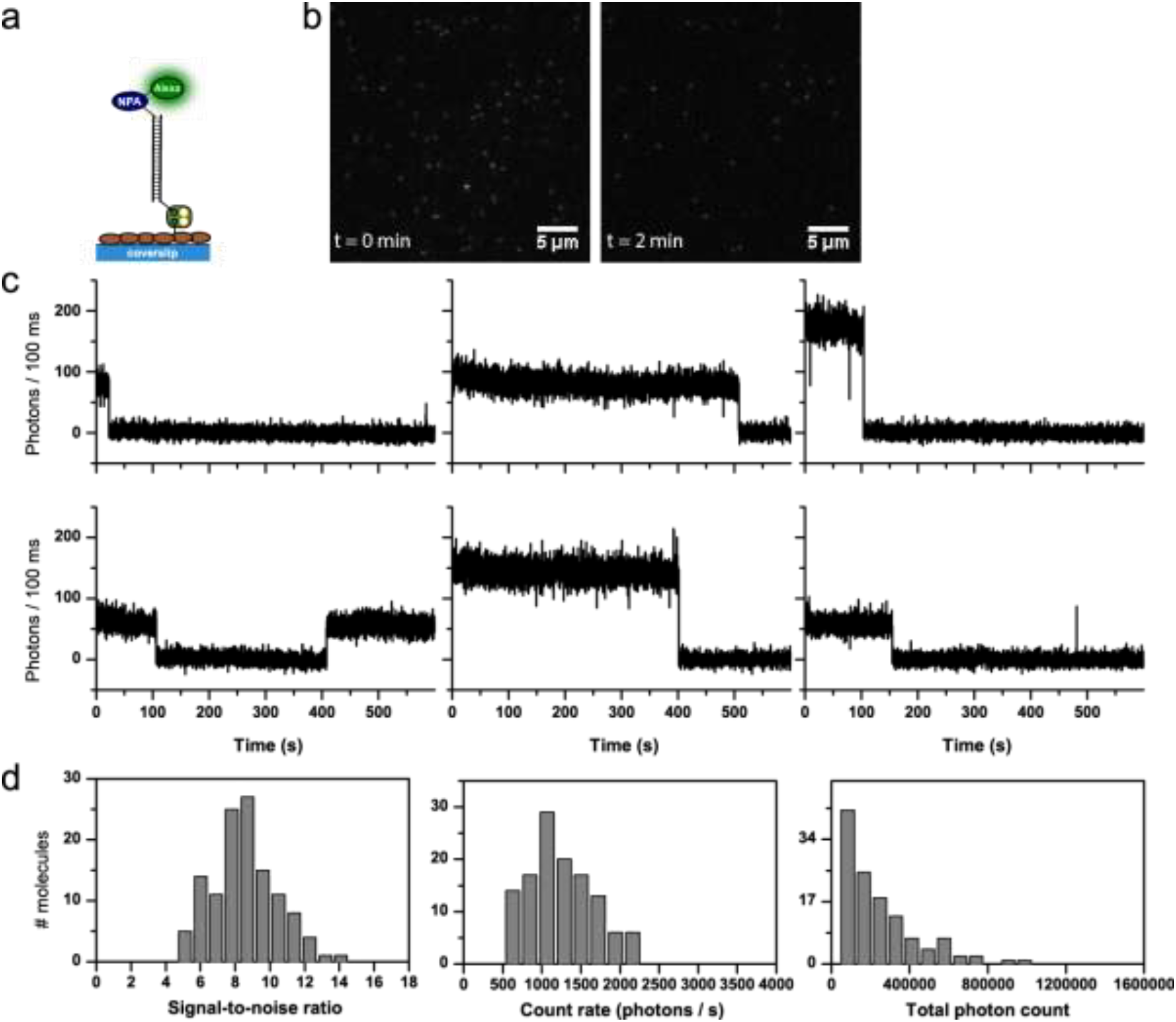
Detailed photophysical characterization of NPA-Alexa555 (GOX buffer, no photostabilizer in solution). a) Schematic representation of photostabilizer-dye conjugates on DNA immobilized on a BSA/BSA-Biotin surface. b) TIRF images at different points in time, brightness and contrast 1904 to 19752 (AD counts). c) Representative time traces. d) Histograms of signal-to-noise ratio, count-rate and total photon count from individual time traces.

**Figure S10:**
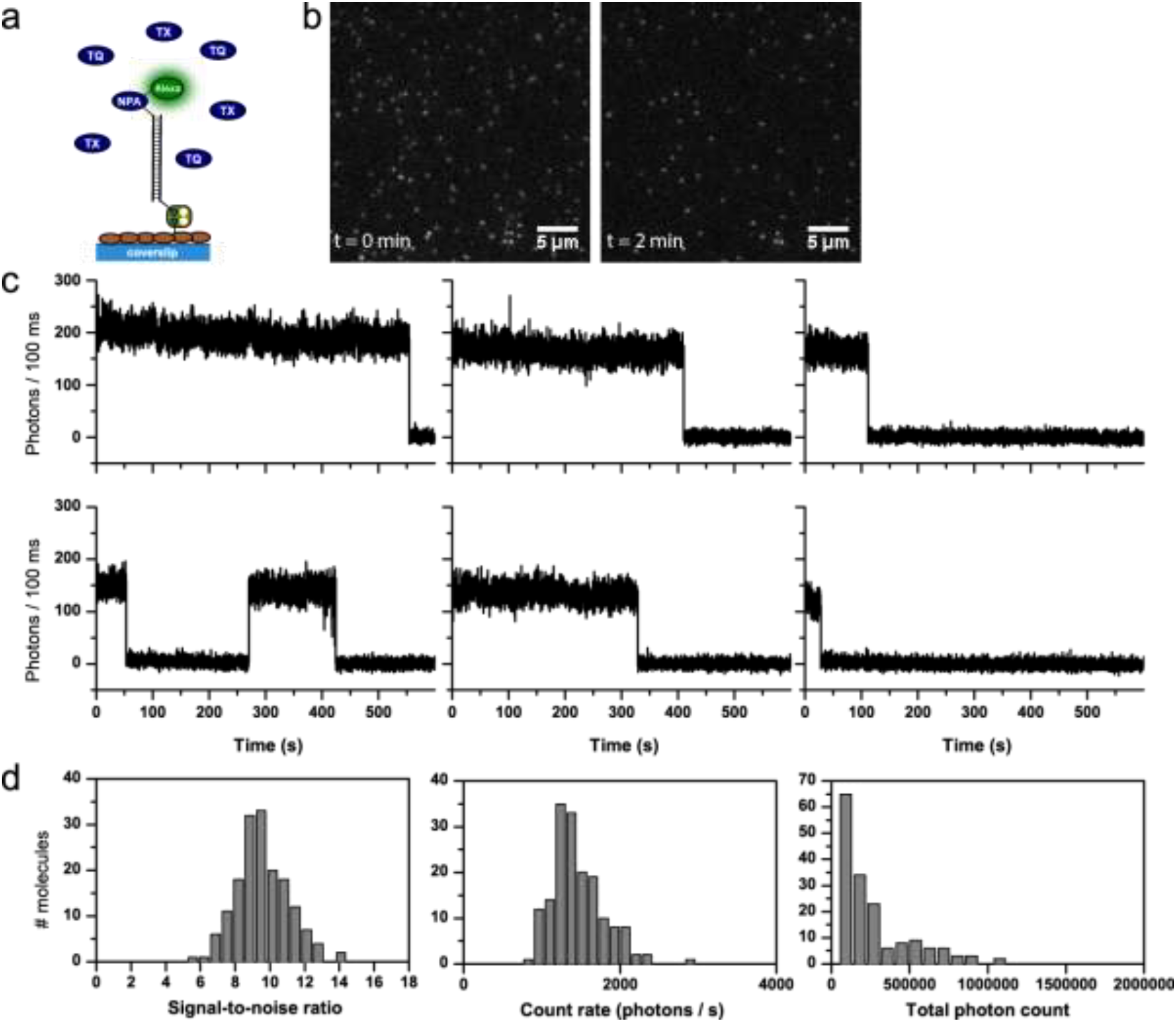
Detailed photophysical characterization of NPA-Alexa555 (buffer with 2 mM TX). a) Schematic representation of photostabilizer-dye conjugates on DNA immobilized on a BSA/BSA-Biotin surface. b) TIRF images at different points in time, brightness and contrast 1888 to 10744 (AD counts). c) Representative time traces. d) Histograms of signal-to-noise ratio, count-rate and total photon count from individual time traces.

**Figure S11:**
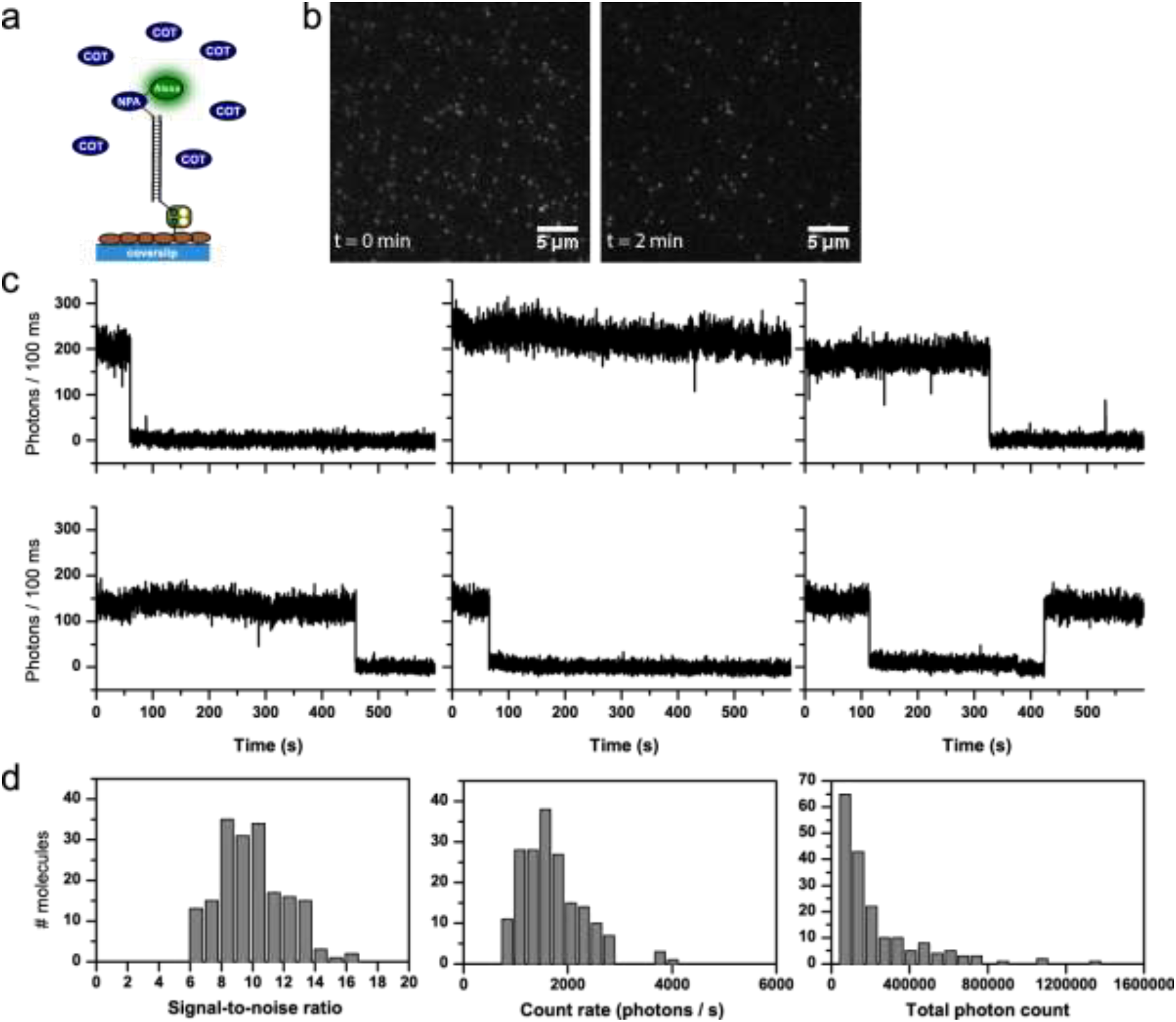
Detailed photophysical characterization of NPA-Alexa555 (buffer with 2 mM COT). a) Schematic representation of photostabilizer-dye conjugates on DNA immobilized on a BSA/BSA-Biotin surface. b) TIRF images at different points in time, brightness and contrast 1939 to 13272 (AD counts). c) Representative time traces. d) Histograms of signal-to-noise ratio, count-rate and total photon count from individual time traces.

**Figure S12:**
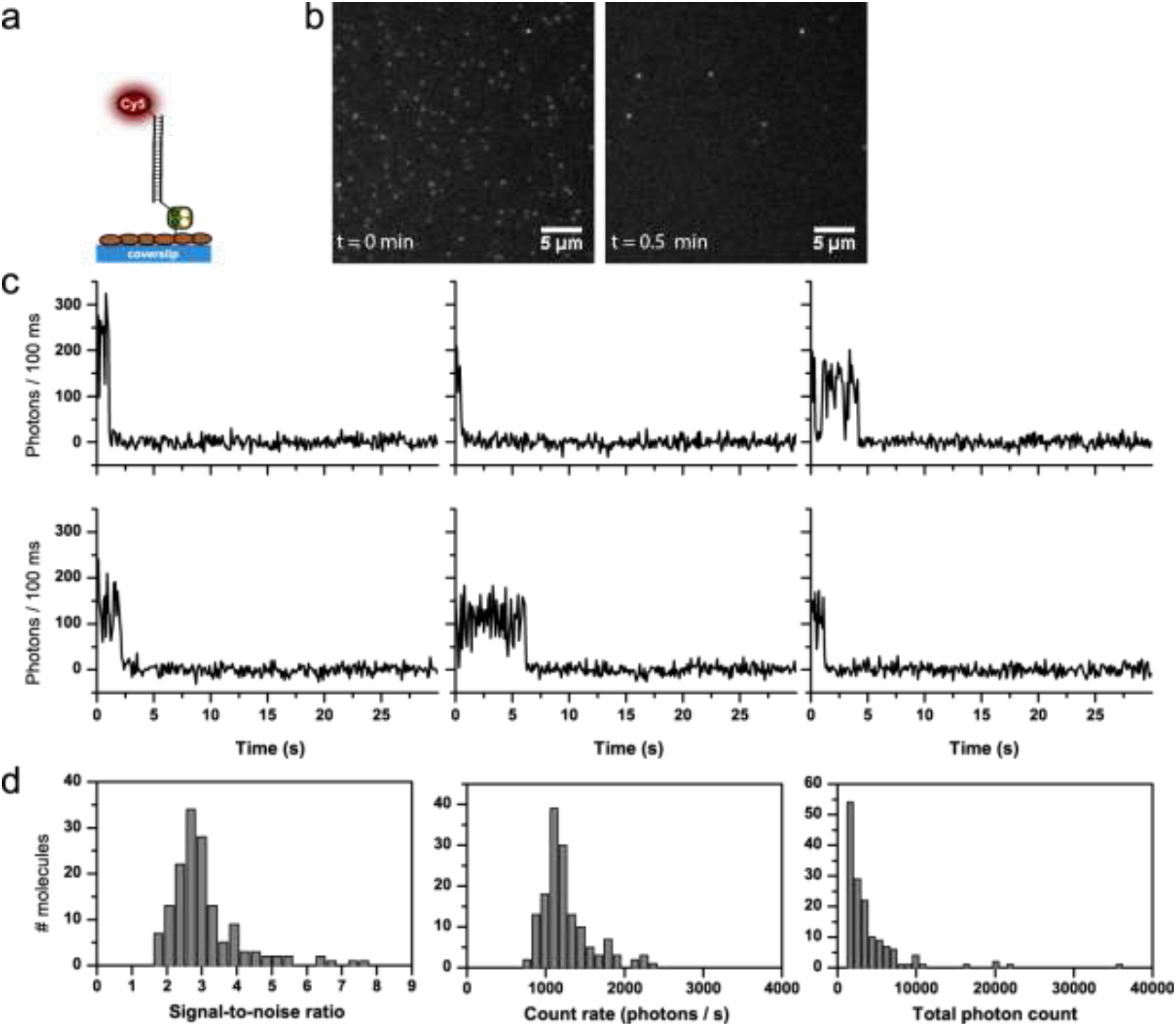
Detailed photophysical characterization of Cy5 (GOX buffer, no photostabilizer in solution). a) Schematic representation of dye conjugation on DNA immobilized on a BSA/BSA-Biotin surface. b) TIRF images at different points in time, brightness and contrast 6766 to 49567 (AD counts). c) Representative time traces. d) Histograms of signal-to-noise ratio, count-rate and total photon count from individual time traces.

**Figure S13:**
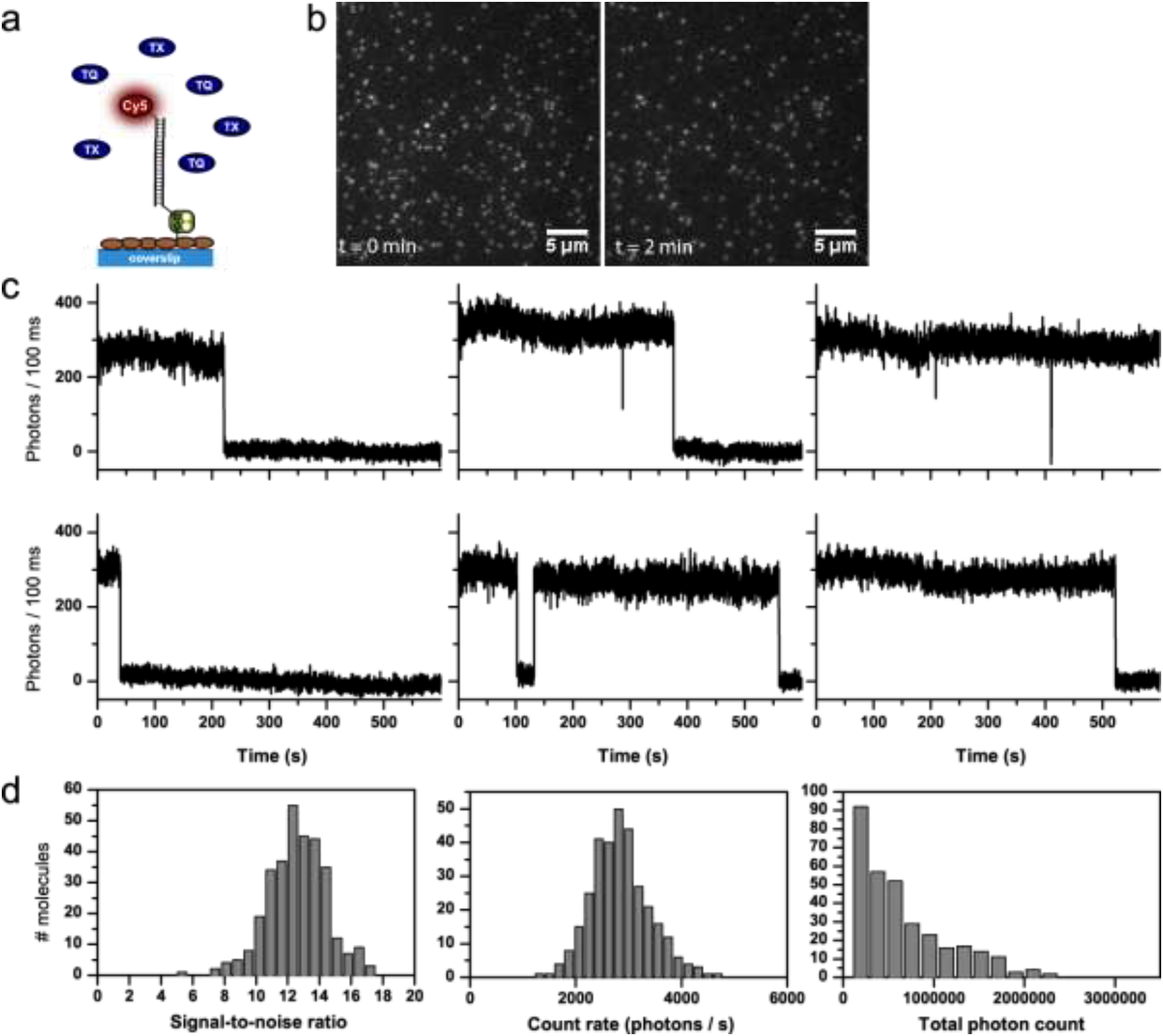
Detailed photophysical characterization of Cy5 (buffer with 2 mM TX). a) Schematic representation of dye conjugation on DNA immobilized on a BSA/BSA-Biotin surface. b) TIRF images at different points in time, brightness and contrast 6705 to 65535 (AD counts). c) Representative time traces. d) Histograms of signal-to-noise ratio, count-rate and total photon count from individual time traces.

**Figure S14:**
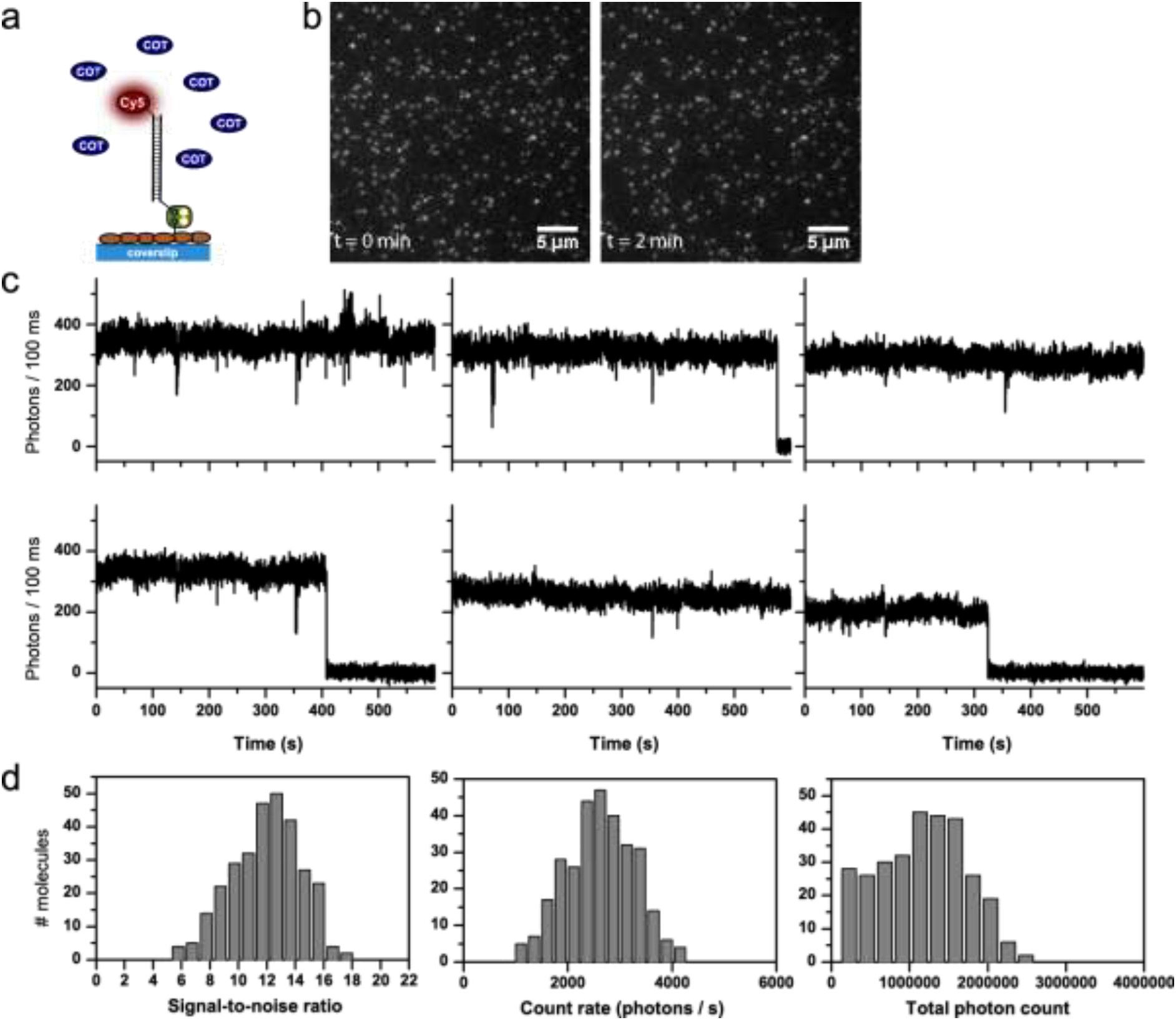
Detailed photophysical characterization of Cy5 (buffer with 2 mM COT). a) Schematic representation of dye conjugation on DNA immobilized on a BSA/BSA-Biotin surface. b) TIRF images at different points in time, brightness and contrast 8355 to 64081 (AD counts). c) Representative time traces. d) Histograms of signal-to-noise ratio, count-rate and total photon count from individual time traces.

**Figure S15:**
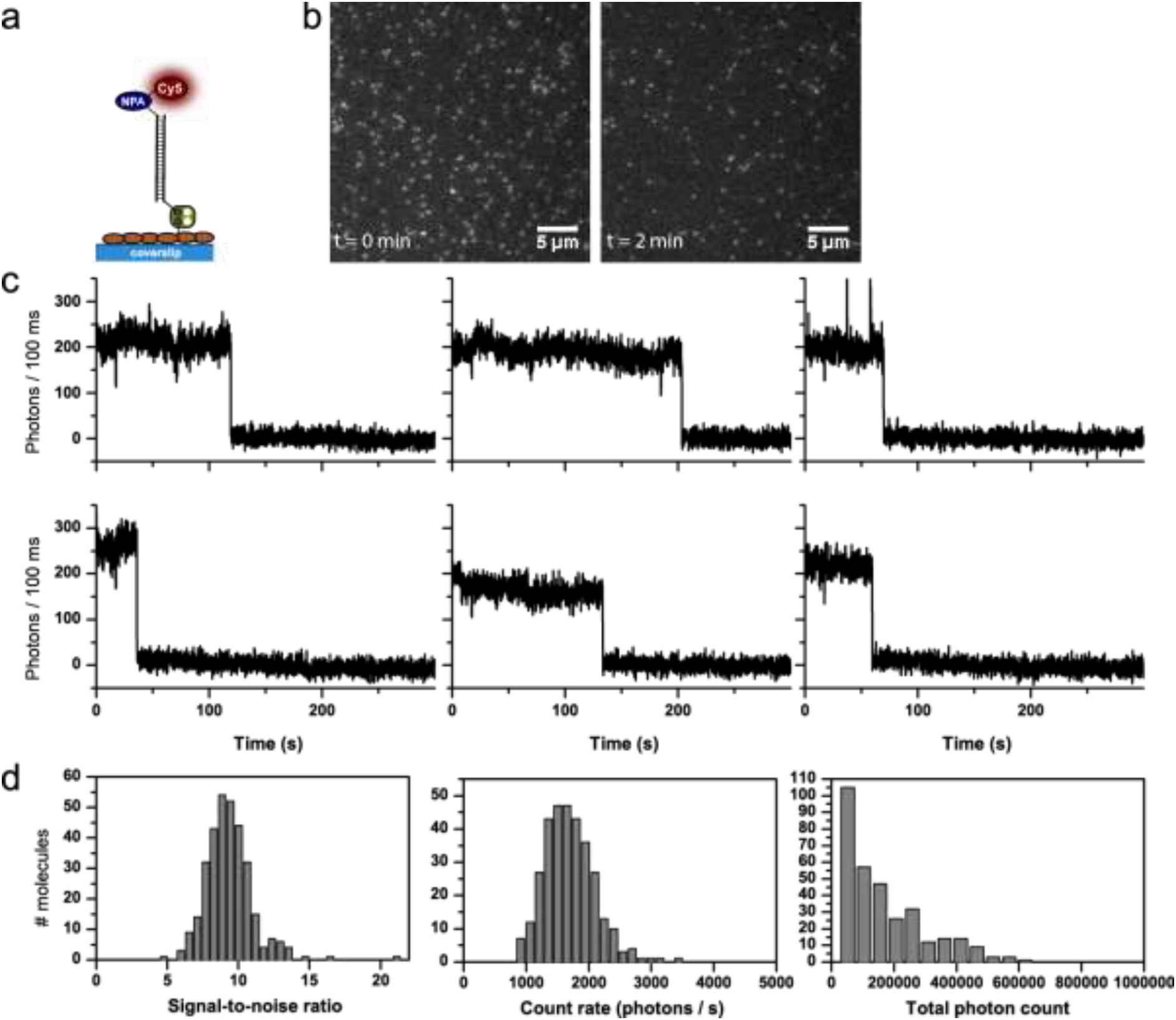
Detailed photophysical characterization of NPA-Cy5 (GOX buffer, no photostabilizer in solution). a) Schematic representation of photostabilizer-dye conjugates on DNA immobilized on a BSA/BSA-Biotin surface. b) TIRF images at different points in time, brightness and contrast 5820 to 43075 (AD counts). c) Representative time traces. d) Histograms of signal-to-noise ratio, count-rate and total photon count from individual time traces.

**Figure S16:**
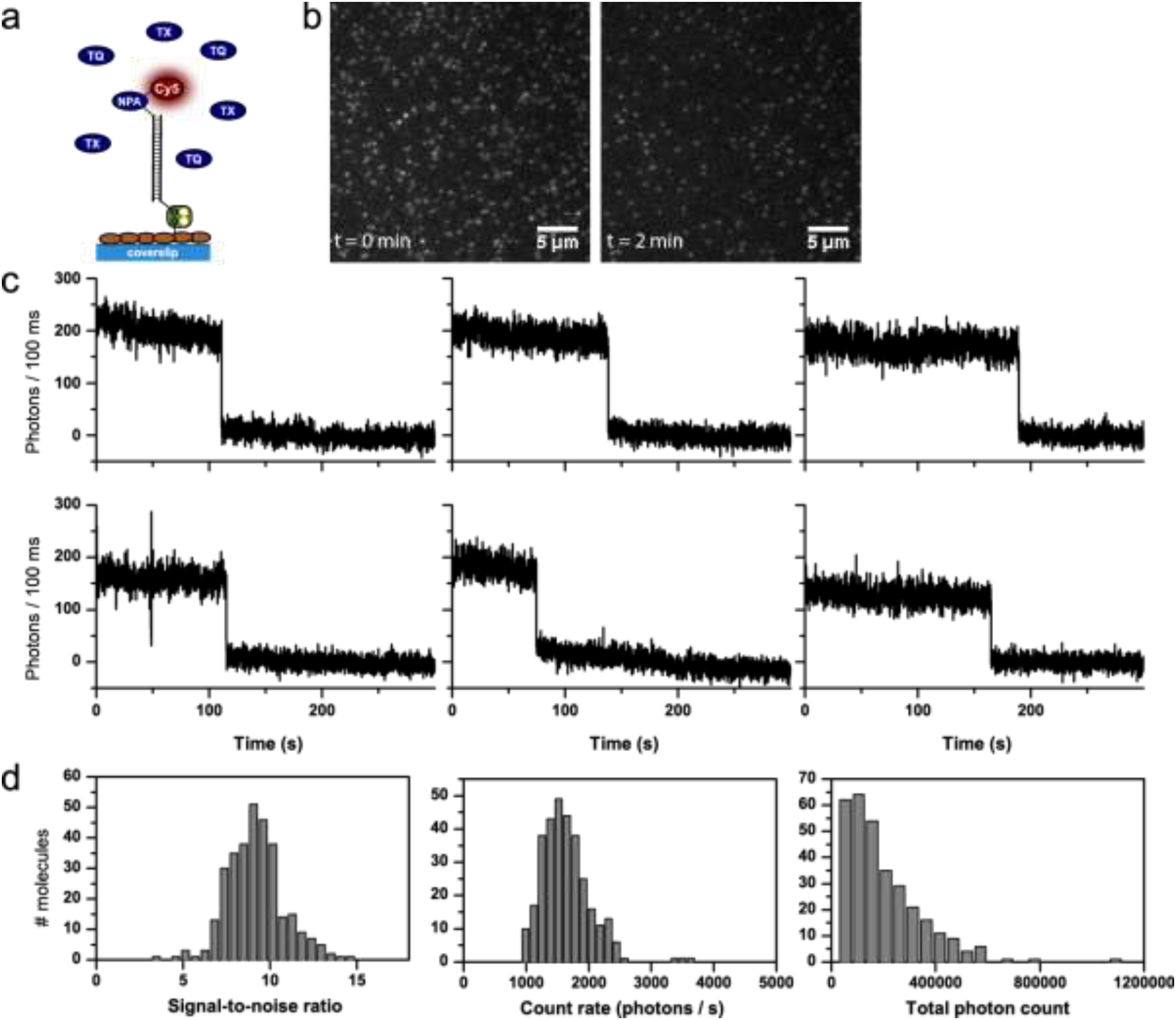
Detailed photophysical characterization of NPA-Cy5 (buffer with 2 mM TX). a) Schematic representation of photostabilizer-dye conjugates on DNA immobilized on a BSA/BSA-Biotin surface. b) TIRF images at different points in time, brightness and contrast 7408 to 52325 (AD counts). c) Representative time traces. d) Histograms of signal-to-noise ratio, count-rate and total photon count from individual time traces.

**Figure S17:**
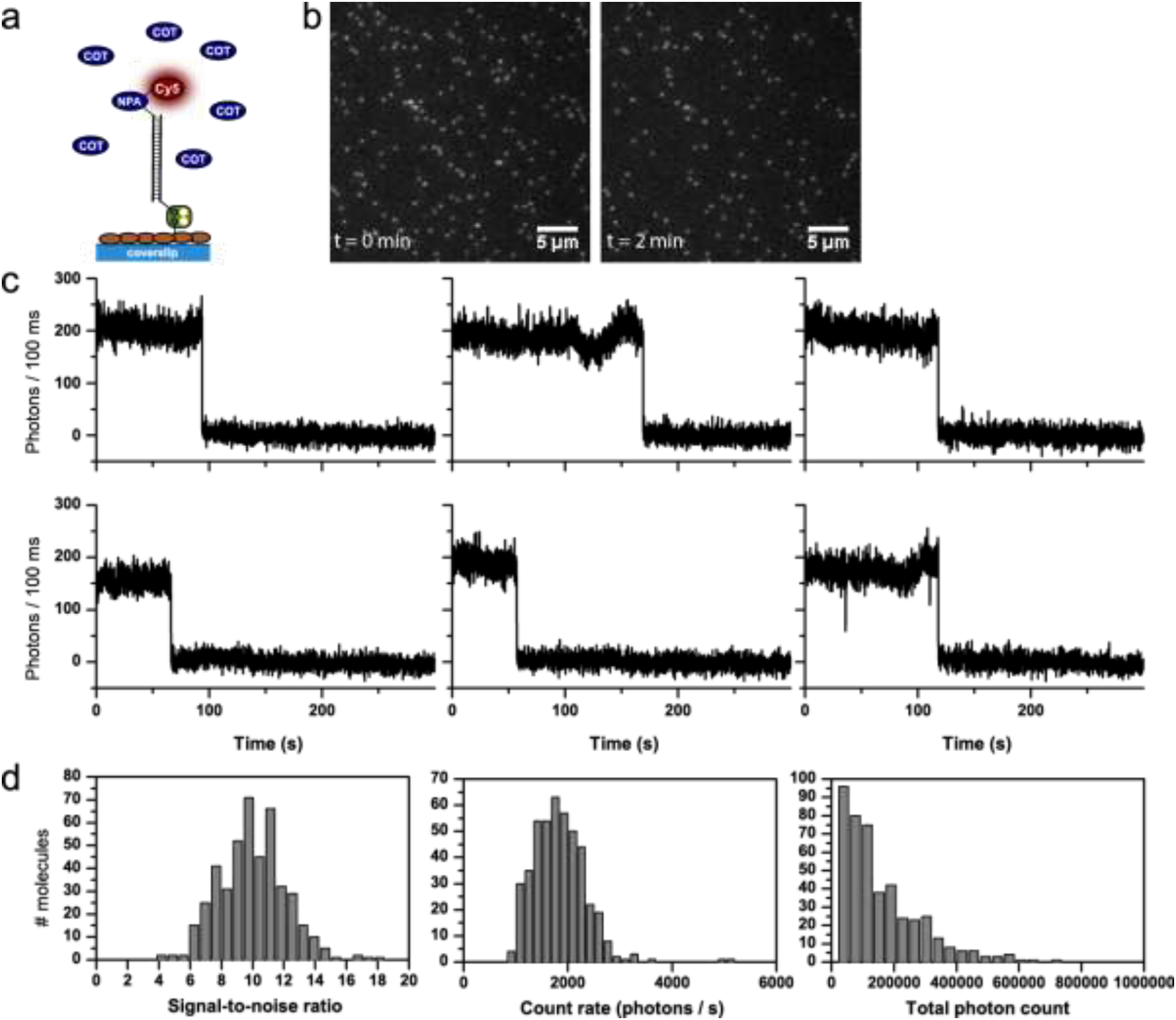
Detailed photophysical characterization of NPA-Cy5 (buffer with 2 mM COT). a) Schematic representation of photostabilizer-dye conjugates on DNA immobilized on a BSA/BSA-Biotin surface. b) TIRF images at different points in time, brightness and contrast 8342 to 54037 (AD counts). c) Representative time traces. d) Histograms of signal-to-noise ratio, count-rate and total photon count from individual time traces.

**Figure S18:**
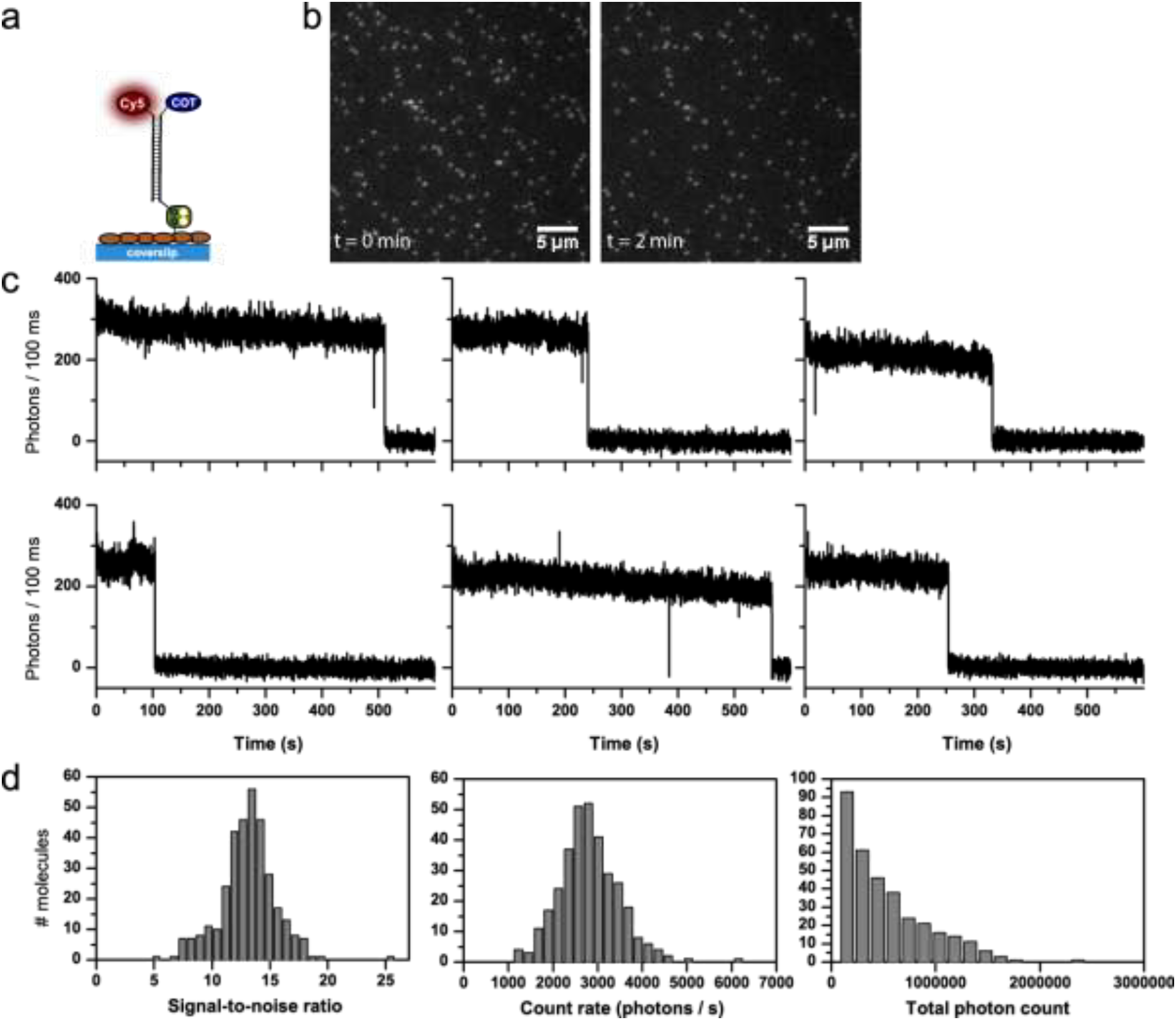
Detailed photophysical characterization of Cy5-COT (GOX buffer, no photostabilizer in solution). a) Schematic representation of photostabilizer-dye conjugates on DNA immobilized on a BSA/BSA-Biotin surface. b) TIRF images at different points in time, brightness and contrast 6413 to 60344 (AD counts). c) Representative time traces. d) Histograms of signal-to-noise ratio, count-rate and total photon count from individual time traces.

**Figure S19:**
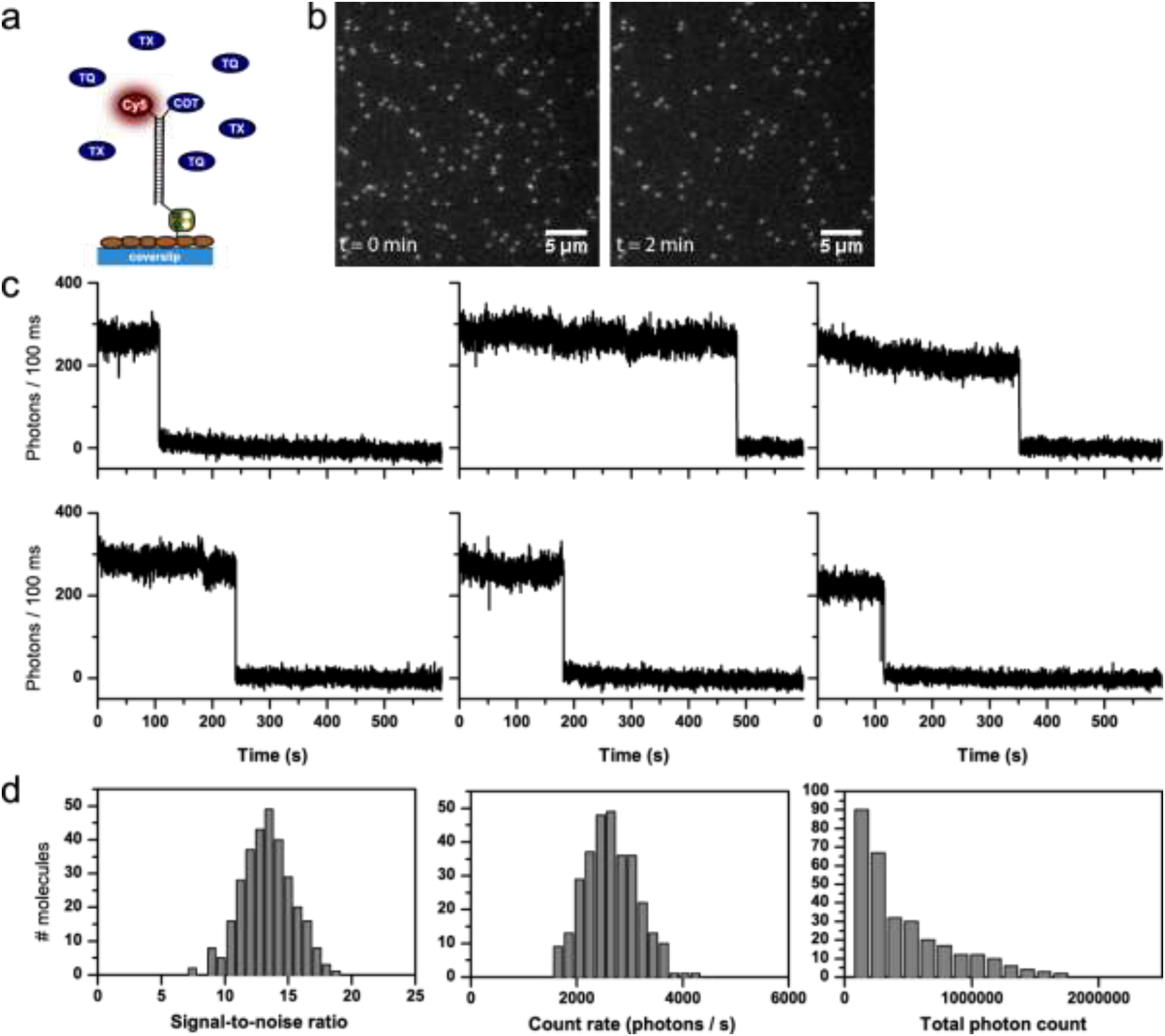
Detailed photophysical characterization of Cy5-COT (buffer with 2 mM TX). a) Schematic representation of photostabilizer-dye conjugates on DNA immobilized on a BSA/BSA-Biotin surface. b) TIRF images at different points in time, brightness and contrast 8143 to 62500 (AD counts). c) Representative time traces. d) Histograms of signal-to-noise ratio, count-rate and total photon count from individual time traces.

**Figure S20:**
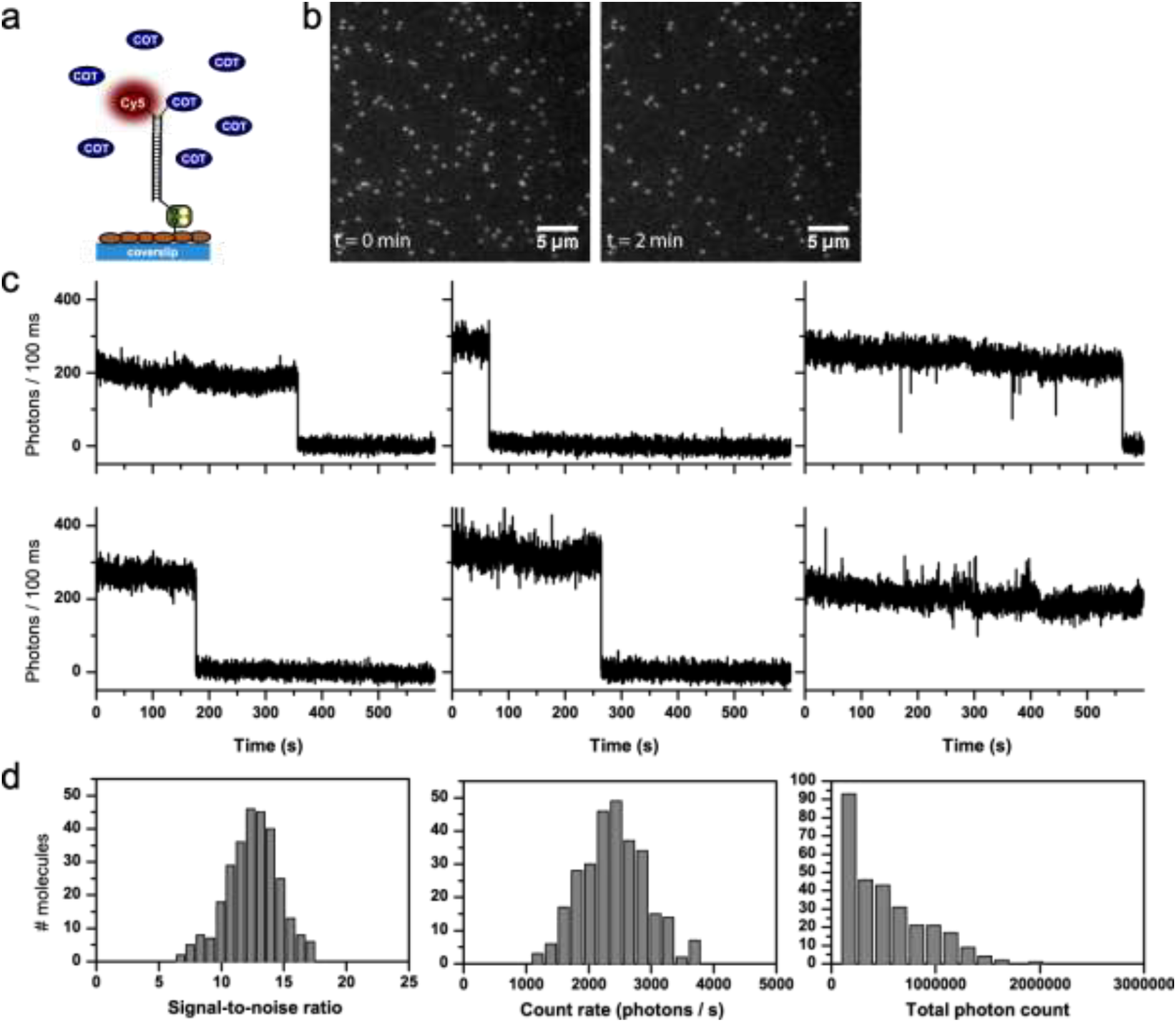
Detailed photophysical characterization of Cy5-COT (buffer with 2 mM TX). a) Schematic representation of photostabilizer-dye conjugates on DNA immobilized on a BSA/BSA-Biotin surface. b) TIRF images at different points in time, brightness and contrast 7804 to 44206 (AD counts). c) Representative time traces. d) Histograms of signal-to-noise ratio, count-rate and total photon count from individual time traces.

**Figure S21:**
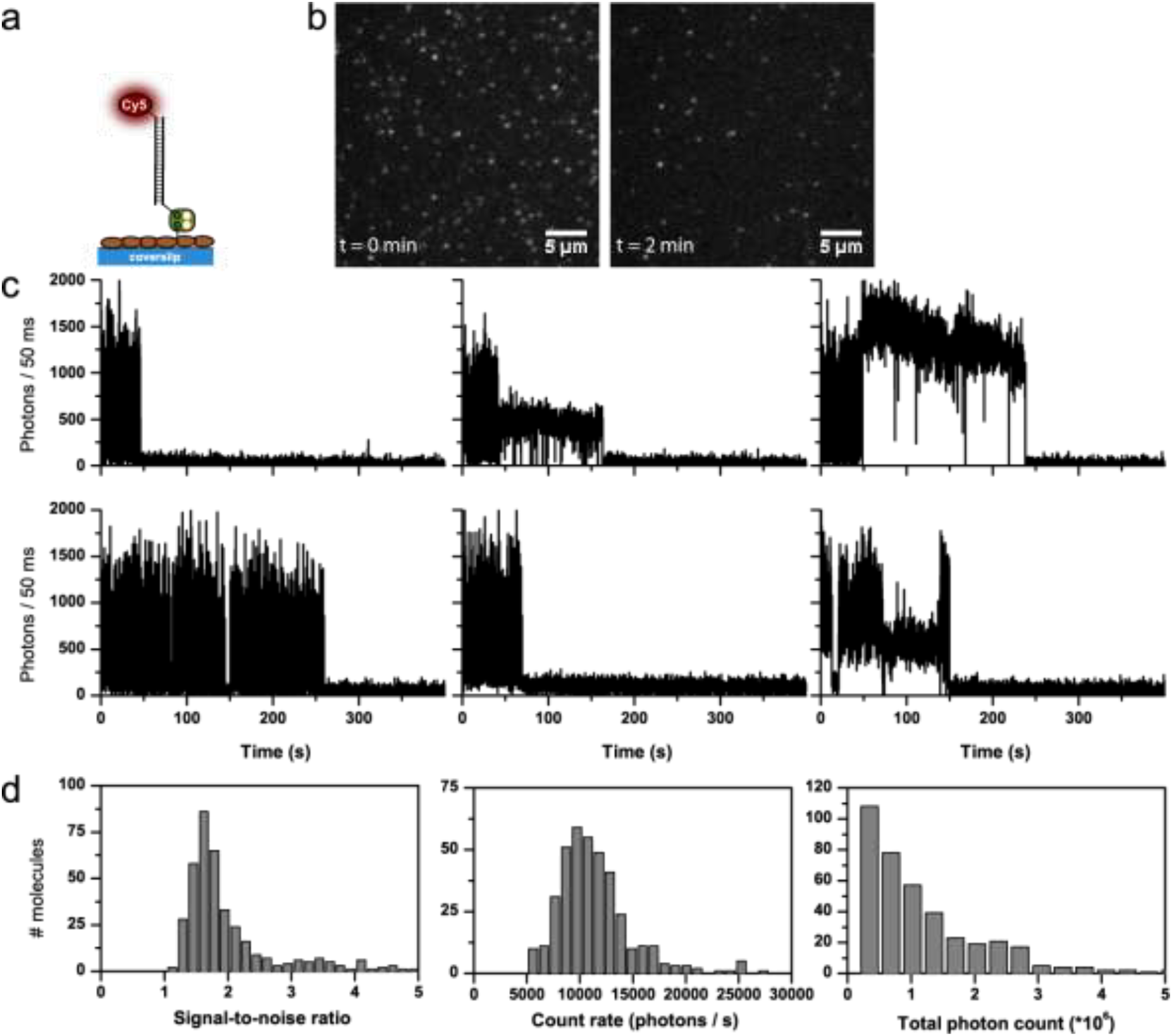
Detailed photophysical characterization of Cy5 (GOX buffer, no photostabilizer in solution) as a control experiment for Figures S23-S24. a) Schematic representation of dye conjugation on DNA immobilized on a BSA/BSA-Biotin surface. b) TIRF images at different points in time, brightness and contrast 3396 to 50600 (AD counts). c) Representative time traces. d) Histograms of signal-to-noise ratio, count-rate and total photon count from individual time traces.

**Figure S22:**
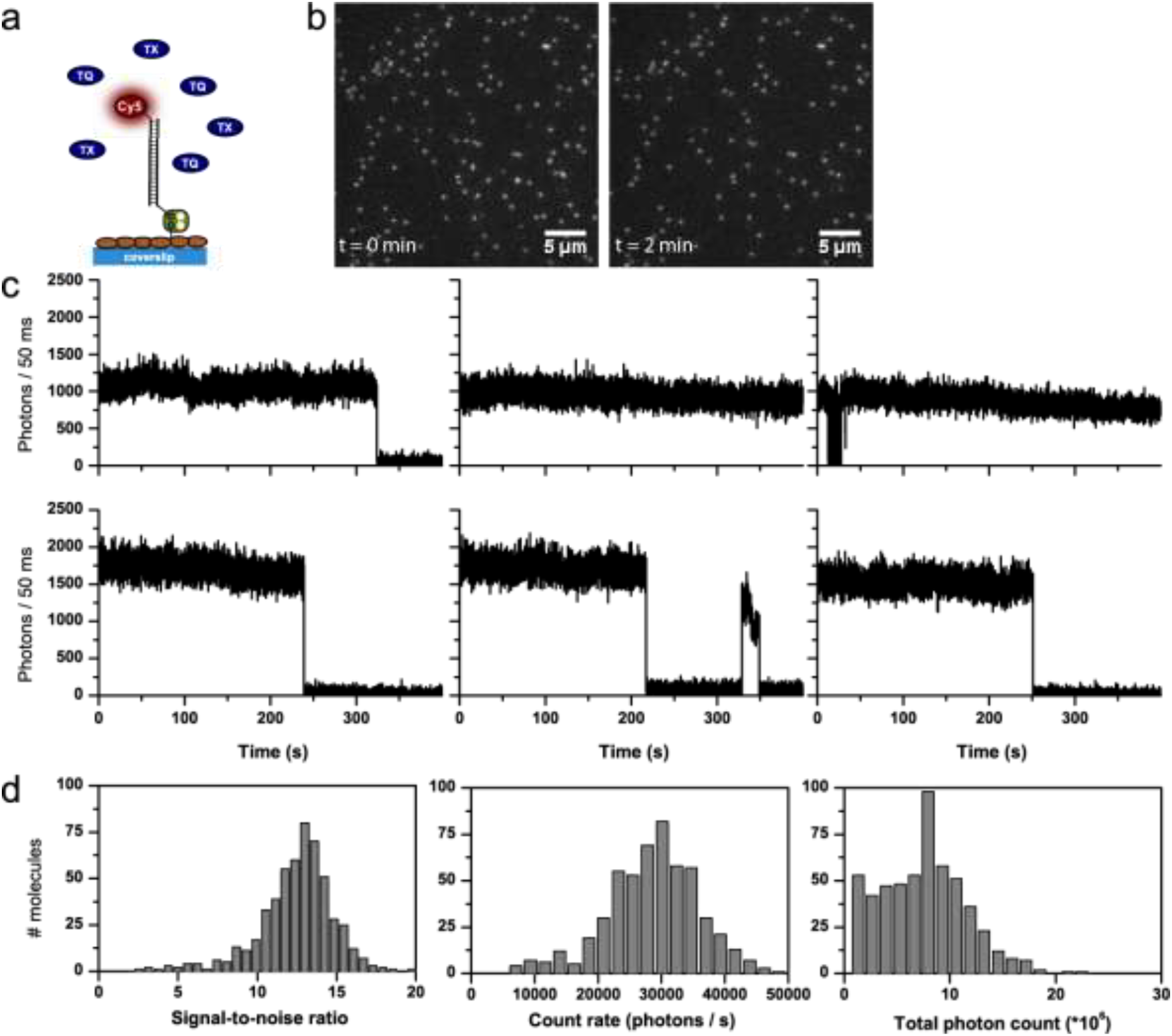
Detailed photophysical characterization of Cy5 (buffer with 2 mM TX) as a control experiment for Figures S23-S24. a) Schematic representation of dye conjugation on DNA immobilized on a BSA/BSA-Biotin surface. b) TIRF images at different points in time, brightness and contrast 3147 to 65535 (AD counts). c) Representative time traces. d) Histograms of signal-to-noise ratio, count-rate and total photon count from individual time traces.

**Figure S23:**
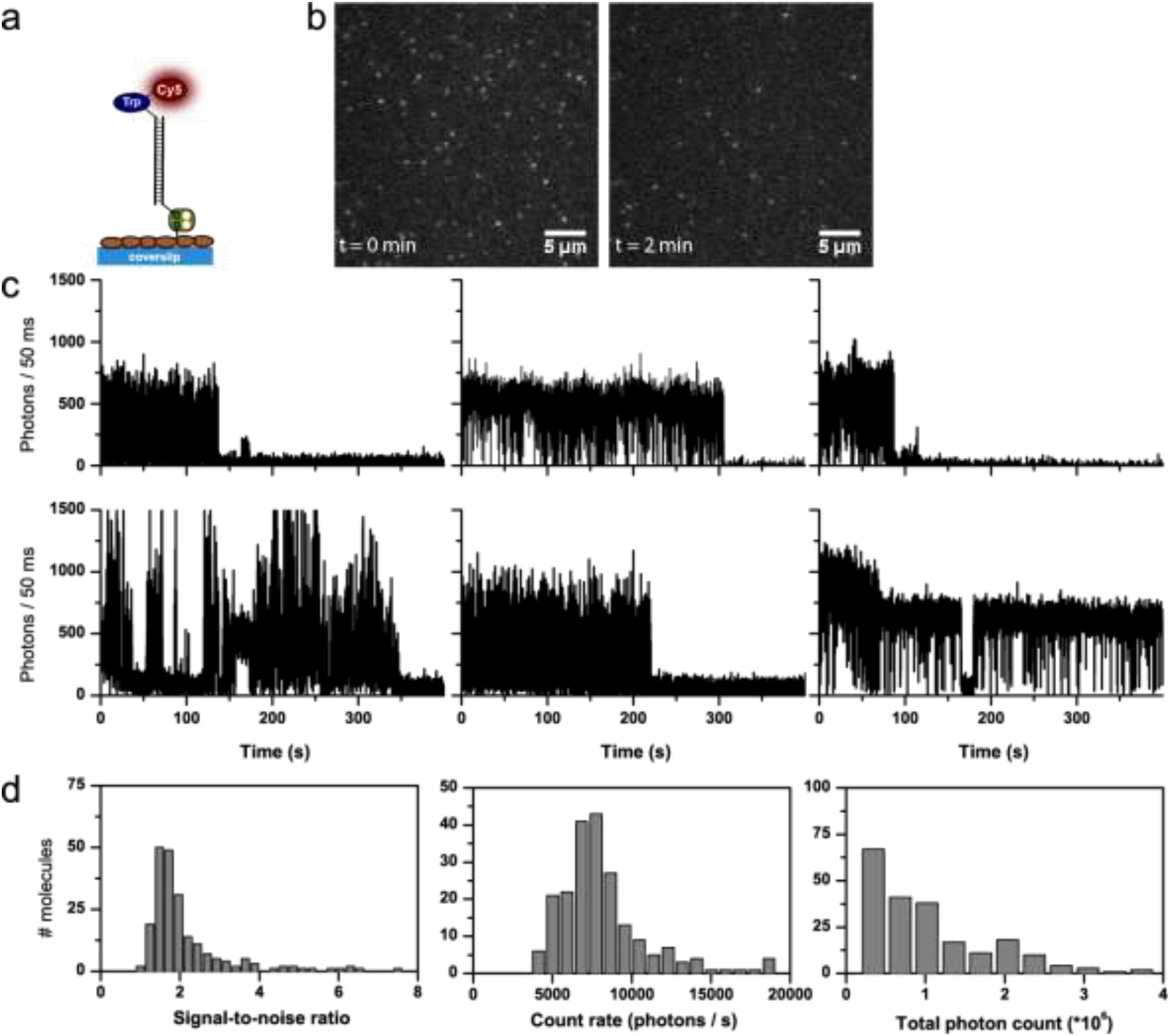
Detailed photophysical characterization of Trp-Cy5 (GOX buffer, no photostabilizer in solution). a) Schematic representation of Trp-dye conjugate on DNA immobilized on a BSA/BSA-Biotin surface. b) TIRF images at different points in time, brightness and contrast 2990 to 29808 (AD counts). c) Representative time traces. d) Histograms of signal-to-noise ratio, count-rate and total photon count from individual time traces.

**Figure S24:**
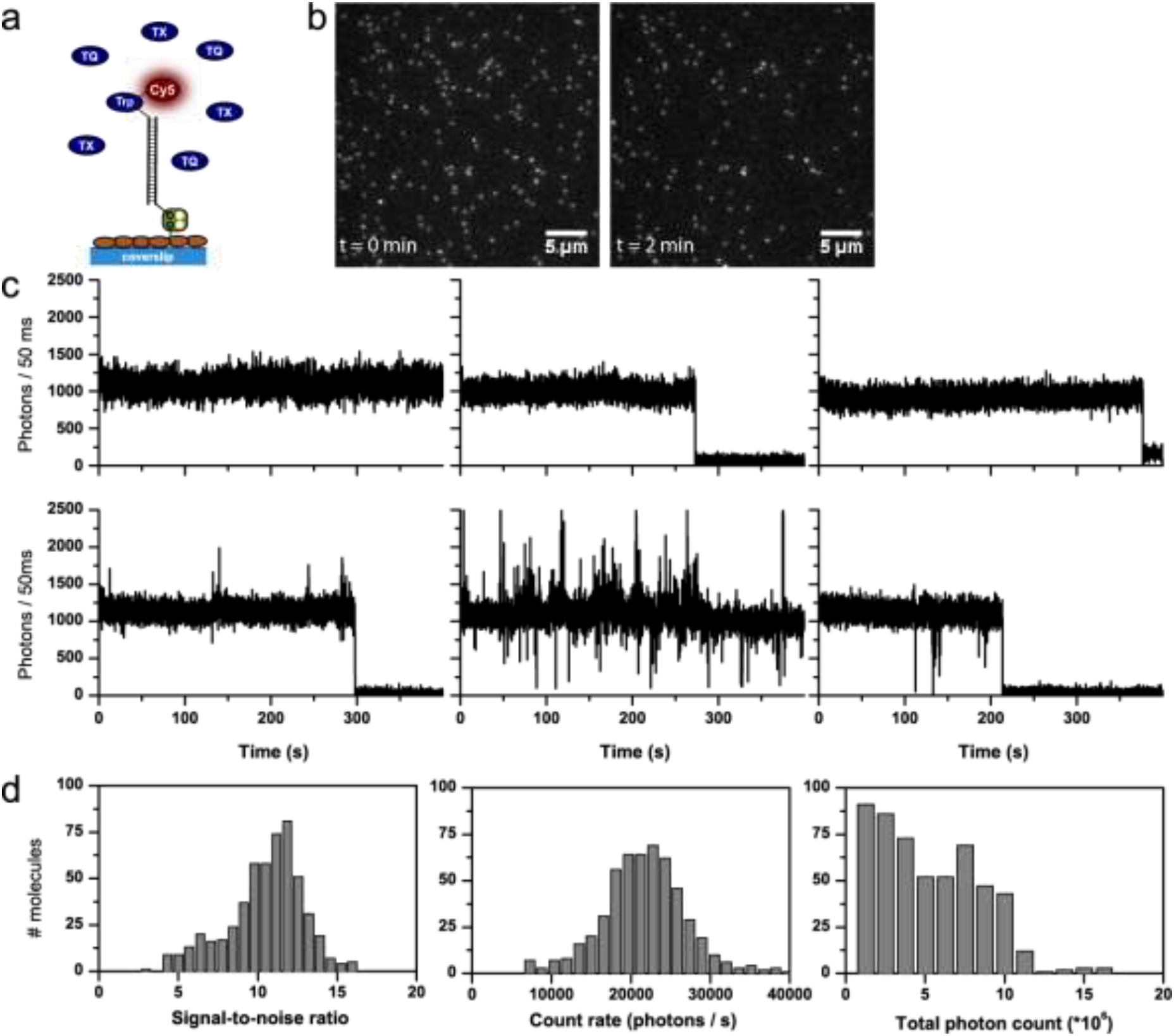
Detailed photophysical characterization of Trp-Cy5 (buffer with 2 mM TX). a) Schematic representation of Trp-dye conjugate on DNA immobilized on a BSA/BSA-Biotin surface. b) TIRF images at different points in time, brightness and contrast 3627 to 50895 (AD counts). c) Representative time traces. d) Histograms of signal-to-noise ratio, count-rate and total photon count from individual time traces.

**Figure S25:**
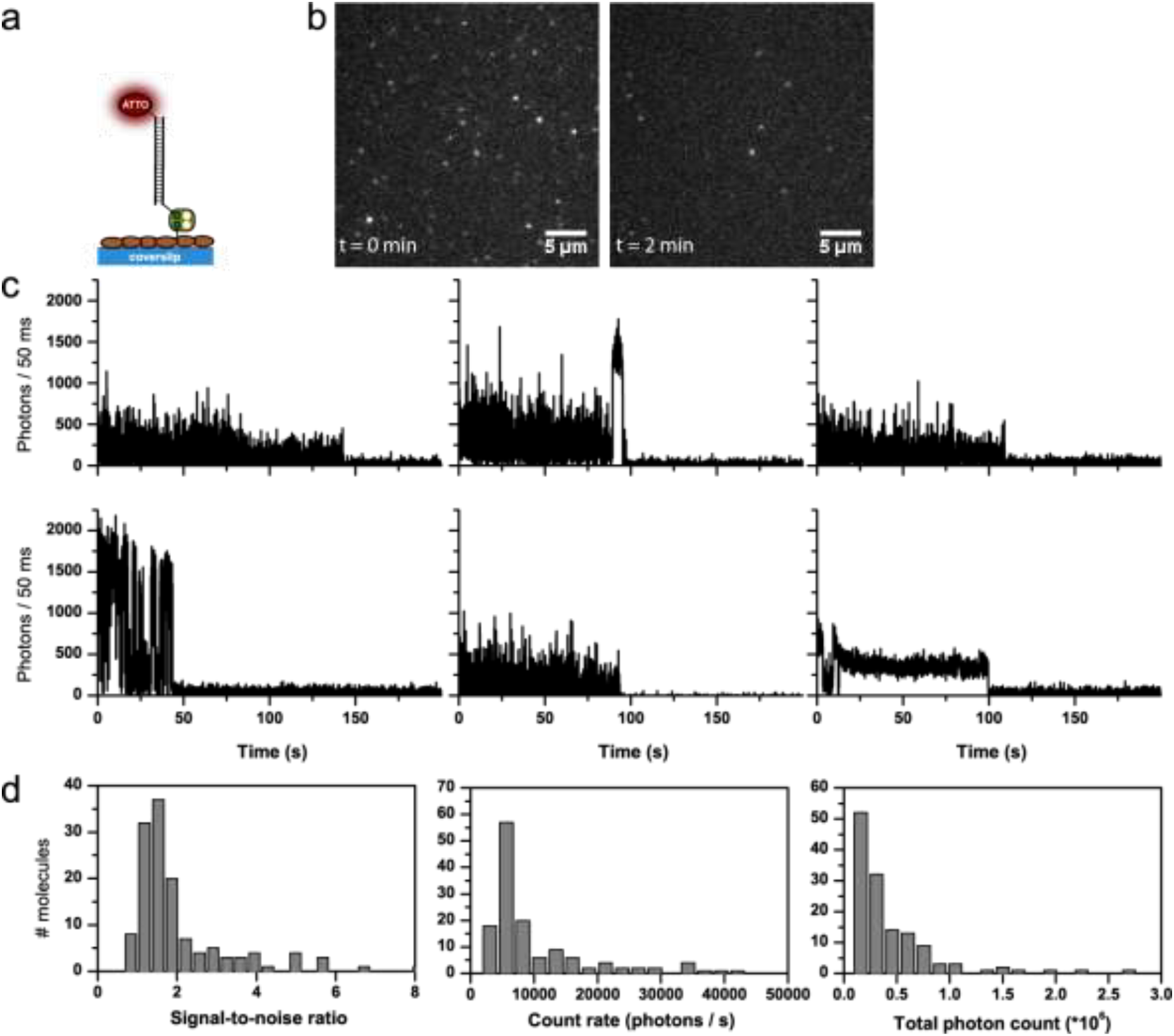
Detailed photophysical characterization of ATTO647N (GOX buffer, no photostabilizer in solution) as a control experiment for Figures S27-S28. a) Schematic representation of dye conjugation on DNA immobilized on a BSA/BSA-Biotin surface. b) TIRF images at different points in time, brightness and contrast 3031 to 26808 (AD counts). c) Representative time traces. d) Histograms of signal-to-noise ratio, count-rate and total photon count from individual time traces.

**Figure S26:**
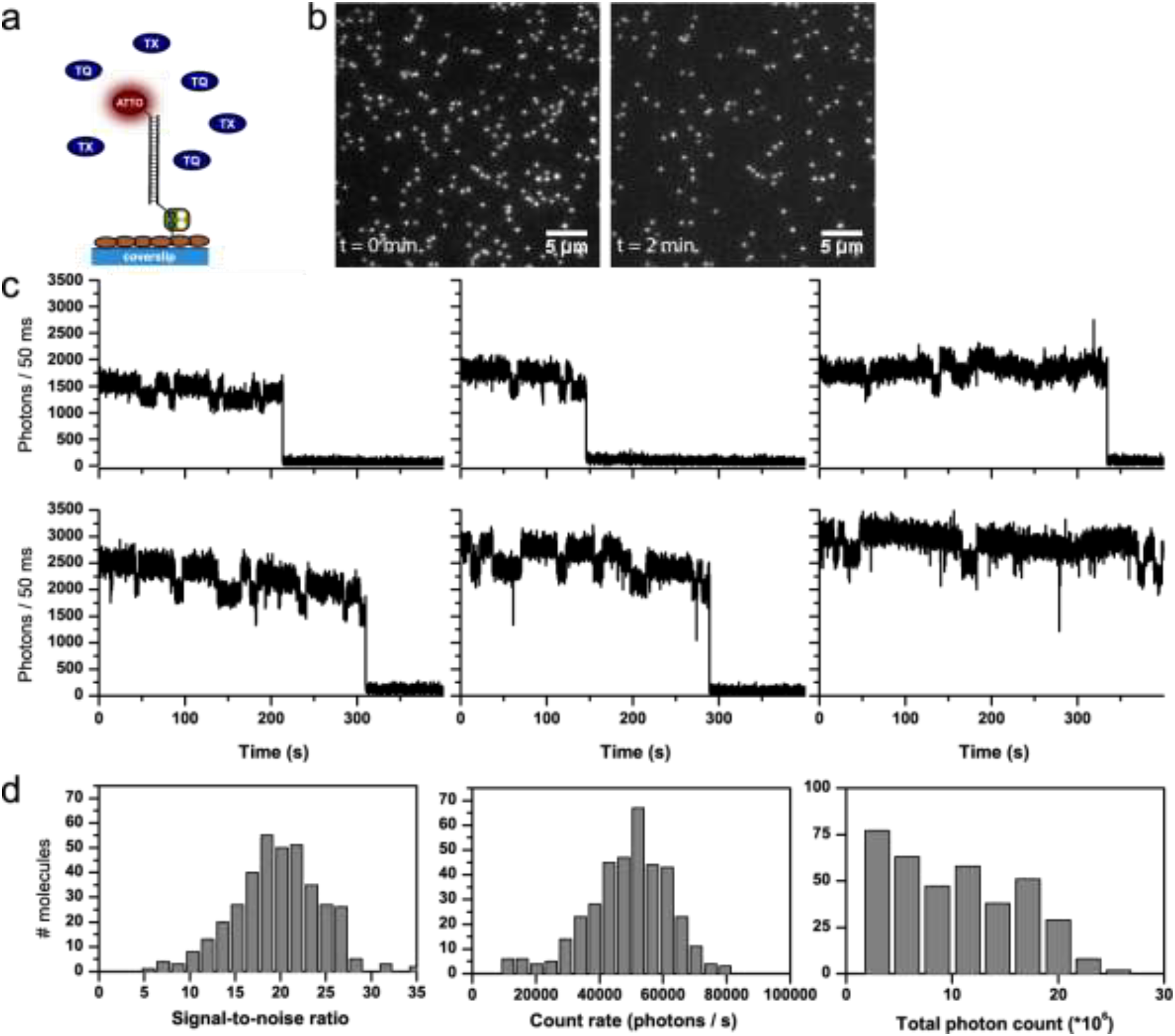
Detailed photophysical characterization of ATTO647N (buffer with 2 mM TX) as a control experiment for Figures S27-S28. a) Schematic representation of dye conjugation on DNA immobilized on a BSA/BSA-Biotin surface. b) TIRF images at different points in time, brightness and contrast 4400 to 65535 (AD counts). c) Representative time traces. d) Histograms of signal-to-noise ratio, count-rate and total photon count from individual time traces.

**Figure S27:**
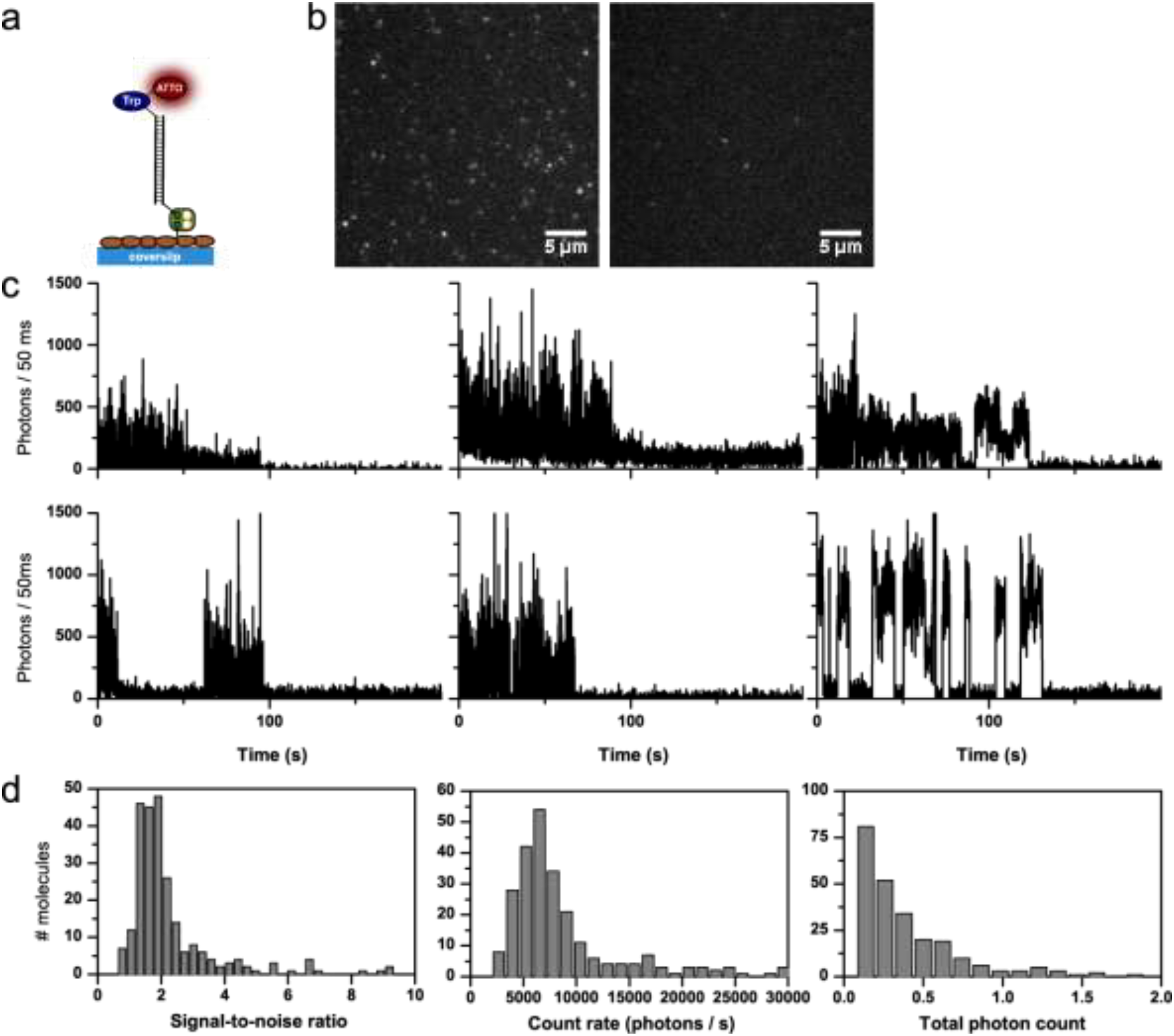
Detailed photophysical characterization of Trp-ATTO647N (GOX buffer, no photostabilizer in solution). a) Schematic representation of Trp-dye conjugates on DNA immobilized on a BSA/BSA-Biotin surface. b) TIRF images at different points in time, brightness and contrast 3780 to 35441 (AD counts). c) Representative time traces. d) Histograms of signal-to-noise ratio, count-rate and total photon count from individual time traces.

**Figure S28:**
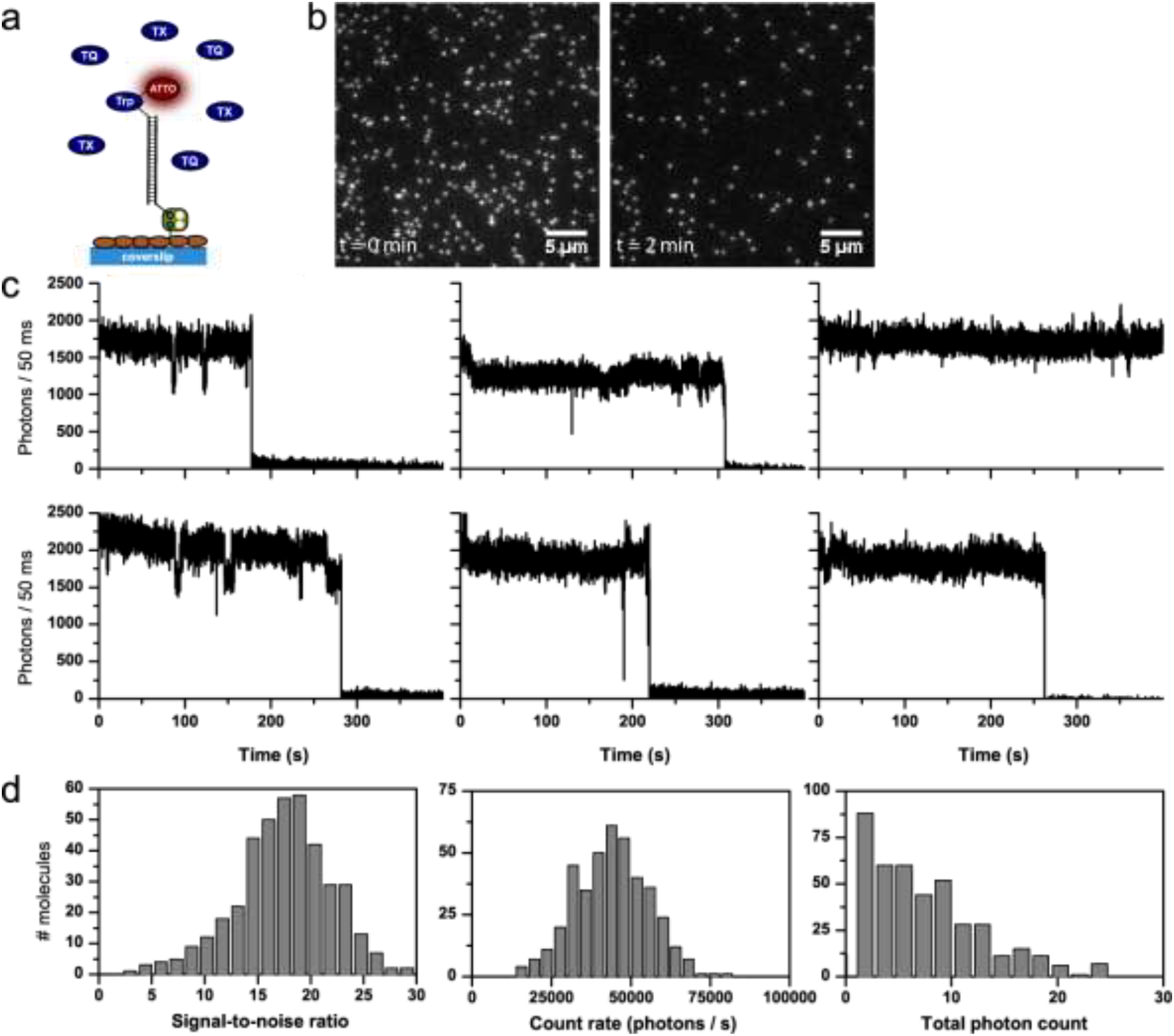
Detailed photophysical characterization of Trp-ATTO647N (buffer with 2 mM TX). a) Schematic representation of Trp-dye conjugate on DNA immobilized on a BSA/BSA-Biotin surface. b) TIRF images at different points in time, brightness and contrast 4686 to 65535 (AD counts). c) Representative time traces. d) Histograms of signal-to-noise ratio, count-rate and total photon count from individual time traces.

**Figure S29:**
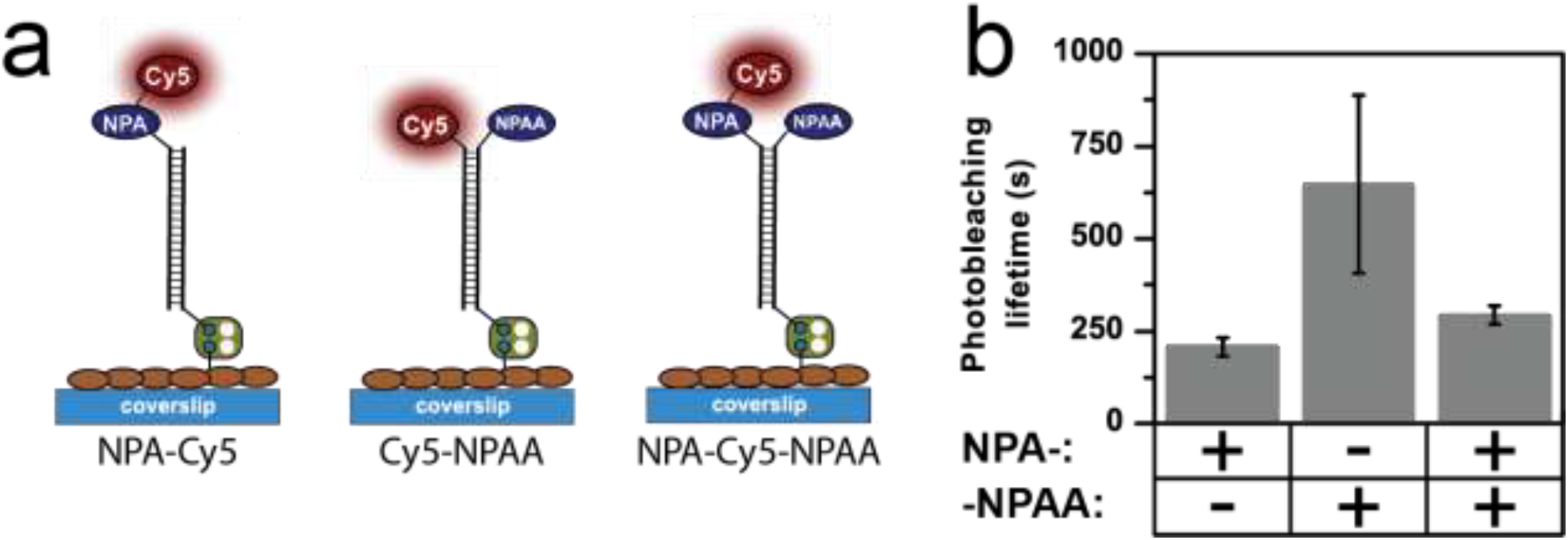
Comparison of Cy5 photostability with different geometries of intramolecular photostabilization by NPA and a combination of both. Error bar show standard deviation of repeats on 3 different days.

**Figure S30:**
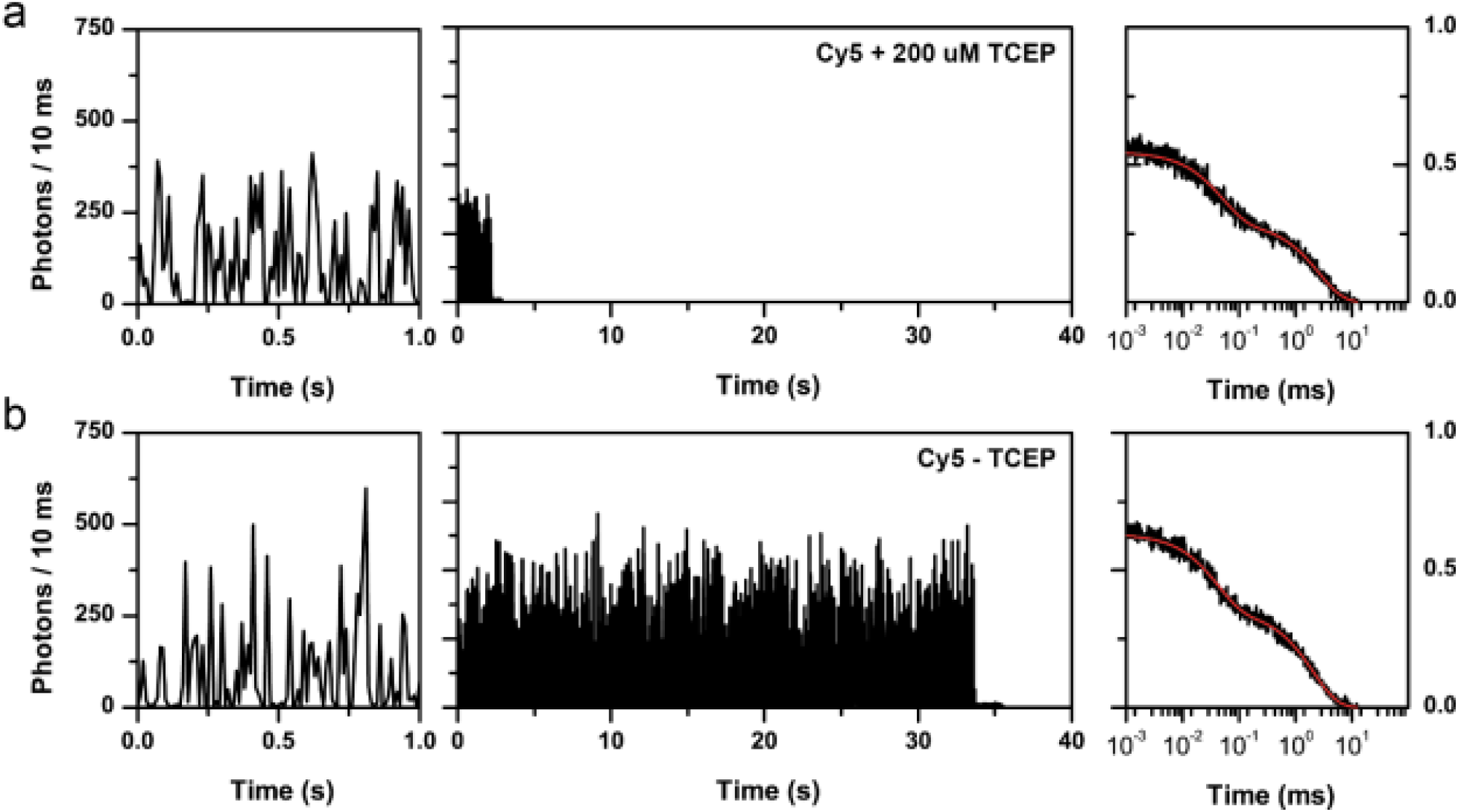
Confocal microscopy transients of Cy5 and autocorrelation a) in the presence of 200 uM TCEP and b) absence of TCEP. The autocorrelation function (ACF) of a) was obtained by summation of 88 individual ACFs while in panel b the fluorescent transient shown was used to calculate the ACF. The fit is shown in red from which revealed off-times of a) 8.7 ± 2.8 ms 75 ± 15 μs, b) 9.5 ± 0.7 ms and 73 ± 18 μs.

**Figure S31:**
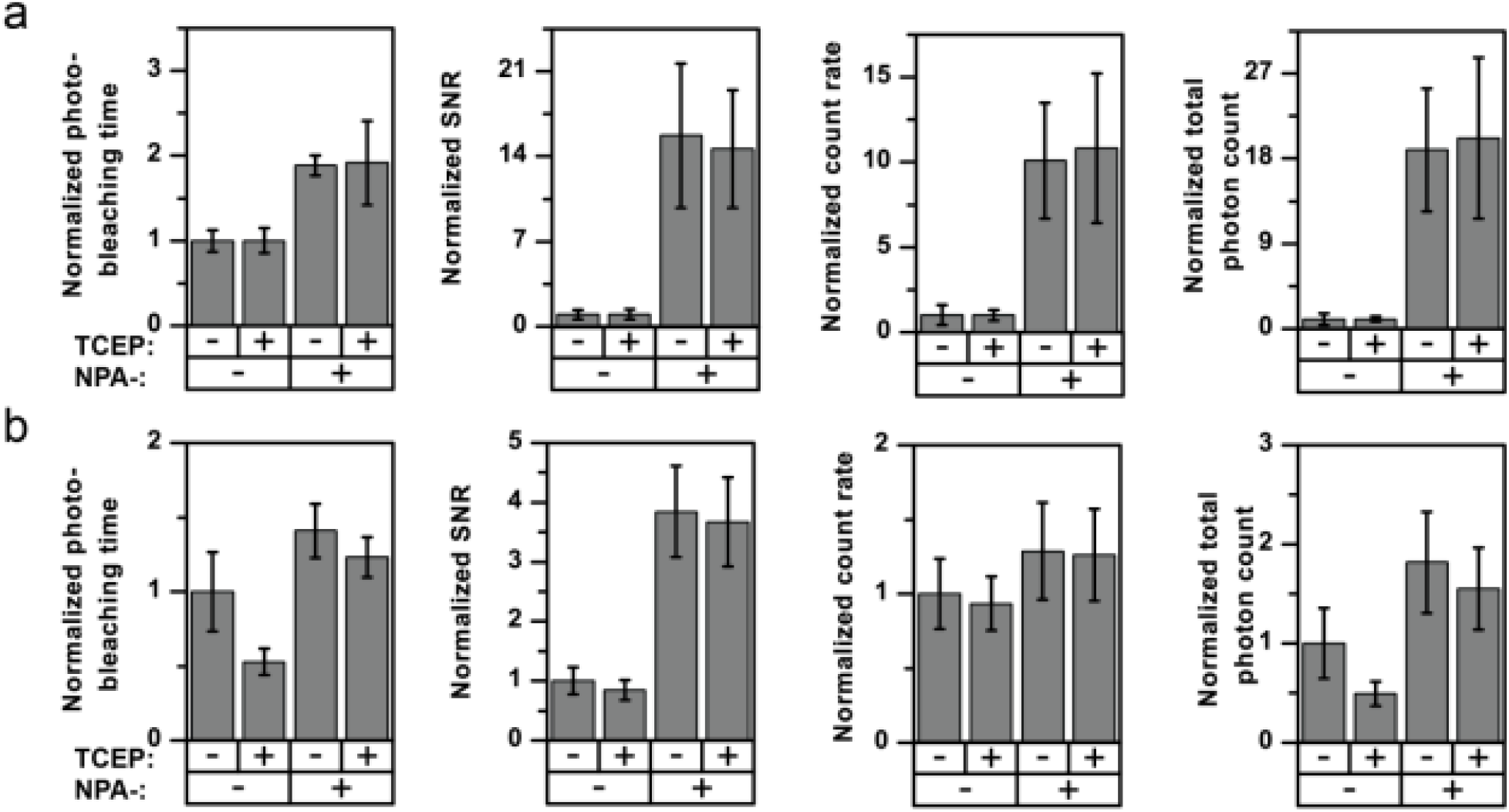
Photophysical characterization of a) ATTO647N and NPA-ATTO647 b) Alexa555 and NPA-Alexa555 in the absence or presence of 200 μM TCEP. All imaging was done under deoxygenated conditions with 400 W cm^-2^ excitation at a) 637 nm or b) 523 nm. Error bars show standard deviation of repeats on 3 different days.

**Figure S32:**
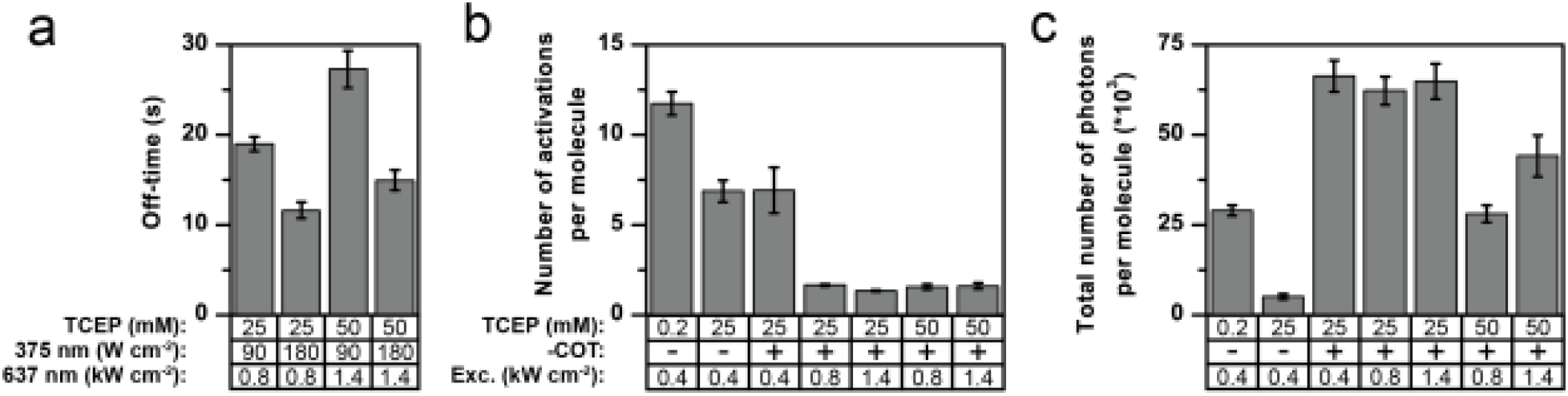
Additional parameters for COT-Cy5 photoactivation using TCEP as photoswitching agent. a) Mean off times between successive activations as a function of 375 nm activation laser power. b) Mean number of activation events per molecule c) Total number of photons detected per activated molecule.

**Figure S33:**
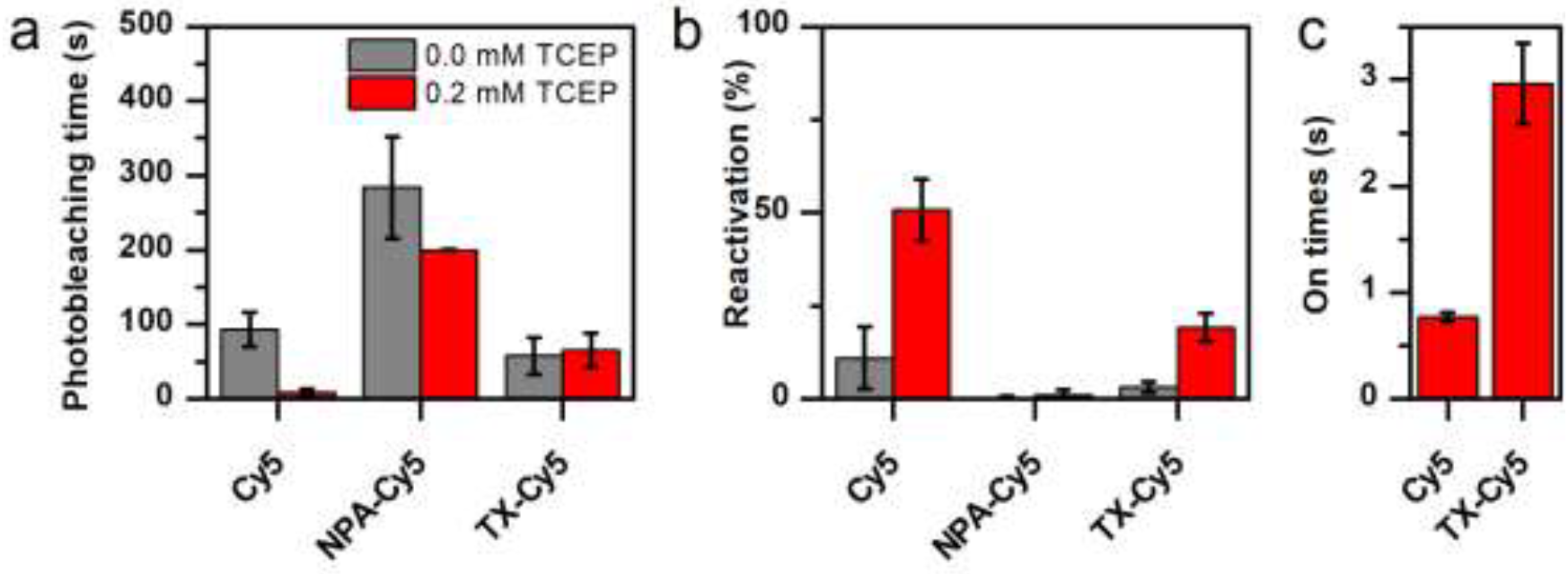
Photoactivation parameters of NPA-Cy5 and TX-Cy5 compared to parent fluorophore Cy5 in the presence and absence of 0.2 mM TCEP, showing (a) Apparent photobleaching time, (b) percentage of fluorophores activated, and (c) on-times of conditions with significant activation. Error bars are standard deviation of repeats (a, b) or SEM (c).

**Figure S34:**
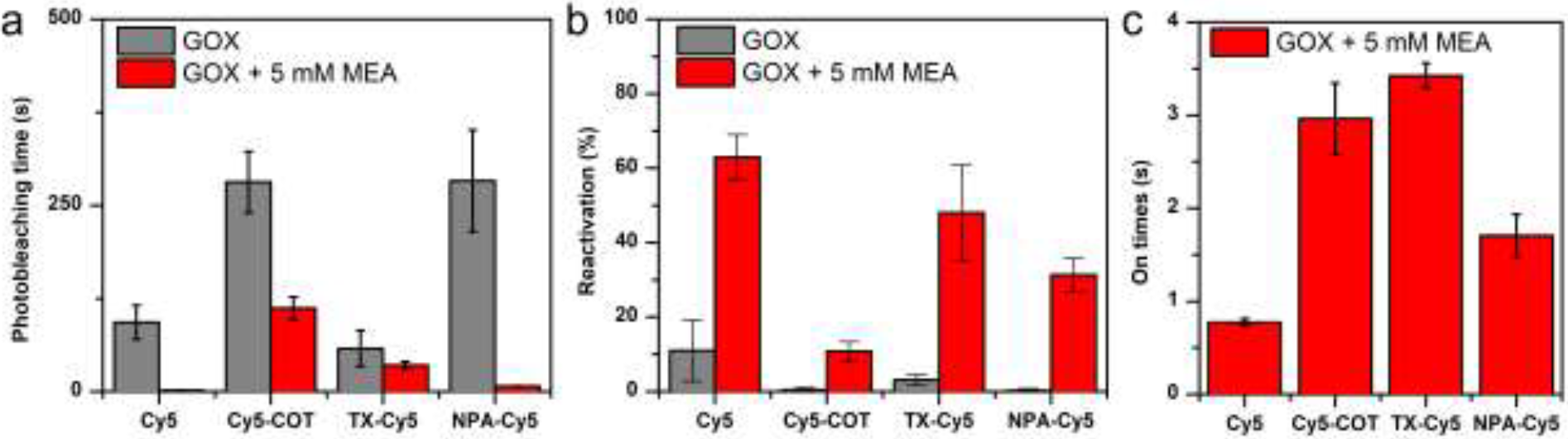
Photoactivation parameters of Cy5-COT, NPA-Cy5 and TX-Cy5 compared to parent fluorophore Cy5 in the presence and absence of 5 mM MEA (cysteamine), showing (a) Apparent photobleaching time, (b) percentage of fluorophores activated, and (c) on-times of conditions with significant activation. Error bars are standard deviation of repeats (a, b) or SEM (c).

**Figure S35:**
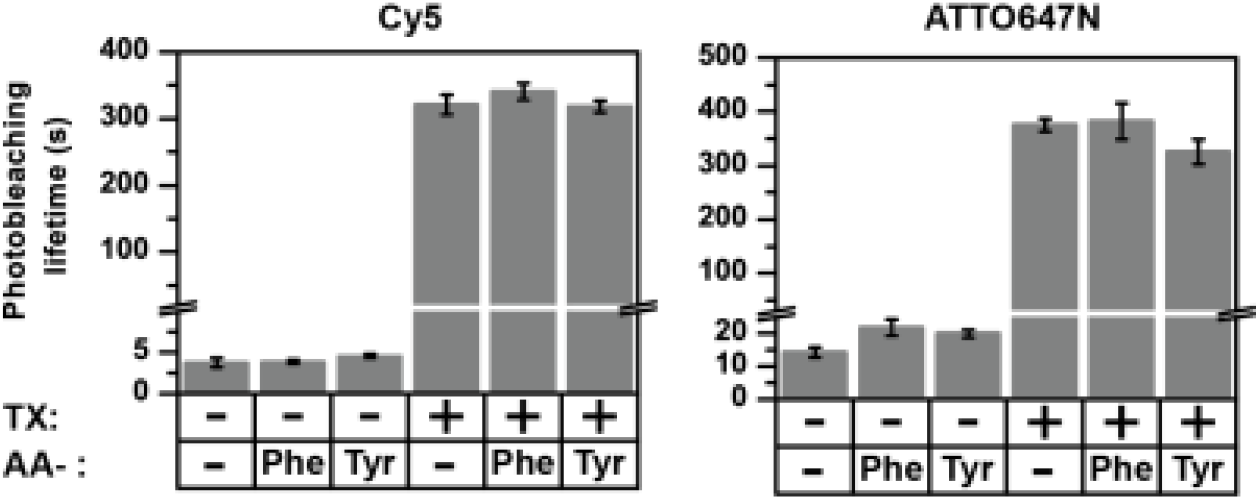
Photobleaching lifetimes of (a) Cy5 and (b) ATTO647N, showing the influence of covalently coupled phenylalanine and tyrosine in the absence and presence of 2 mM TX. All measurements were done in the absence of oxygen with 637 nm excitation (800 W cm^-2^). Error bars show standard deviation of repeats on 2 different days. Labelled ssDNA samples were prepared as described previously using amino-acid scaffolding.^2^

### 1. Details of chemical synthesis

#### Synthesis of the brominated cyclooctatetraene via a modified literature procedure.^1^

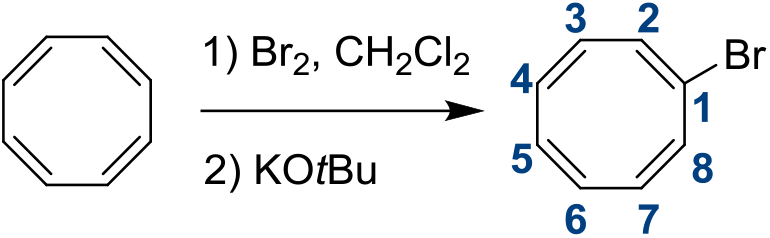

##### 1-Bromocyclooctratetraene (1)

A solution of cyclooctatetraene (1.0 mL, 8.68 mmol) in DCM (10 mL) was cooled to −70 °C under argon atmosphere. At this point, bromine (0.46 mL, 8.95 mmol) in 6 mL DCM was added dropwise to the reaction mixture and stirred for 1.5 h. Then, a solution of KO*t*Bu (2.37 g, 12.2 mmol) in 6 mL THF was slowly added drop by drop to the solution, warmed to −60 °C and stirred for additionally 4 h. The reaction mixture was warmed to - 10 °C, poured into ice water and the aqueous layer was extracted with cooled diethyl ether (3 ×5 mL). The combined organic phases were dried over MgSO_4_, filtered and concentrated to obtain a brownish oil without purification. Yield 1.24 g, (6.77 mmol, 78%)

**TLC:** DCM/MeOH 95:5, R_f_ **(1)** = 0.98.

**^1^H NMR** (400 MHz, CDCl_3_) δ = 6.22 (s, 1H, H-8), 5.98 – 5.74 (m, 5H, H-3, H-4, H-5, H-6, H-7), 5.67-5.60 (m, 1H, H-2) ppm.

**^13^C-NMR** (400 MHz, CDCl3): δ 133.4 1(C), 133.3 (1C), 133.0 (1C), 132.6 (1C), 132.3 (1C), 131.1 (2C), 121.6 (1C) ppm.

**HRMS** (M+H+) calculated for C_8_H_7_Br 182.9804 g mol^-1^, found 182.9803 g mol^-1^.

#### Synthesis of the alcohol derivative of cyclooctatetraene

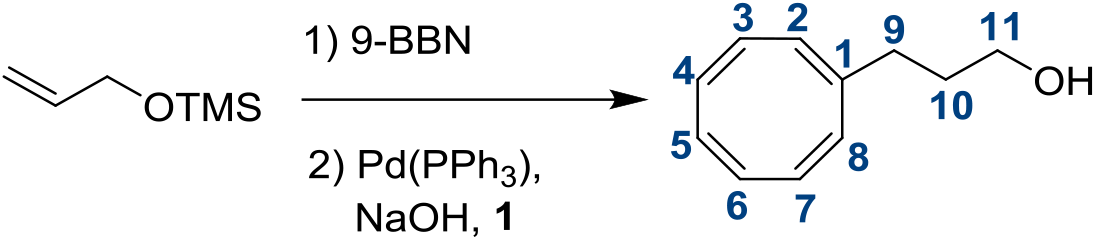

##### Cyclooctatetraenyl-propanol (2)

Allyloxytrimethylsilane (1.0 mL, 5.93 mmol) in 5 mL dry THF and 9-BBN (13 mL, ~ 6.5 mmol) were stirred at 0° C under argon for 2.5 h. Then, Pd(PPh_3_)_4_ (100.3 mg, 0.09 mmol), 5 mL 3 M NaOH and **1** (1.3 g, 7.10 mmol) were added and the reaction mixture was refluxed for 15 h. Afterwards, the mixture was cooled and diluted with hex/EtOAc (1:1), washed with brine and water, dried over MgSO_4_ and concentrated. The crude product was purified by column chromatography (SiO_2_, DCM/MeOH 97:3) to yield a yellowish oil. Yield: 0.639 g (3.94 mmol, 56%).

**TLC:** DCM/MeOH 97:3, R_f_ **(2)** = 0.63.

**^1^H NMR** (400 MHz, CDCl_3_) δ = 5.85 – 5.73 (m, 6H, H-3, H-4, H-5, H-6, H-7, H-8), 5.60 (s, 1H, H-2), 3.70 (tr, *J* = 6.3 Hz, 2H, H-11), 2.14 (tr, *J* = 5.8 Hz, 2H, H-9), 1.68 (quint, *J* = 7.2 Hz, 2H, H-10) ppm.

#### Synthesis of the carboxylic acid derivate of cyclooctatetraene

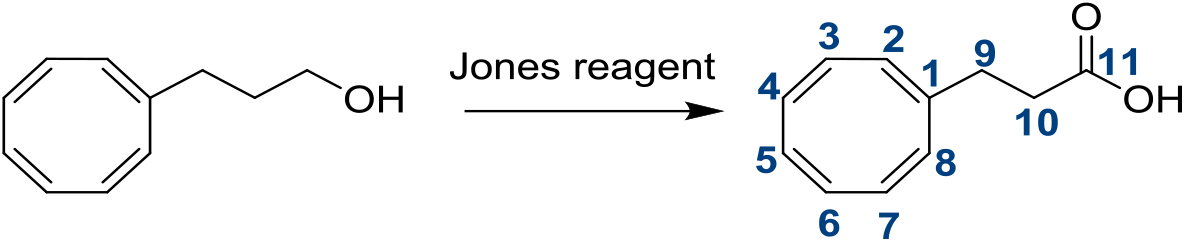

##### Cyclooctatetraenyl-propanoic acid (3)

A solution of **2** (23.6 mg, 0.15 mmol) in 0.5 mL acetone and 100 μL Jones reagent (3 M) were stirred at 0 °C for 1 h. At this point, the reaction was quenched with MeOH and the solution was concentrated. The crude product was dissolved in EtOAc/H_2_O and the water layer was extracted with EtOAc. Then, the combined organic phases were dried over Na_2_SO_4_ and concentrated to receive the crude acid as a yellowish oil. Yield: not determined.

**TLC:** DCM/MeOH 96:4, R_f_ (**3**) = 0.26.

**^1^H NMR** (400 MHz, CDCl_3_) δ = 5.90 – 5.69 (m, 6H, H-3, H-4, H-5, H-6, H-7, H-8), 5.62 (s, 1H, H-2), 2.49 (tr, *J* = 7.4 Hz, 2H, H-10), 2.40 - 2.32 (m, 2H, H-9) ppm.

#### Synthesis of the cyclooctatetraene NHS ester derivate COT-NHS

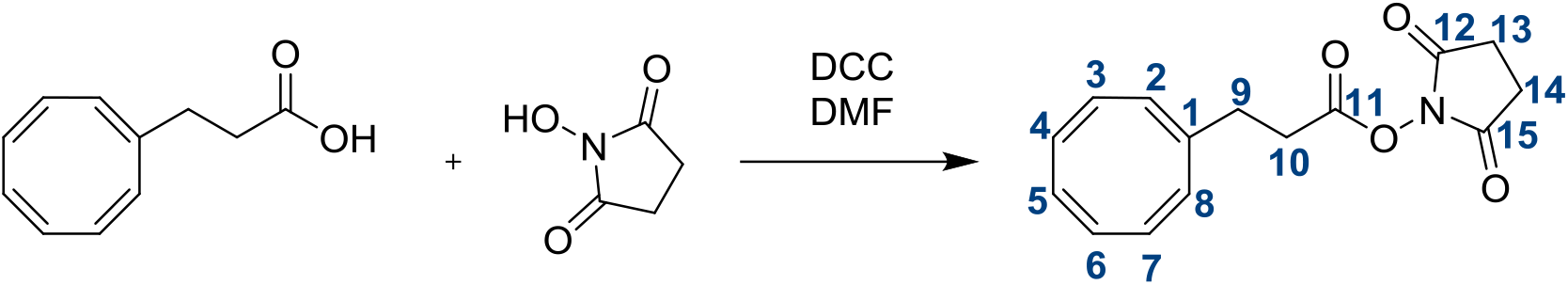

##### COT-NHS (4)

To a solution of **3** (20 mg, 0.11 mmol) and *N*-hydroxysuccinimide (NHS) (32 mg, 0.28 mmol), in 1.5 mL DMF, *N,N´*-dicyclohexyl carbodiimide (DCC) (59 mg, 0.29 mmol) was added at 0 °C. The resulting mixture was stirred for 19 h at rt. Then, the reaction solution was concentrated and the crude product was purified by column chromatography (SiO_2_, DCM/MeOH 19:1) to gain a light yellow oil. Yield: 8.2 mg (0.03 mmol, 26%). **TLC:** DCM/MeOH 19:1, R_f_ **(4)** = 0.95.

**Figure S36:**
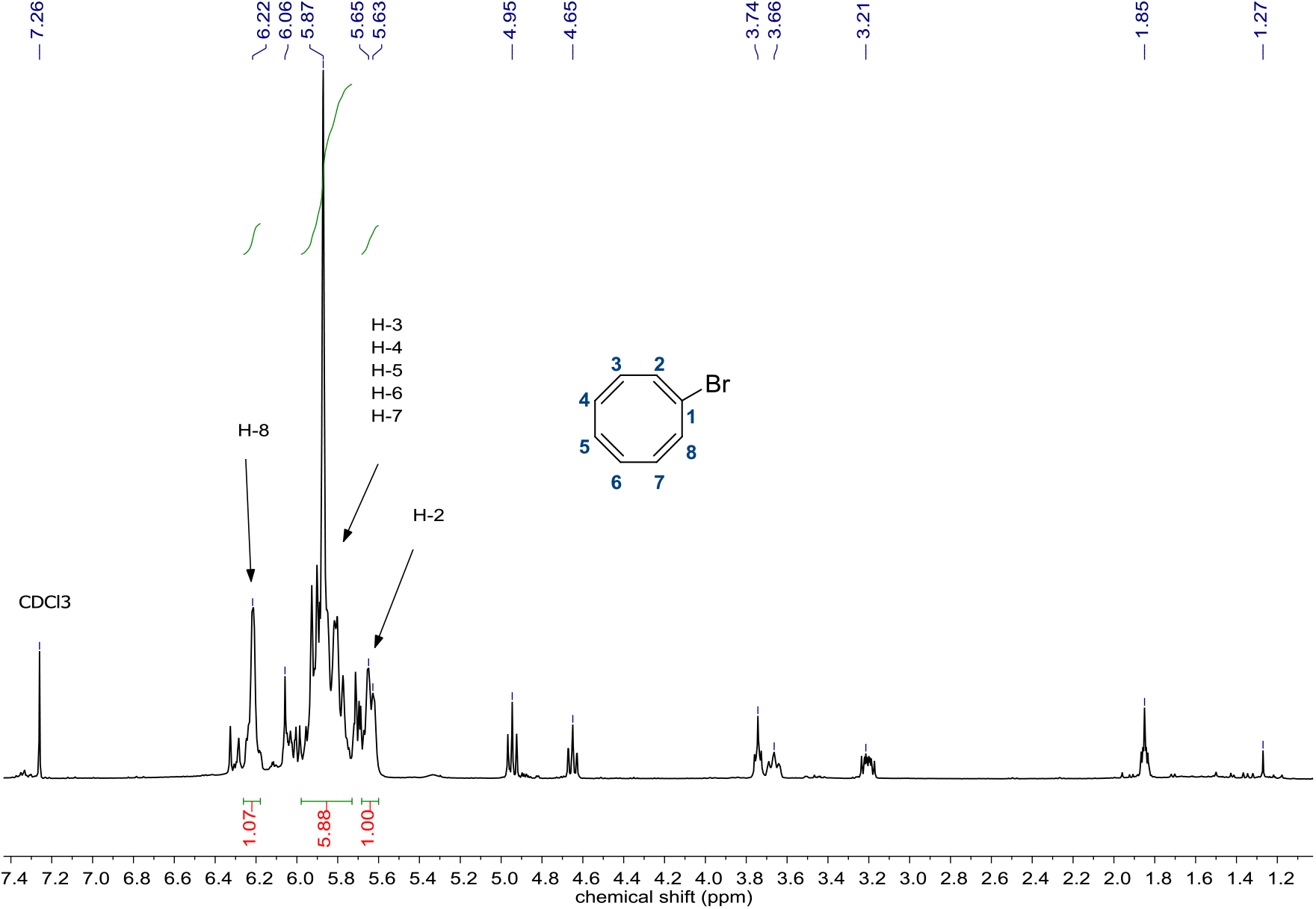
^1^H-NMR spectrum of brominated cylcooctatetraene **1**.

**Figure S37:**
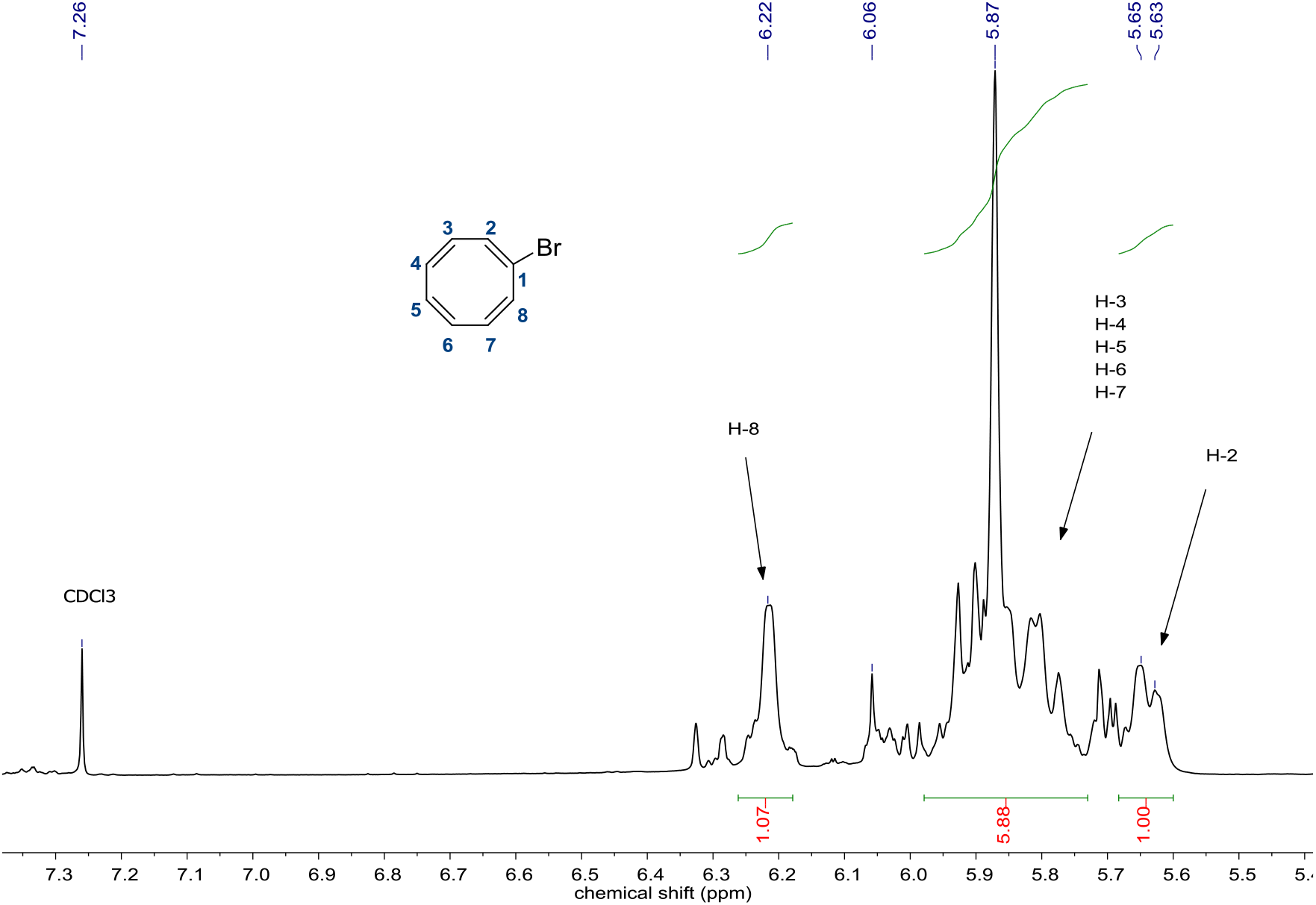
^1^H-NMR spectrum of brominated cylcooctatetraene **1** (7.35-5.42 ppm).

**Figure S38:**
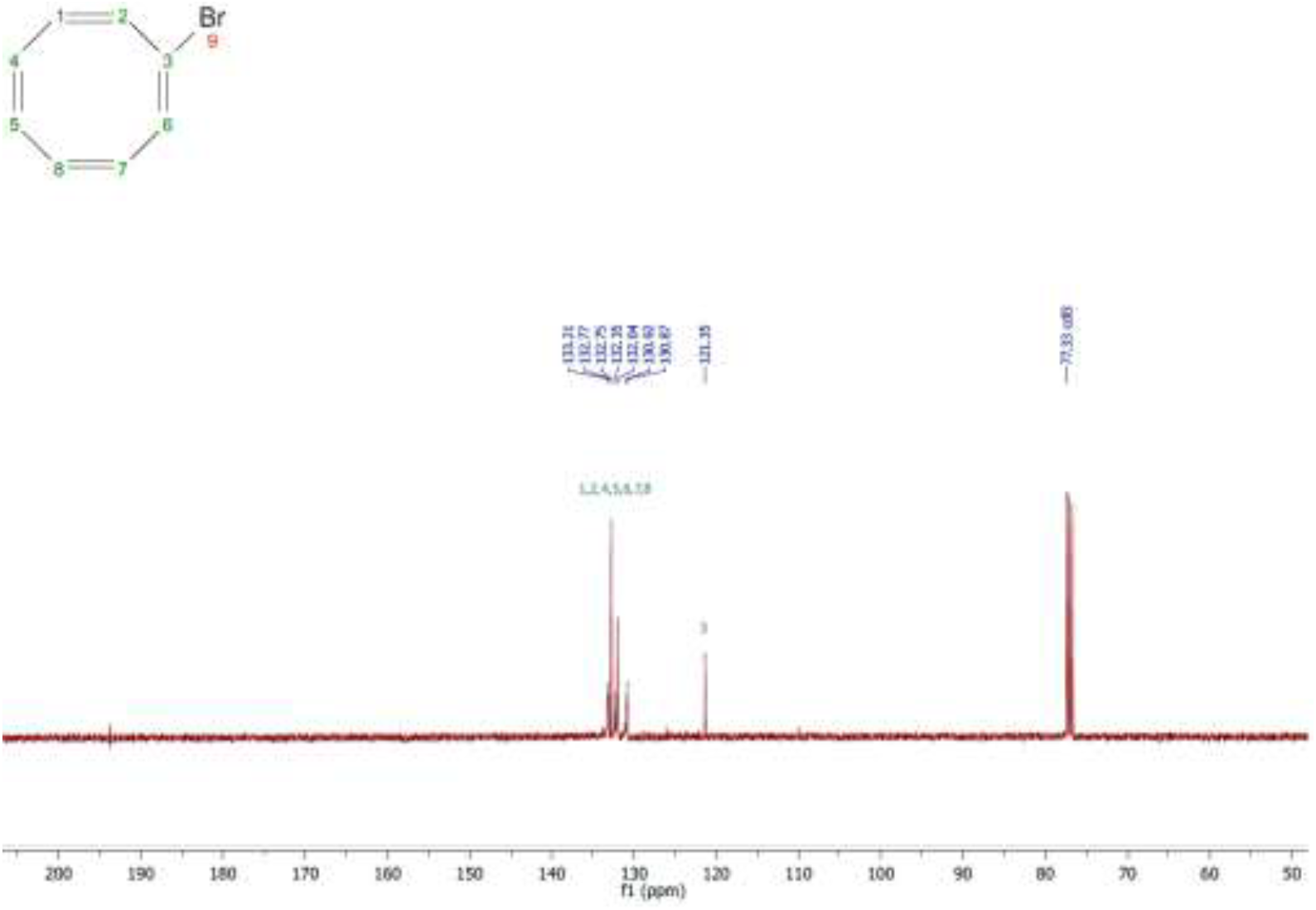
^13^C-NMR spectrum of bominated cylcooctatetraene **1**.

**Figure S39:**
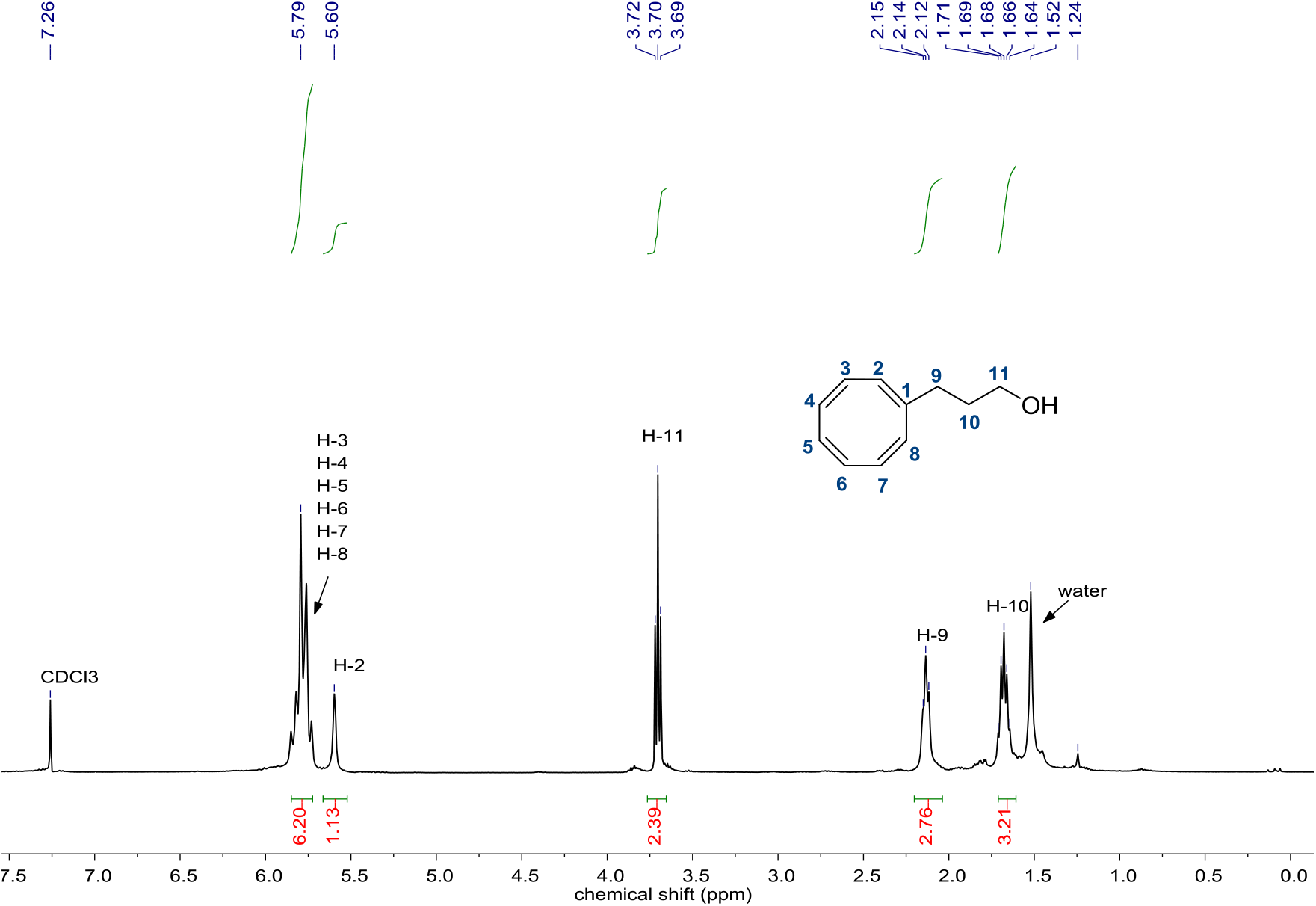
^1^H-NMR spectrum of alcohol **2**.

**Figure S40:**
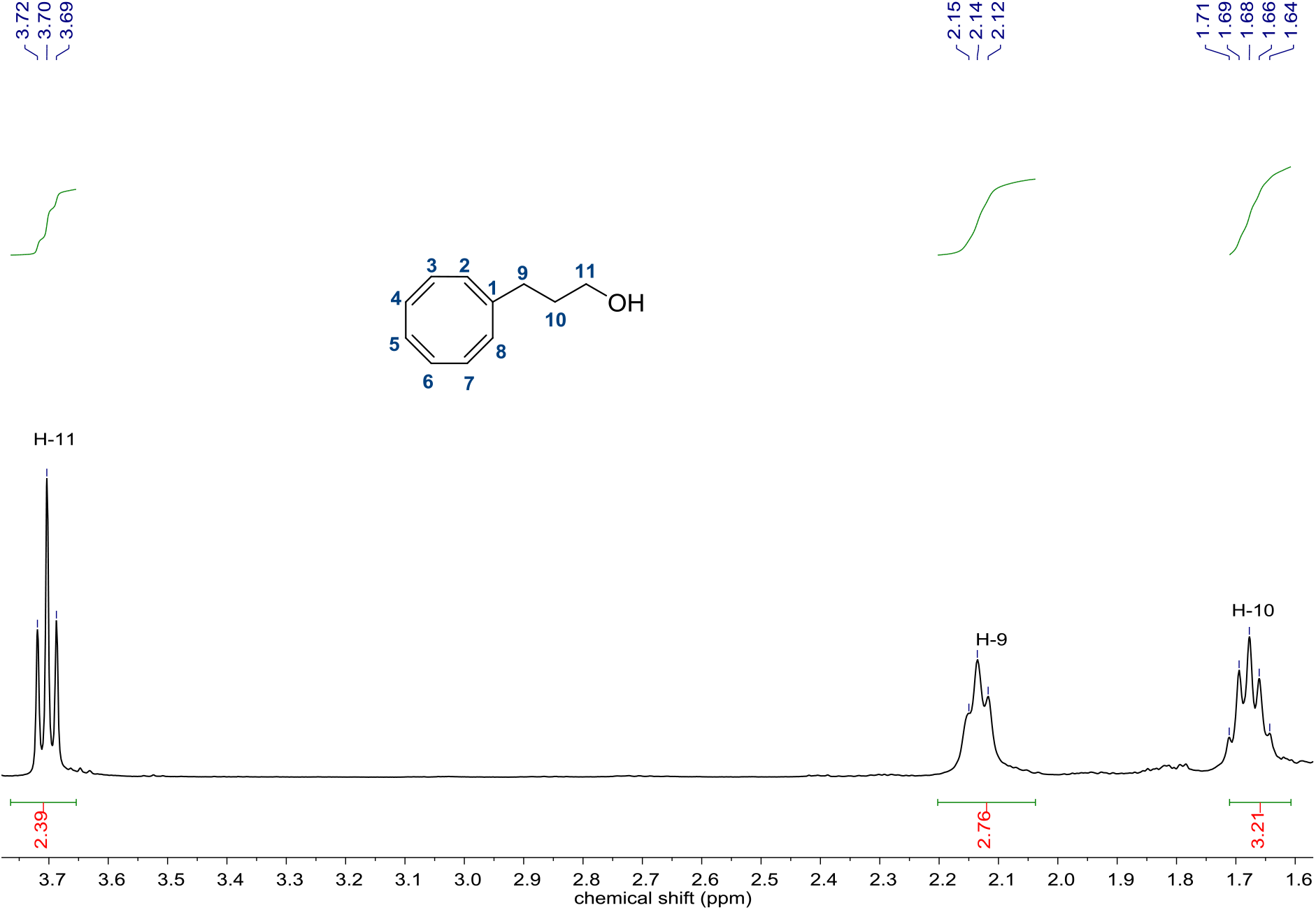
^1^H-NMR spectrum of alcohol **2** (3.78-1.57 ppm).

**Figure S41:**
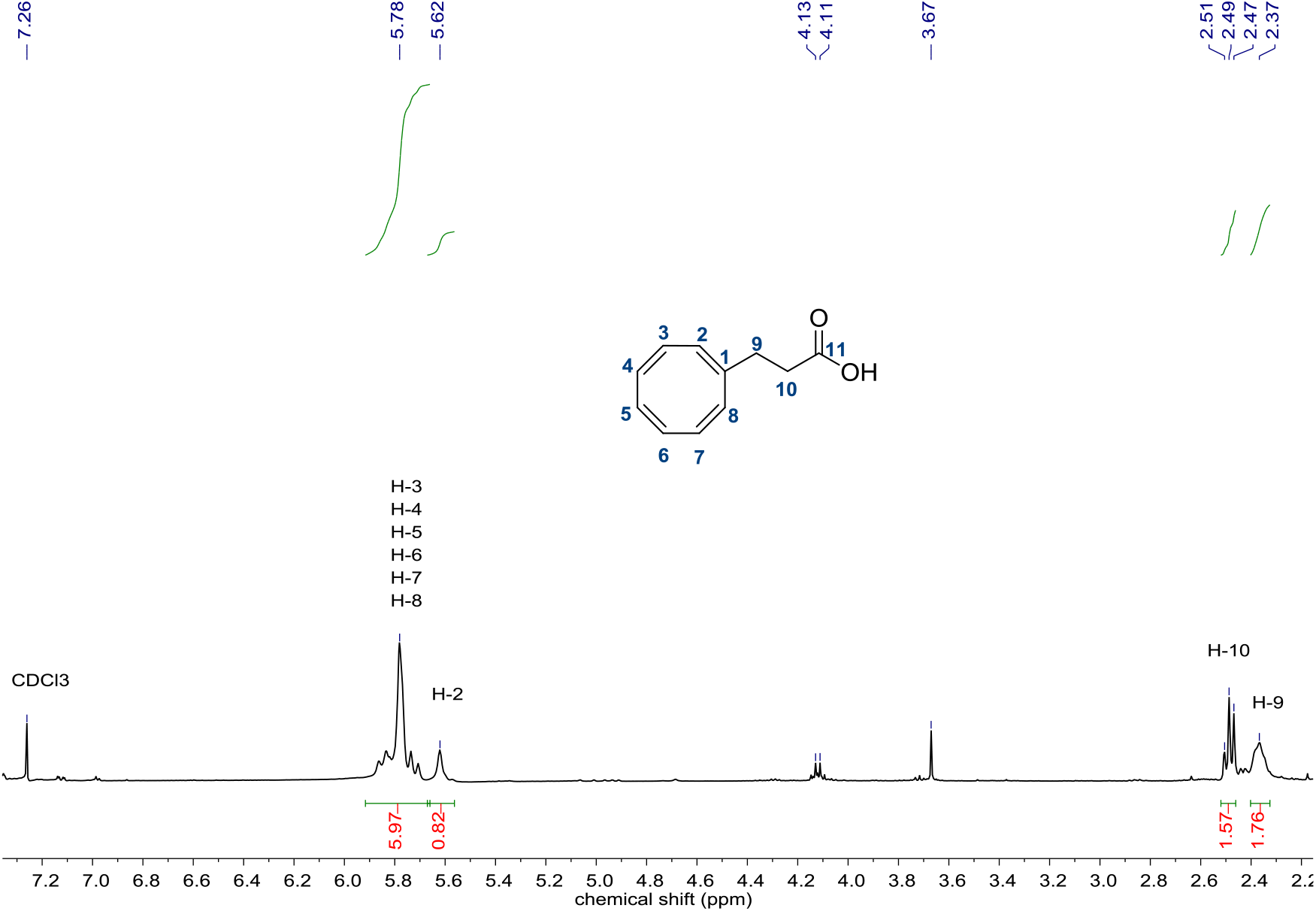
^1^H-NMR spectrum of acid **3**.

**Figure S42:**
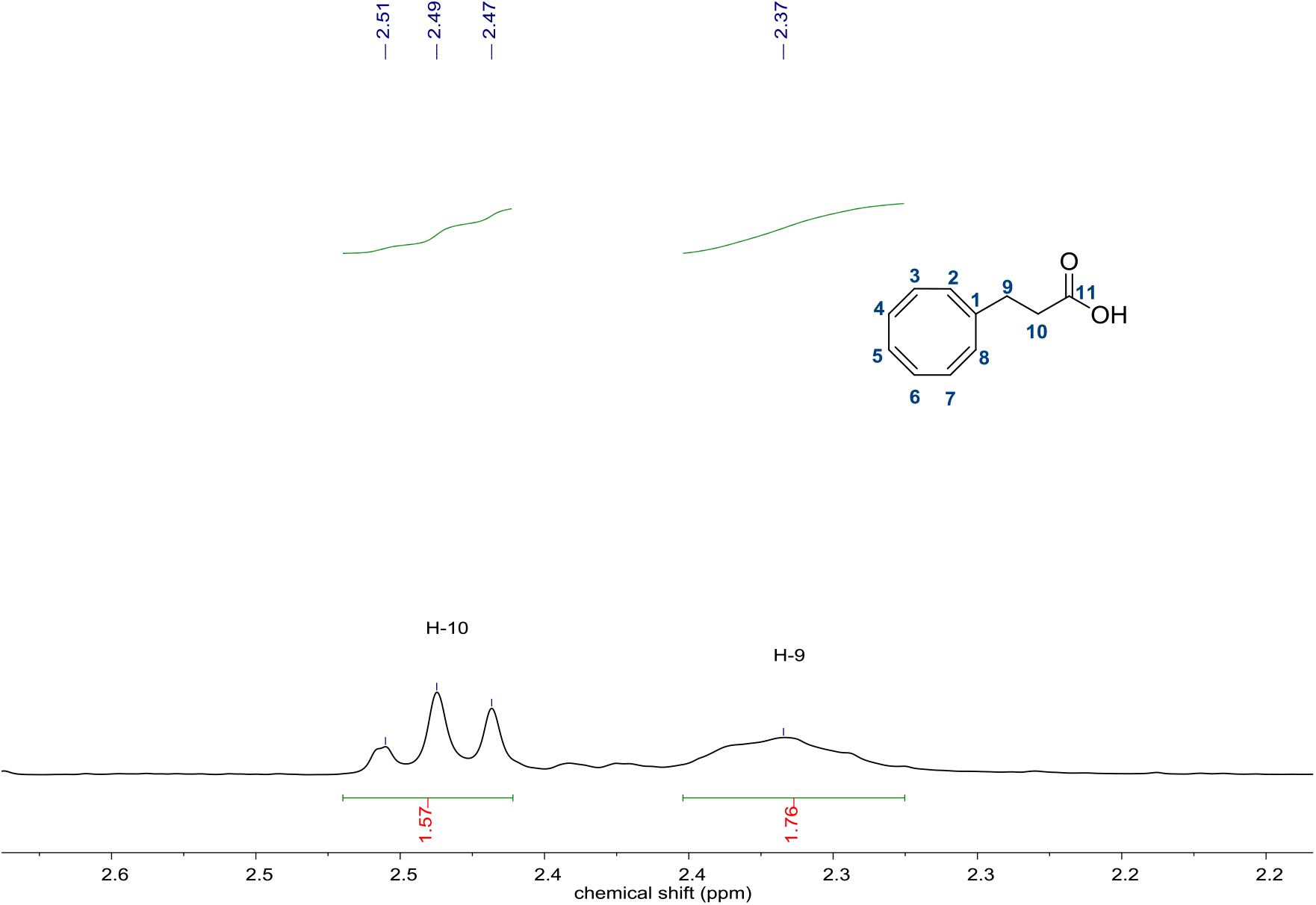
^1^H-NMR spectrum of acid **3** (2.86-2.12 ppm).

**Figure S43:**
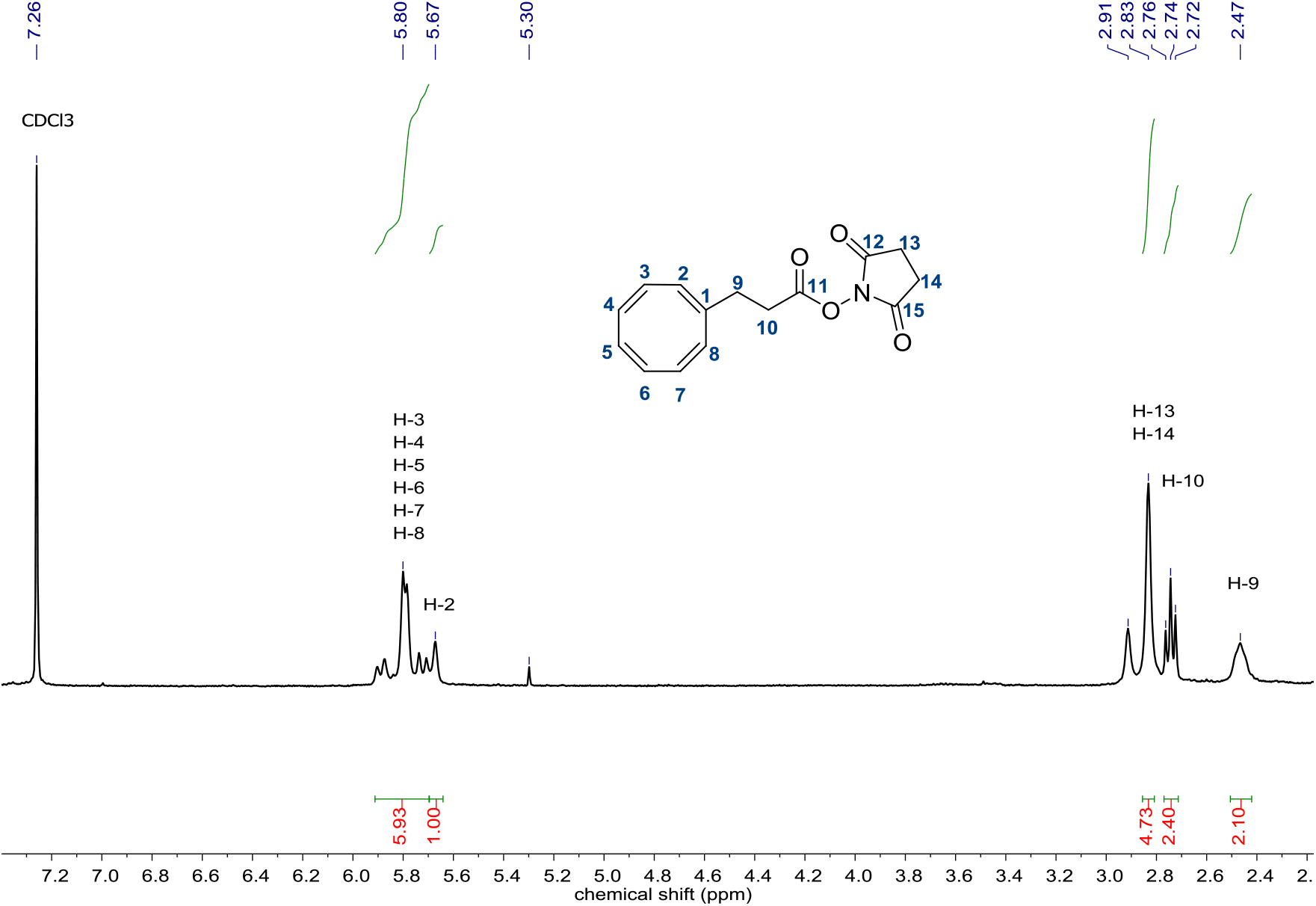
^1^H-NMR spectrum of reactive ester **COT-NHS**.

**Figure S44:**
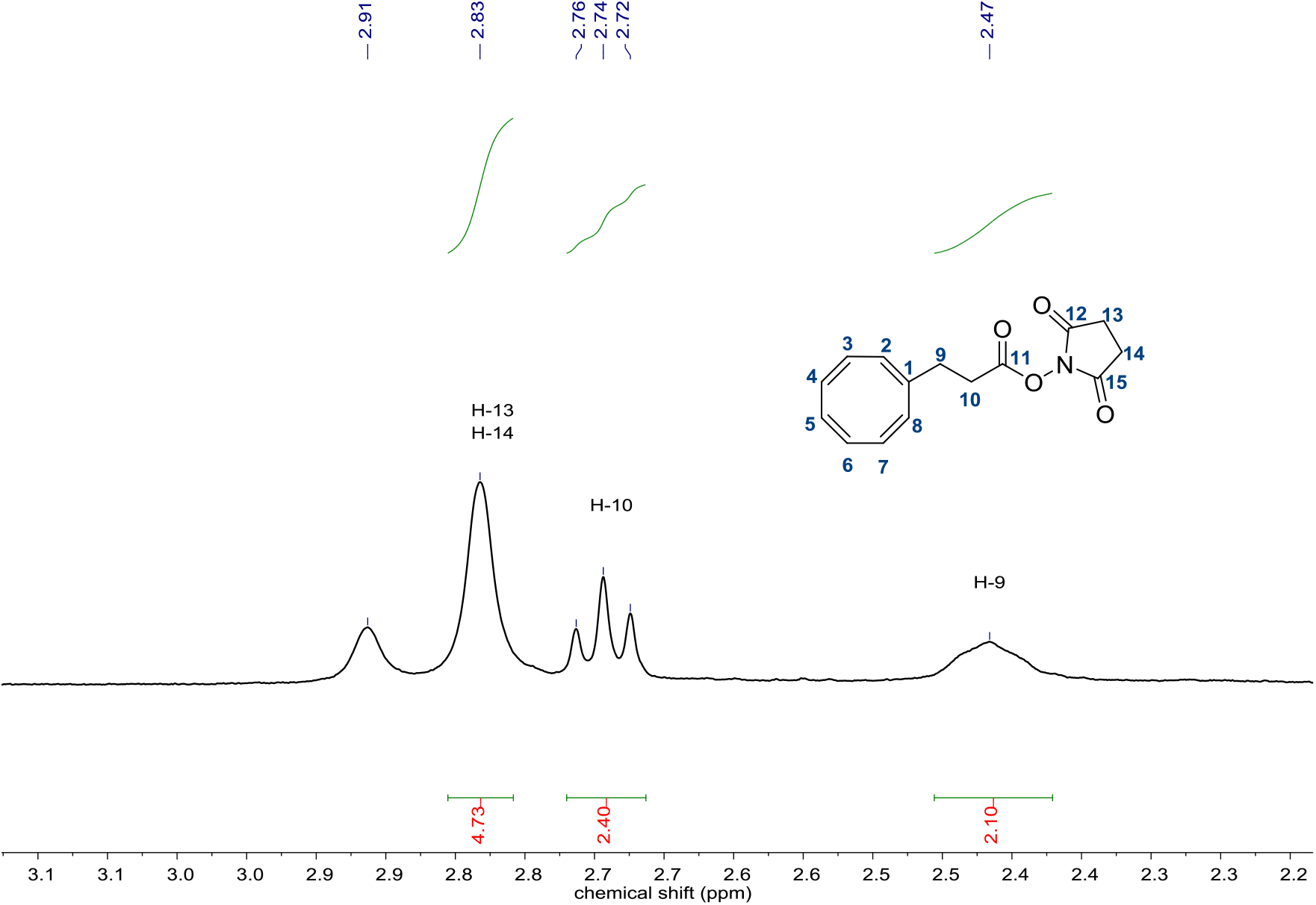
^1^H-NMR spectrum of reactive ester COT-NHS (3.22-2.16 ppm).

**Figure S45:**
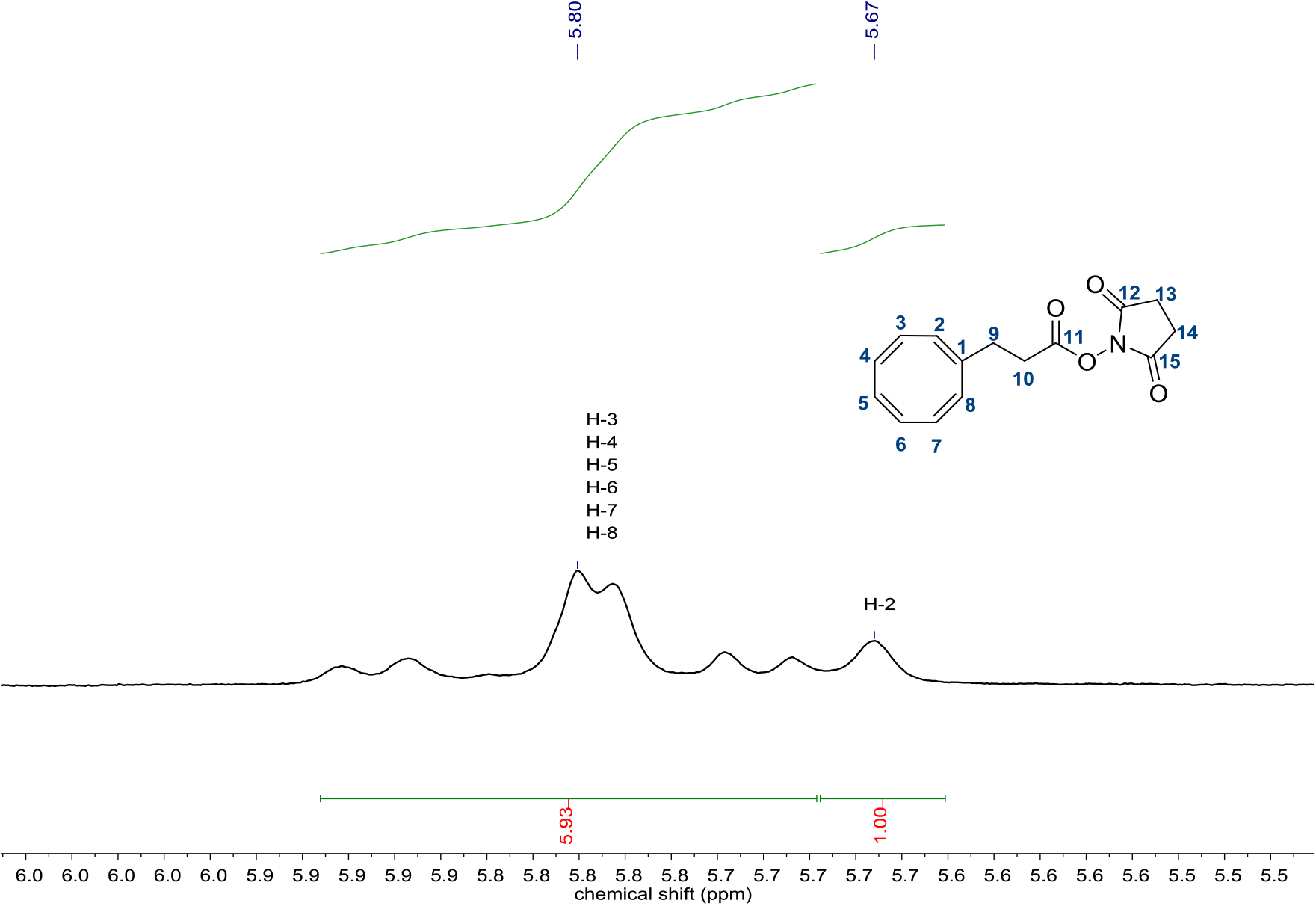
^1^H-NMR spectrum of reactive ester COT-NHS (6.05-5.49 ppm).

#### Synthesis of a DNA-COT conjugate

##### P2 COT

The lyophilized ssDNA-NH_2_ named P2 (Biotin-5’-CGT CCA GAG GAA TCG AAT ATT A-3’-NH_2_) was resuspended in MilliQ water and the concentration was adjusted to 20 μM in 0.2 M NaHCO_3_ buffer (pH 8.35). To this solution, 50 equivalents of **COT-NHS** was added in 100 μL of DMF and the mixture was vortexed thoroughly. After incubation overnight, the oligonucleotide was purified on illustra NAP 5 column (*vide supra*) and isolated by preparative rp-HPLC (gradient 1, *vide supra*) to yield **P2-COT**. The yield was found to be 32% for coupling **COT-NHS** to ssDNA. MS (MALDI-TOF): 7537 g mol^-1^ (found), 7540 g mol^-1^ (calc.)

**Figure S46:**
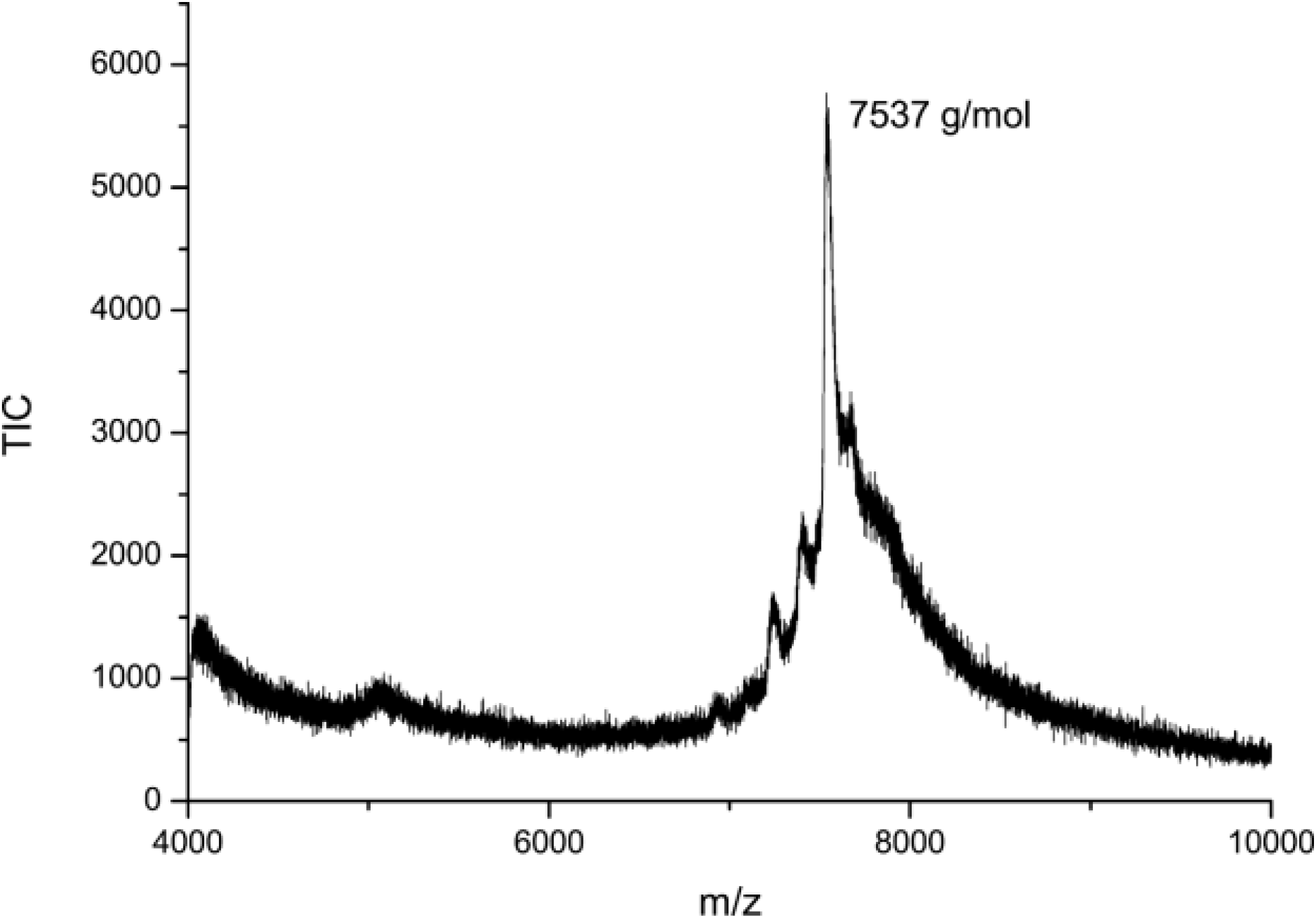
MALDI-TOF mass spectrum of P2-COT.

#### Characterization of DNA-Trp-fluorophore conjugates

**Figure S47:**
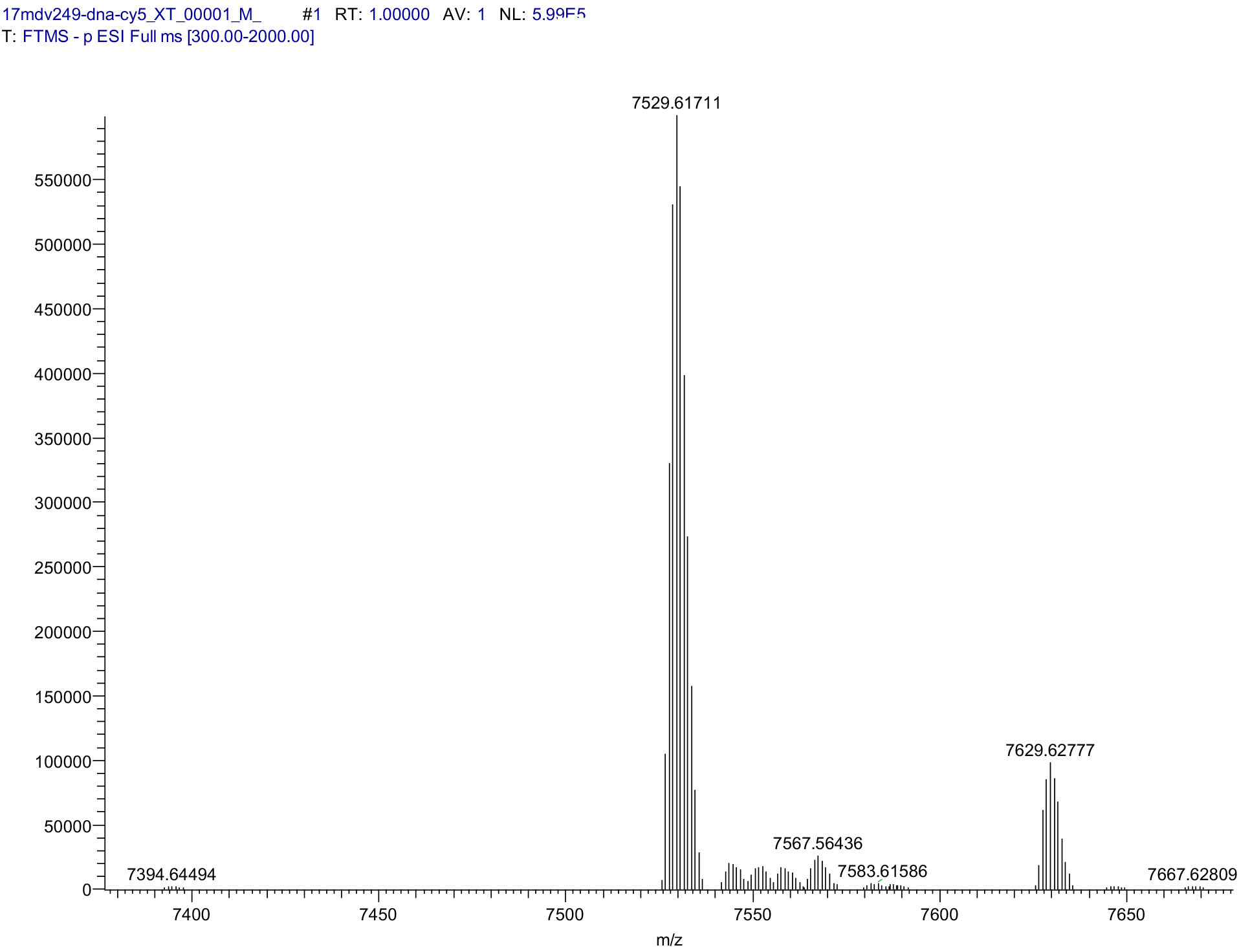
ESI-MS Spectrum of P1-Tryp-Cy5.

**Figure S48:**
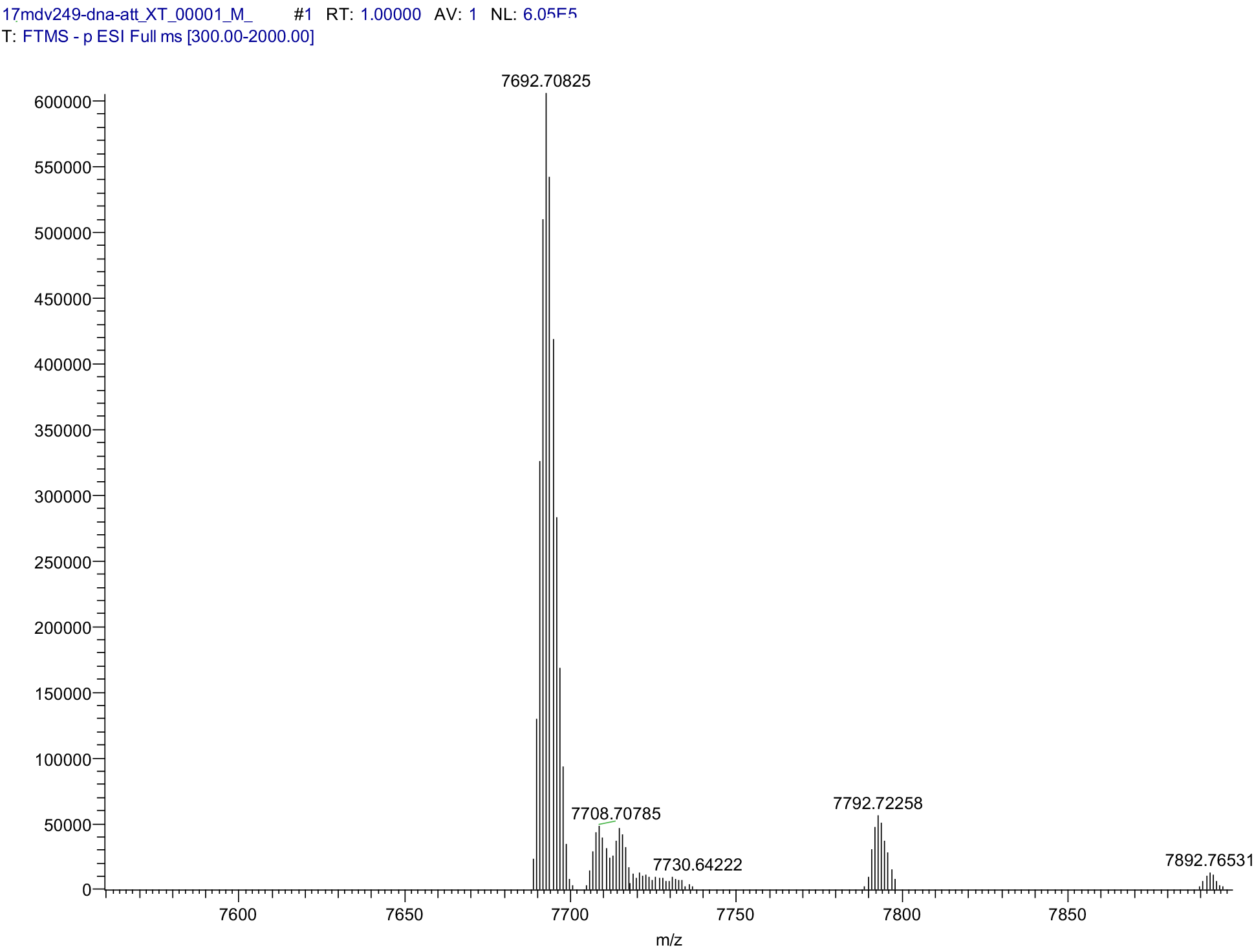
ESI-MS Spectrum of P1-Tryp-ATTO 647N.

